## Supplemental Material for "Single-component multilayered self-assembling protein nanoparticles presenting glycan-trimmed uncleaved prefusion optimized envelope trimers as HIV-1 vaccine candidates"

### fig. S1

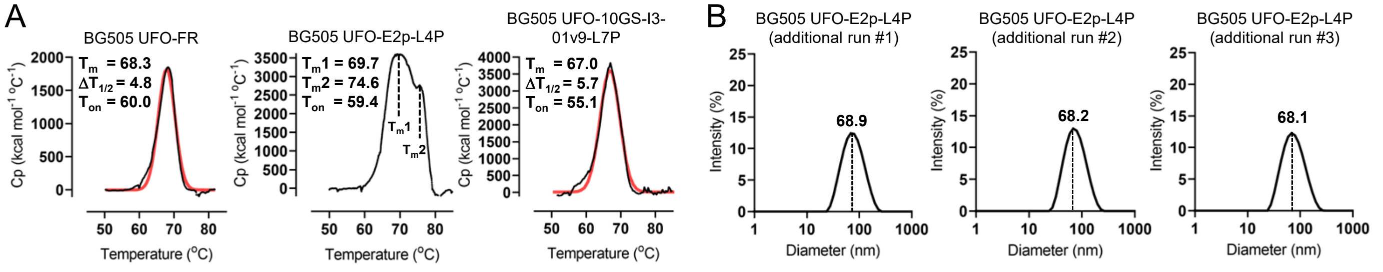


C

**>BG505 UFO-FR**

```
MDAMKRGLCCVLLLCGAVFVSPSQEIHARFRRGARSRAENLWVTVYYGVPVWKDAETTLFCASDAKAYDTEKHNVWATHACVPTDPNPQEIHLENVTEEFNM
WKNNMVEQMHTDIISLWDQSLKPCVKLTPLCVTLQCTNVTNNITDDMRGELKNCSFNMTTELRDKKQKVYSLFYRLDVVQINENQGNRSNNSNKEYRLINCN
TSAITQACPKVSFEPIPIHYCAPAGFAILKCKDKKFNGTGPCPSVSTVQCTHGIKPVVSTQLLLNGSLAEEEVMIRSENITNNAKNILVQFNTPVQINCTRP
NNNTRKSIRIGPGQAFYATGDIIGDIRQAHCNVSKATWNETLGKVVKQLRKHFGNNTIIRFANSSGGDLEVTTHSFNCGGEFFYCNTSGLFNSTWISNTSVQ
GSNSTGSNDSITLPCRIKQIINMWQRIGQAMYAPPIQGVIRCVSNITGLILTRDGGSTNSTTETFRPGGGDMRDNWRSELYKYKVVKIEPLGVAPTRCKRRV
VGGGGGSGGGGSAVGIGAVFLGFLGAAGSTMGAASMTLTVQARNLLSGNPDWLPDMTVWGIKQLQARVLAVERYLRDQQLLGIWGCSGKLICCTNVPWNSSW
SNRNLSEIWDNMTWLQWDKEISNYTQIIYGLLEESQNQQEKNEQDLLALDASGDIIKLLNEQVNKEMQSSNLYMSMSSWCYTHSLDGAGLFLFDHAAEEYEH
AKKLIIFLNENNVPVQLTSISAPEHKFEGLTQIFQKAYEHEQHISESINNIVDHAIKSKDHATFNFLQWYVAEQHEEEVLFKDILDKIELIGNENHGLYLAD
QYVKGIAKSRKS
```

**>BG505 UFO-E2p-LD4-PADRE (or E2p-L4P)**

```
MDAMKRGLCCVLLLCGAVFVSPSQEIHARFRRGARSRAENLWVTVYYGVPVWKDAETTLFCASDAKAYDTEKHNVWATHACVPTDPNPQEIHLENVTEEFNM
WKNNMVEQMHTDIISLWDQSLKPCVKLTPLCVTLQCTNVTNNITDDMRGELKNCSFNMTTELRDKKQKVYSLFYRLDVVQINENQGNRSNNSNKEYRLINCN
TSAITQACPKVSFEPIPIHYCAPAGFAILKCKDKKFNGTGPCPSVSTVQCTHGIKPVVSTQLLLNGSLAEEEVMIRSENITNNAKNILVQFNTPVQINCTRP
NNNTRKSIRIGPGQAFYATGDIIGDIRQAHCNVSKATWNETLGKVVKQLRKHFGNNTIIRFANSSGGDLEVTTHSFNCGGEFFYCNTSGLFNSTWISNTSVQ
GSNSTGSNDSITLPCRIKQIINMWQRIGQAMYAPPIQGVIRCVSNITGLILTRDGGSTNSTTETFRPGGGDMRDNWRSELYKYKVVKIEPLGVAPTRCKRRV
VGGGGGSGGGGSAVGIGAVFLGFLGAAGSTMGAASMTLTVQARNLLSGNPDWLPDMTVWGIKQLQARVLAVERYLRDQQLLGIWGCSGKLICCTNVPWNSSW
SNRNLSEIWDNMTWLQWDKEISNYTQIIYGLLEESQNQQEKNEQDLLALDASGAAAKPATTEGEFPETREKMSGIRRAIAKAMVHSKHTAPHVTLMDEADVT
KLVAHRKKFKAIAAEKGIKLTFLPYVVKALVSALREYPVLNTAIDDETEEIIQKHYYNIGIAADTDRGLLVPVIKHADRKPIFALAQEINELAEKARDGKLT
PGEMKGASCTITNIGSAGGQWFTPVINHPEVAILGIGRIAEKPIVRDGEIVAAPMLALSLSFDHRMIDGATAQKALNHIKRLLSDPELLLMGGGGSFSEEQK
KALDLAFYFDRRLTPEWRRYLSQRLGLNEEQIERWFRRKEQQIGWSHPQFEKGSAKFVAAWTLKAAA
```

**>BG505 UFO-10GS-I3-01v9-LD7-PADRE (or I3-01v9-L7P)**

```
MDAMKRGLCCVLLLCGAVFVSPSQEIHARFRRGARSRAENLWVTVYYGVPVWKDAETTLFCASDAKAYDTEKHNVWATHACVPTDPNPQEIHLENVTEEFNM
WKNNMVEQMHTDIISLWDQSLKPCVKLTPLCVTLQCTNVTNNITDDMRGELKNCSFNMTTELRDKKQKVYSLFYRLDVVQINENQGNRSNNSNKEYRLINCN
TSAITQACPKVSFEPIPIHYCAPAGFAILKCKDKKFNGTGPCPSVSTVQCTHGIKPVVSTQLLLNGSLAEEEVMIRSENITNNAKNILVQFNTPVQINCTRP
NNNTRKSIRIGPGQAFYATGDIIGDIRQAHCNVSKATWNETLGKVVKQLRKHFGNNTIIRFANSSGGDLEVTTHSFNCGGEFFYCNTSGLFNSTWISNTSVQ
GSNSTGSNDSITLPCRIKQIINMWQRIGQAMYAPPIQGVIRCVSNITGLILTRDGGSTNSTTETFRPGGGDMRDNWRSELYKYKVVKIEPLGVAPTRCKRRV
VGGGGGSGGGGSAVGIGAVFLGFLGAAGSTMGAASMTLTVQARNLLSGNPDWLPDMTVWGIKQLQARVLAVERYLRDQQLLGIWGCSGKLICCTNVPWNSSW
SNRNLSEIWDNMTWLQWDKEISNYTQIIYGLLEESQNQQEKNEQDLLALDASGGGGSGGGGSMKMEELFKKHKIVAVLRANSVEEAKMKALAVFVGGVHLIE
ITFTVPDADTVIKELSFLKELGAIIGAGTVTSVEQCRKAVESGAEFIVSPHLDEEISQFCKEKGVFYMPGVMTPTELVKAMKLGHTILKLFPGEVVGPQFVK
AMKGPFPNVKFVPTGGVNLDNVCEWFKAGVLAVGVGSALVKGTIAEVAAKAAAFVEKIRGCTEGGGGSSPAVDIGDRLDELEKALEALSAEDGHDDVGQRLE
SLLRRWNSRRADGSAKFVAAWTLKAAA
```

**fig S1. Design and in vitro characterization of BG505 UFO trimer-presenting SApNPs.** (**A**) Additional thermostability data for BG505 UFO trimer-presenting FR (left), E2p-L4P, and I3-01v9-L7P NPs, with $T_m$ ($T_{m1}$ and $T_{m2}$ for E2p), $\Delta T1/2$, and $T_{on}$ measured by differential scanning calorimetry (DSC) and labeled on the plots. For FR and I3-01v9 NPs, the raw and Gaussian-fitted DSC data are shown in black and red lines, respectively. (**B**) Additional particle size data for BG505 UFO trimer-presenting E2p-L4P NP measured by dynamic light scattering using a Zetasizer. The average particle size is labeled on the plots. (**C**) Construct sequences of BG505 UFO trimer-presenting FR (left), E2p-L4P, and I3-01v9-L7P NPs, with the gene fragments of leader sequence, restriction site, BG505 UFO Env, flexible linker, NP-forming subunit, locking domain (LD), and PADRE highlighted in yellow, green, gray, light magenta, cyan, olive green, and red shades, respectively.

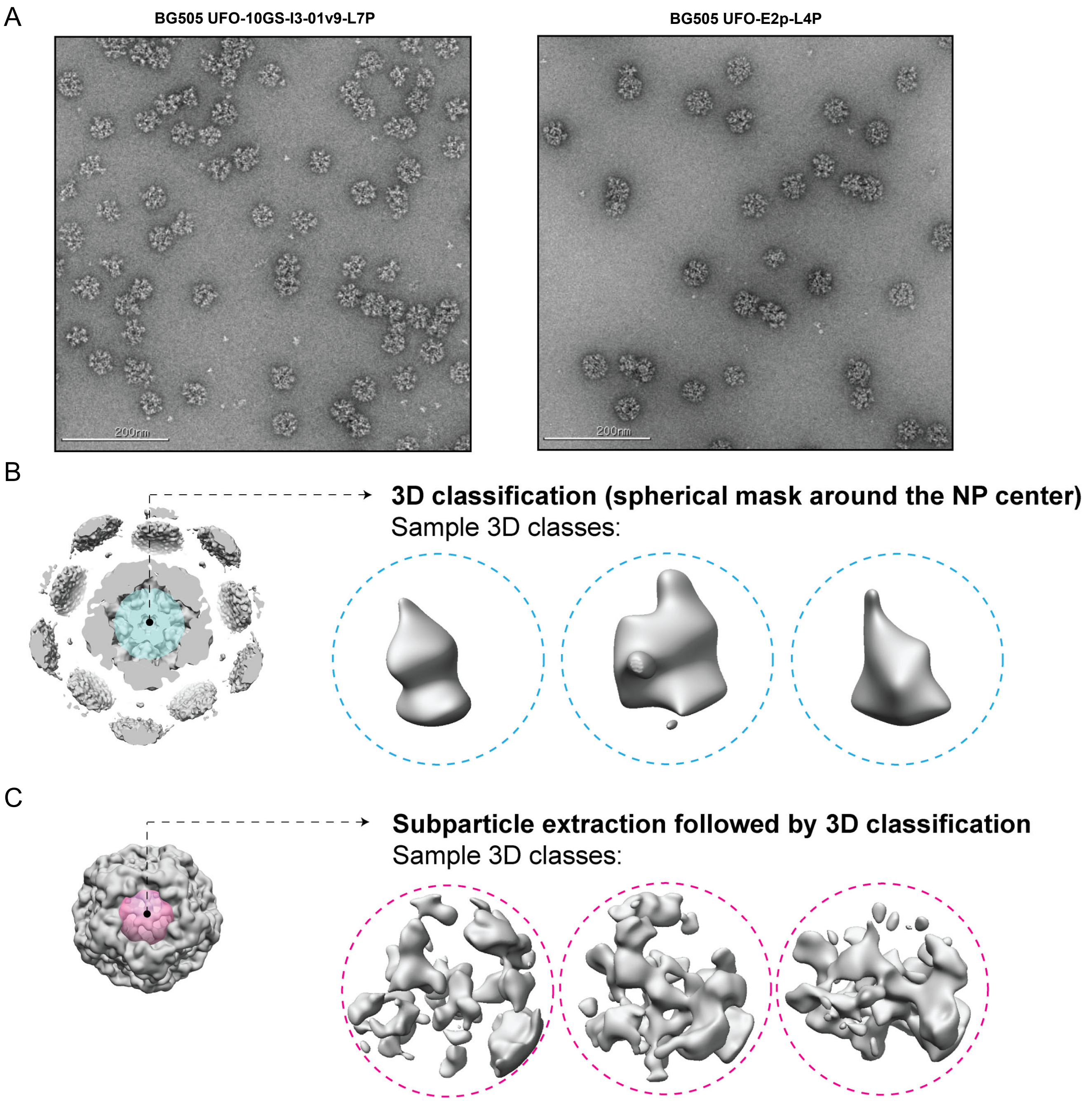


**fig S2. Cryo-EM analysis of BG505 UFO-E2p-L4P and UFO-10GS-I3-01v9-L7PSApNPs.** (**A**) Negative stain EM of the two multilayered SApNP immunogens to validate their structural integrity. For BG505 UFO-E2p-L4P SApNP, cryo-EM data are shown for (**B**) 3D classification (spherical mask around the NP center) and (**C**) Subparticle extraction followed by 3D classification.

fig. S3

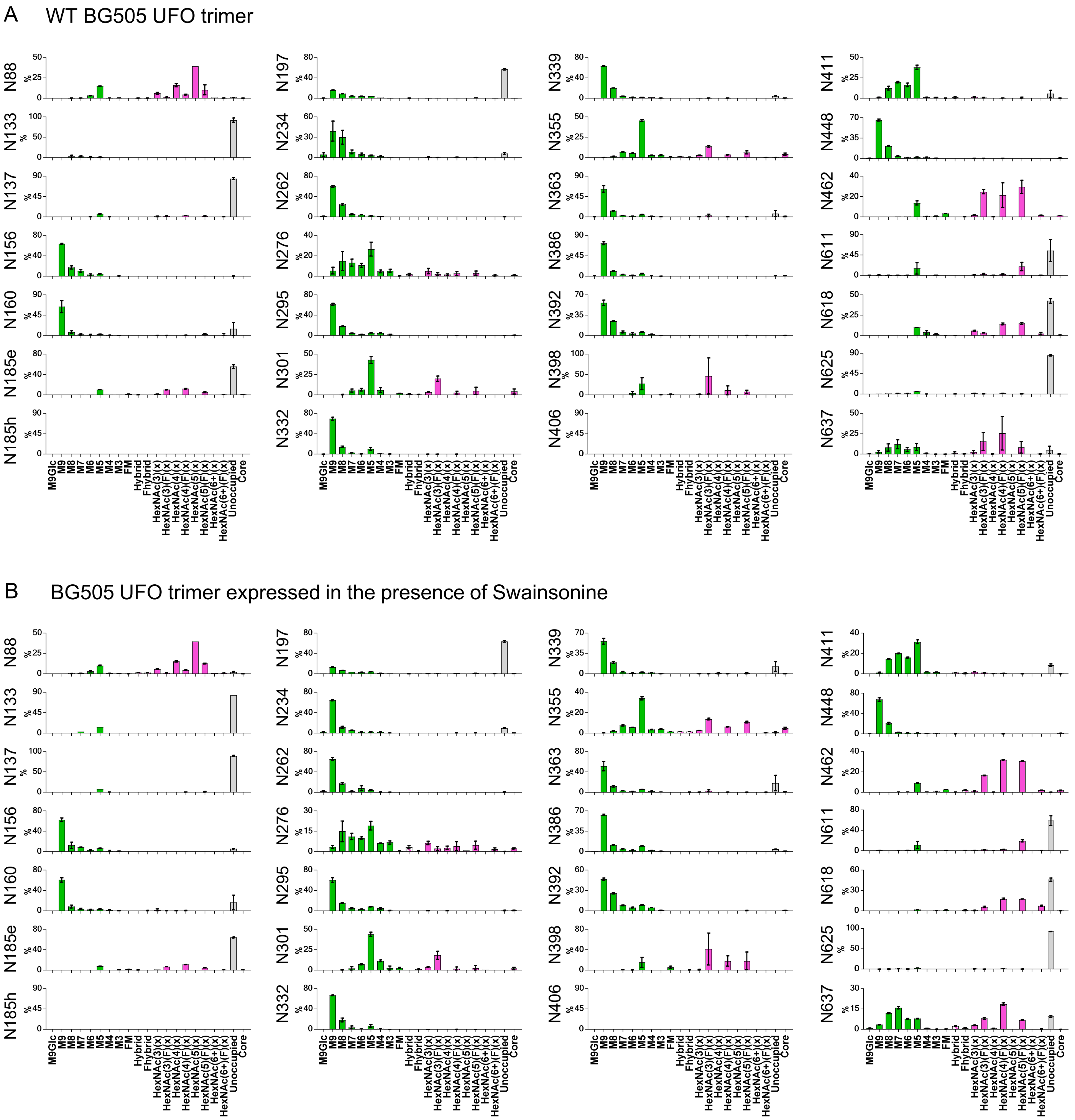
A
WT BG505 UFO trimer
N88
N133
N137
N156
N160
N185e
N185h
N197
N234
N262
N276
N295
N301
N332
N339
N355
N363
N386
N392
N398
N406
N411
N448
N462
N611
N618
N625
N637
%
M9Glc
M9
M8
M7
M6
M5
M4
M3
FM
Hybrid
Fhybrid
HexNAc(3)(x)
HexNAc(3)(F)(x)
HexNAc(4)(x)
HexNAc(4)(F)(x)
HexNAc(5)(x)
HexNAc(5)(F)(x)
HexNAc(6+)(x)
HexNAc(6+)(F)(x)
Unoccupied
Core
B
BG505 UFO trimer expressed in the presence of Swainsonine

C BG505 UFO trimer expressed in the presence of kifunensine

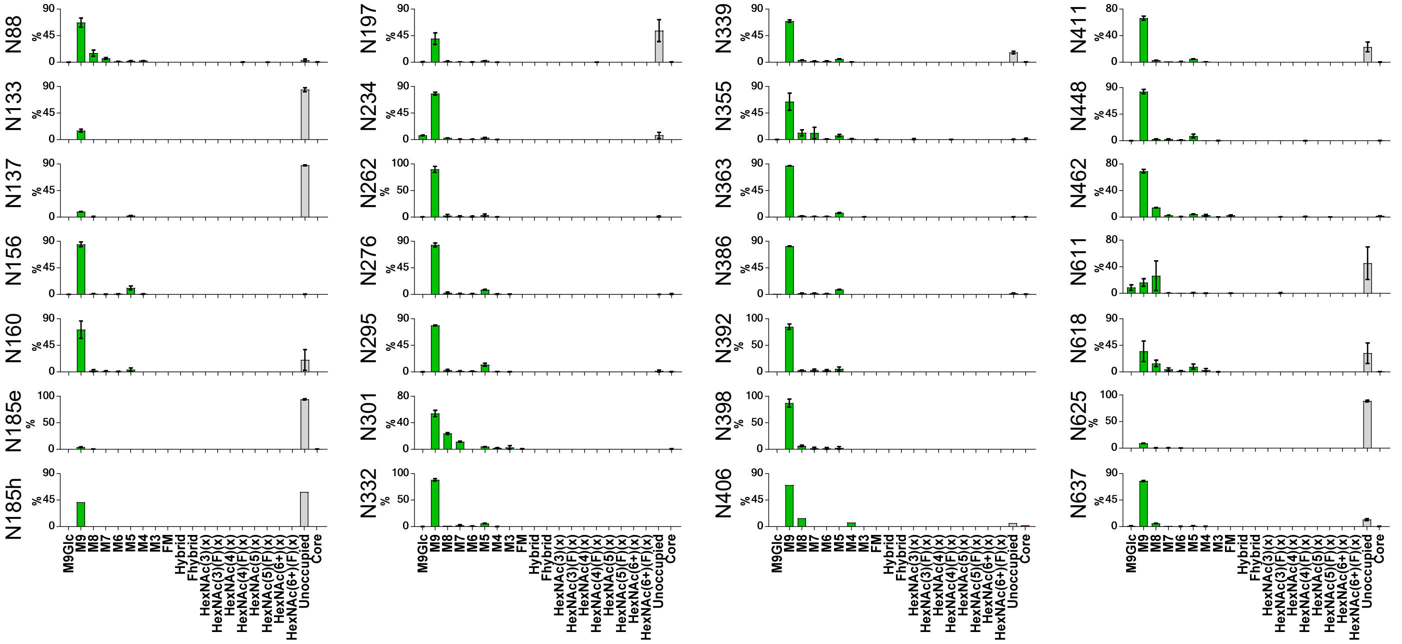


D WT BG505 UFO-FR SApNP

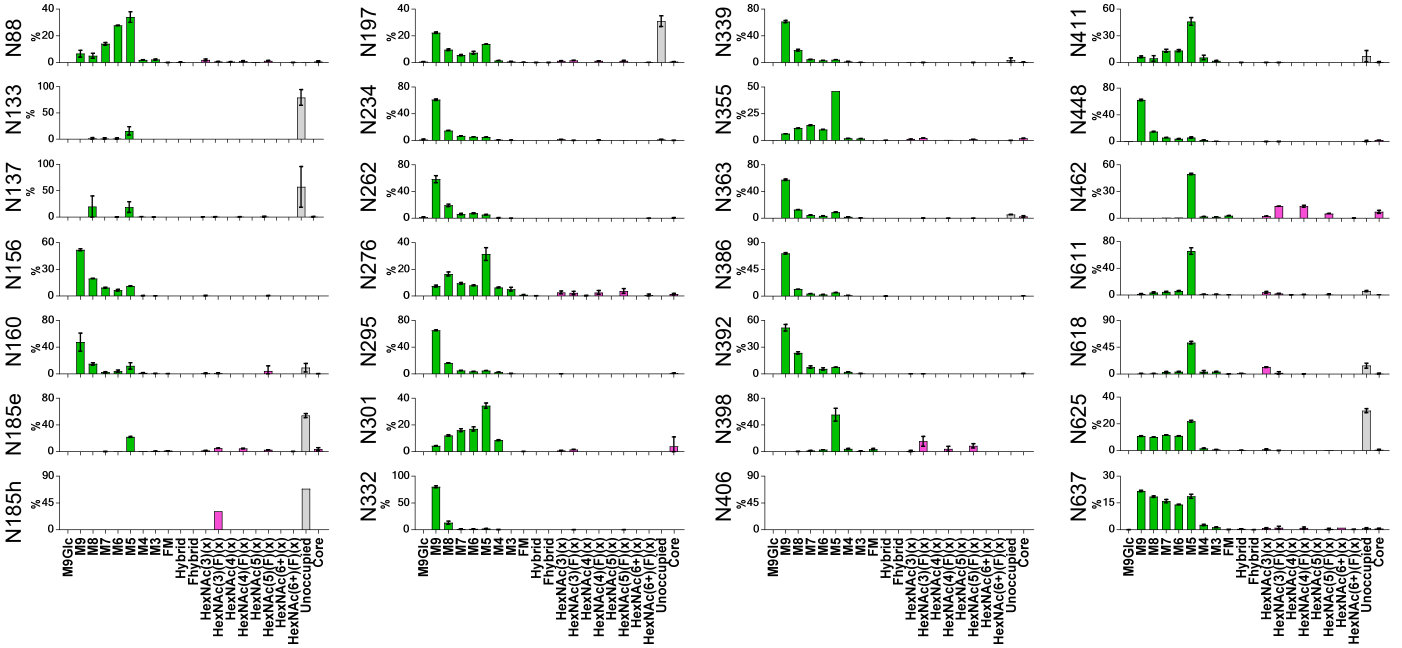

E WT BG505 UFO-E2p-L4P SApNP

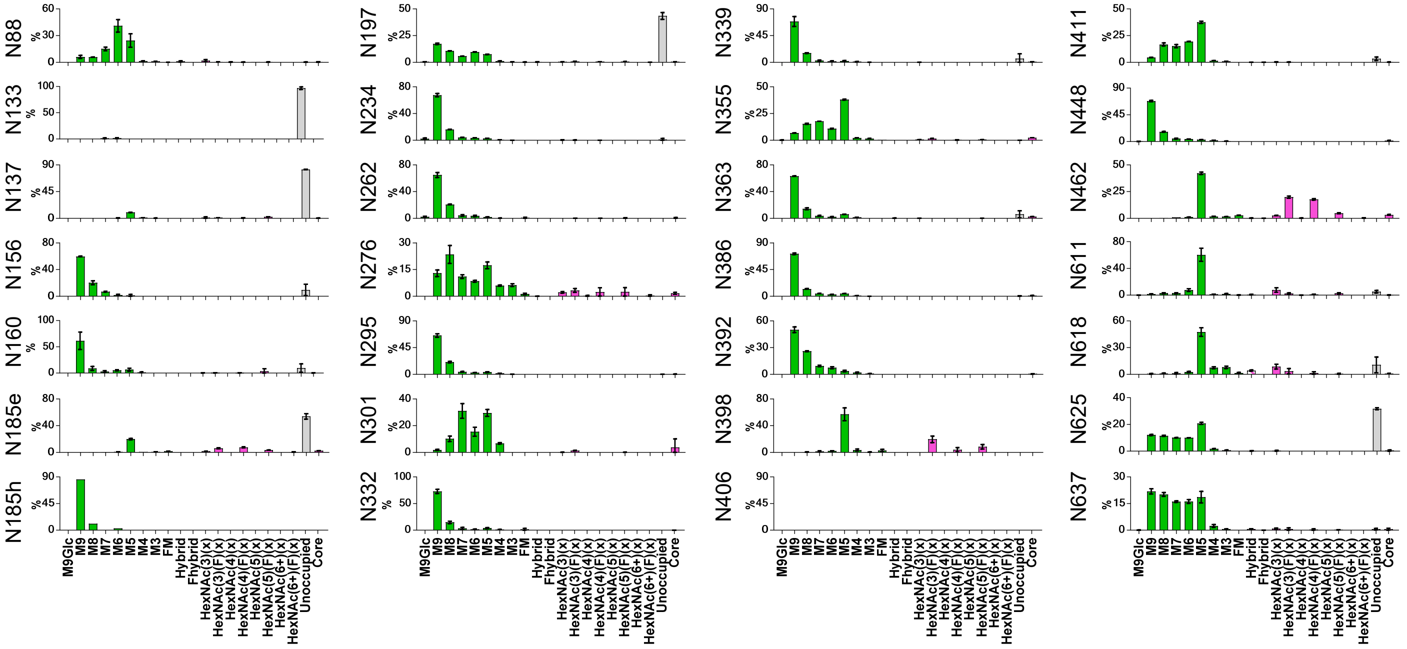


F WT BG505 UFO-10GS-I3-01v9-L7P SApNP

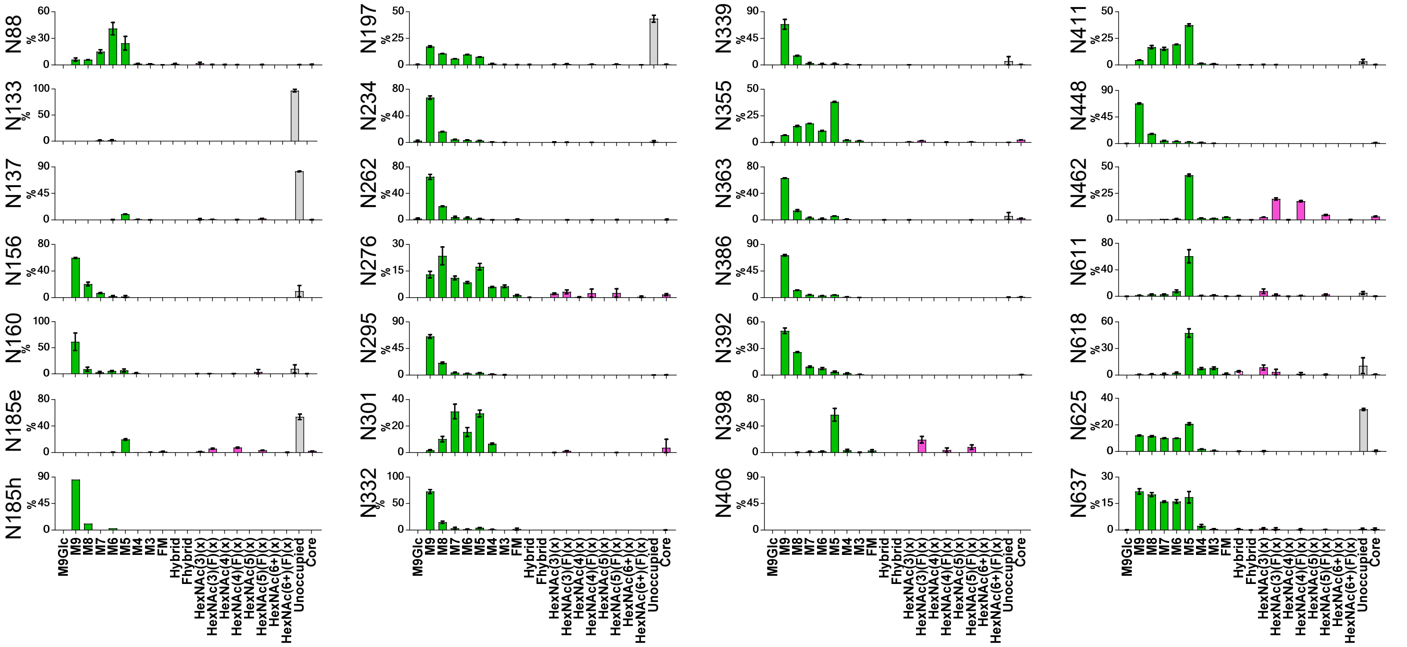

#### fig. S3 (continued)

G

**Summary of $OD_{450}$ values in ELISA of glycan-trimmed I3-01v9 and E2p SApNPs binding to an anti-MBP antibody and bNAb VRC01[a].**

**A. BG505 UFO-10GS-I3-01v9-L7P SApNP**

| | μg/ml | No endo H treatment | Endo H-treated sample-1 | Endo H-treated sample-2 | Endo H-treated sample-3 | Endo H-treated sample-4 | Endo H-treated sample-5 | Endo H-treated sample-6 | Endo H-treated sample-7 | Endo H-treated sample-8 | Endo H-treated sample-9 | Endo H-treated sample-10 | Endo Hf |
|---|---|---|---|---|---|---|---|---|---|---|---|---|---|
| VRC01 | 1 | 2.61 | 2.77 | 2.66 | 2.72 | 2.72 | 2.75 | 2.81 | 2.79 | 2.70 | 2.74 | 2.62 | 0.09 |
| | 0.1 | 2.63 | 2.73 | 2.73 | 2.60 | 2.82 | 2.80 | 2.71 | 2.69 | 2.72 | 2.71 | 2.51 | 0.06 |
| | 0.01 | 1.87 | 2.18 | 2.12 | 2.08 | 2.22 | 2.16 | 2.08 | 2.16 | 2.07 | 2.15 | 2.16 | 0.05 |
| | 0.001 | 0.45 | 0.69 | 0.69 | 0.66 | 0.64 | 0.66 | 0.65 | 0.64 | 0.63 | 0.66 | 0.64 | 0.06 |

| | dilution | No endo H treatment | Endo H-treated sample-1 | Endo H-treated sample-2 | Endo H-treated sample-3 | Endo H-treated sample-4 | Endo H-treated sample-5 | Endo H-treated sample-6 | Endo H-treated sample-7 | Endo H-treated sample-8 | Endo H-treated sample-9 | Endo H-treated sample-10 | Endo Hf |
|---|---|---|---|---|---|---|---|---|---|---|---|---|---|
| MBP mAb | ×466 | 0.09 | 0.09 | 0.10 | 0.10 | 0.09 | 0.11 | **0.15** | 0.10 | 0.10 | 0.07 | 0.06 | 2.32 |
| | ×4660 | 0.05 | 0.08 | 0.10 | 0.10 | 0.08 | 0.10 | 0.13 | 0.09 | 0.09 | 0.06 | 0.06 | 2.10 |
| | ×46600 | 0.05 | 0.06 | 0.08 | 0.07 | 0.06 | 0.06 | 0.08 | 0.07 | 0.06 | 0.05 | 0.05 | 0.89 |
| | ×466000 | 0.05 | 0.06 | 0.06 | 0.06 | 0.05 | 0.06 | 0.06 | 0.06 | 0.05 | 0.05 | 0.05 | 0.25 |

**B. BG505 UFO-E2p-L4P SApNP**

| | μg/ml | No endo H treatment | Endo H-treated sample-1 | Endo H-treated sample-2 | Endo H-treated sample-3 | Endo H-treated sample-4 | Endo H-treated sample-5 | Endo H-treated sample-6 | Endo H-treated sample-7 | Endo H-treated sample-8 | Endo H-treated sample-9 | Endo H-treated sample-10 | Endo Hf |
|---|---|---|---|---|---|---|---|---|---|---|---|---|---|
| VRC01 | 1 | 2.78 | 2.78 | 2.82 | 2.76 | 2.62 | 2.63 | 2.71 | 2.68 | 2.80 | 2.69 | 2.72 | 0.09 |
| | 0.1 | 2.60 | 2.76 | 2.53 | 2.63 | 2.61 | 2.65 | 2.48 | 2.61 | 2.72 | 2.67 | 2.69 | 0.06 |
| | 0.01 | 1.70 | 2.14 | 2.18 | 2.24 | 2.13 | 2.10 | 2.10 | 2.02 | 2.18 | 2.12 | 1.96 | 0.05 |
| | 0.001 | 0.38 | 0.62 | 0.65 | 0.68 | 0.62 | 0.66 | 0.59 | 0.62 | 0.63 | 0.61 | 0.47 | 0.06 |

| | dilution | No endo H treatment | Endo H-treated sample-1 | Endo H-treated sample-2 | Endo H-treated sample-3 | Endo H-treated sample-4 | Endo H-treated sample-5 | Endo H-treated sample-6 | Endo H-treated sample-7 | Endo H-treated sample-8 | Endo H-treated sample-9 | Endo H-treated sample-10 | Endo Hf |
|---|---|---|---|---|---|---|---|---|---|---|---|---|---|
| MBP mAb | ×466 | 0.06 | 0.18 | 0.24 | 0.22 | 0.20 | 0.19 | 0.15 | **0.30** | 0.20 | 0.18 | 0.17 | 2.32 |
| | ×4660 | 0.06 | 0.13 | 0.17 | 0.17 | 0.16 | 0.14 | 0.12 | 0.25 | 0.16 | 0.13 | 0.13 | 2.10 |
| | ×46600 | 0.07 | 0.08 | 0.10 | 0.10 | 0.09 | 0.08 | 0.07 | 0.13 | 0.09 | 0.08 | 0.07 | 0.89 |
| | ×466000 | 0.07 | 0.06 | 0.07 | 0.07 | 0.07 | 0.06 | 0.06 | 0.08 | 0.06 | 0.06 | 0.06 | 0.25 |

[a] Wildtype SApNP (without endo H treatment) and endo Hf (a fusion of endo H and MBP) were included in the ELISA binding analysis as controls, shown in the first and last columns respectively.

**fig S3. Site-specific glycan analysis of BG505 UFO trimer and SApNPs.** Site-specific glycan profiles are shown for the following immunogens: (**A**) WT BG505 UFO trimer. (**B**) BG505 UFO trimer expressed in the presence of Swainsonine, (**C**) BG505 UFO trimer expressed in the presence of kifunensine, (**D**) WT BG505 UFO-FR SApNP, (**E**) WT BG505 UFO-E2p-L4P SApNP, and (**F**) WT BG505 UFO-10GS-I3-01v9-L7P SApNP. **(G)** The OD450 readout obtained from an ELISA binding test of endo H-treated SApNPs against an MBP-specific mouse antibody and bNAb VRC01. Ten samples generated from different productions run were tested in this analysis. Four concentrations were tested for VRC01, while four dilutions were used in the ELISA test against an MBP-specific mouse antibody (The initial ×466 dilution was based on the manufacturer's recommendation). The OD450 values obtained from the highest antibody concentration in the MBP mAb binding assay are highlighted in grey shade, with the highest OD450 values colored in red.

**fig. S4**

A **WT BG505 UFO trimer (AHC)**

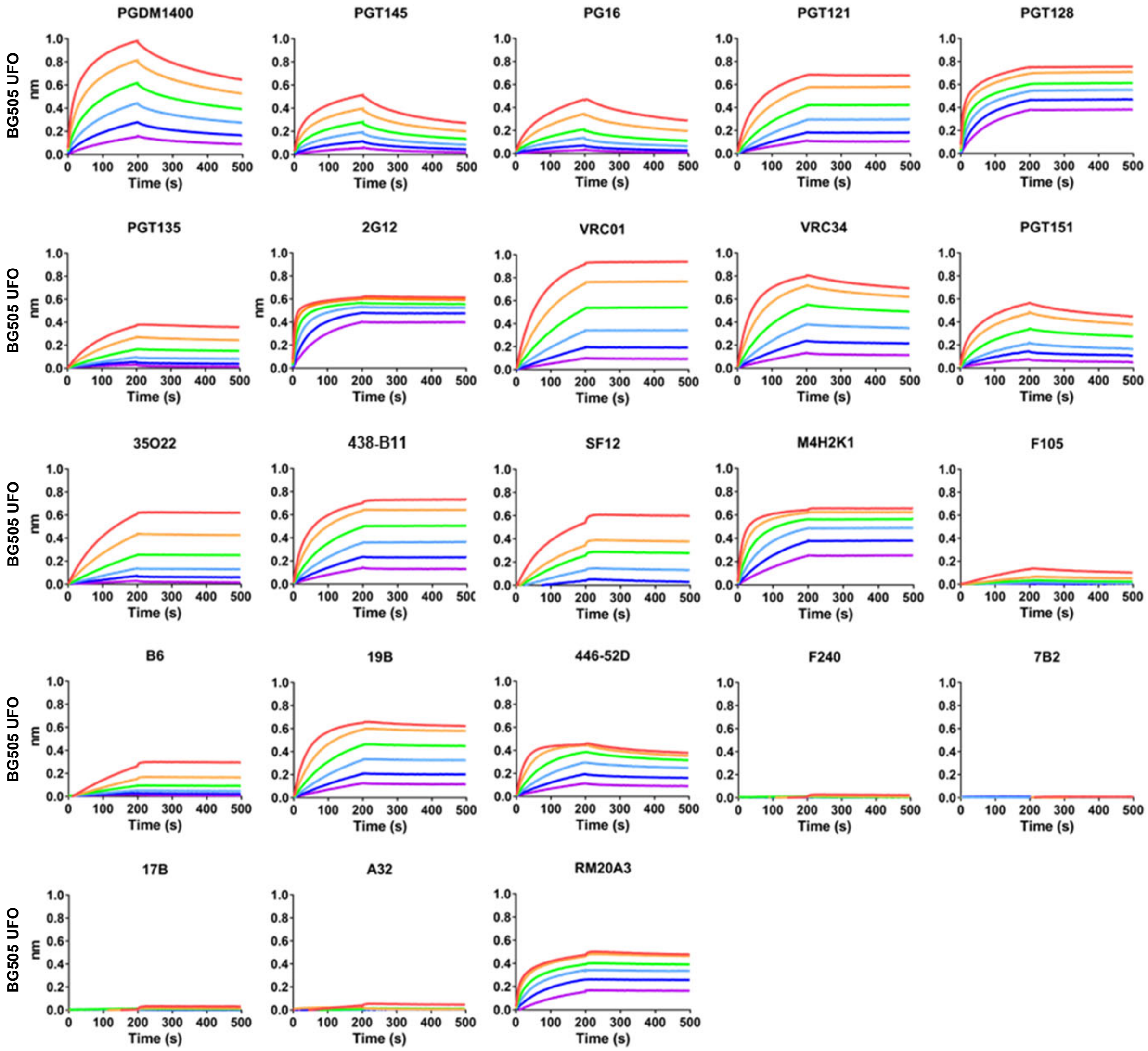

**fig. S4 (continued 1)**

B **BG505 UFO trimer expressed in the presence of kifunensine (AHC)**

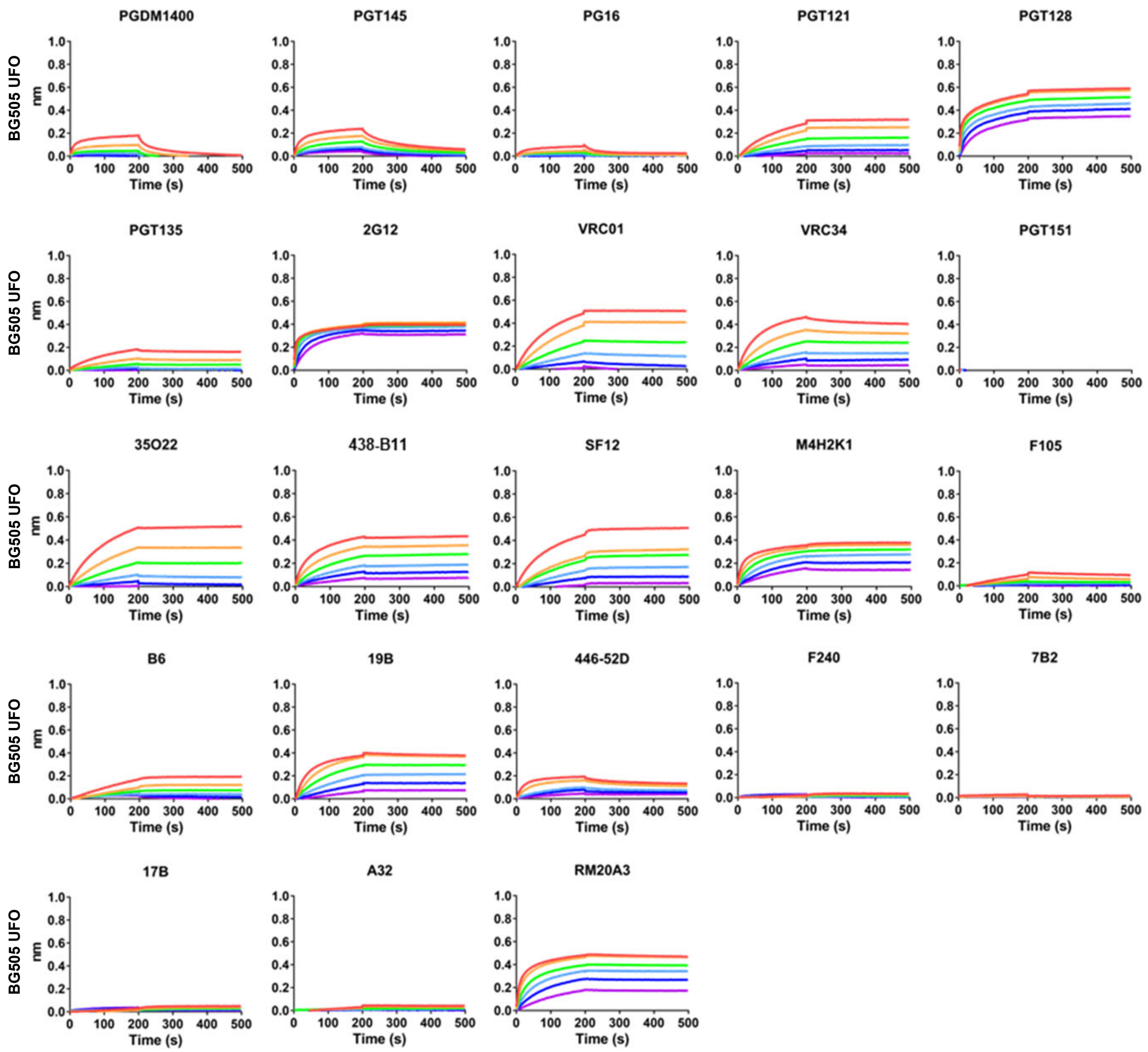

C **BG505 UFO trimer treated by endo H (AHC)**

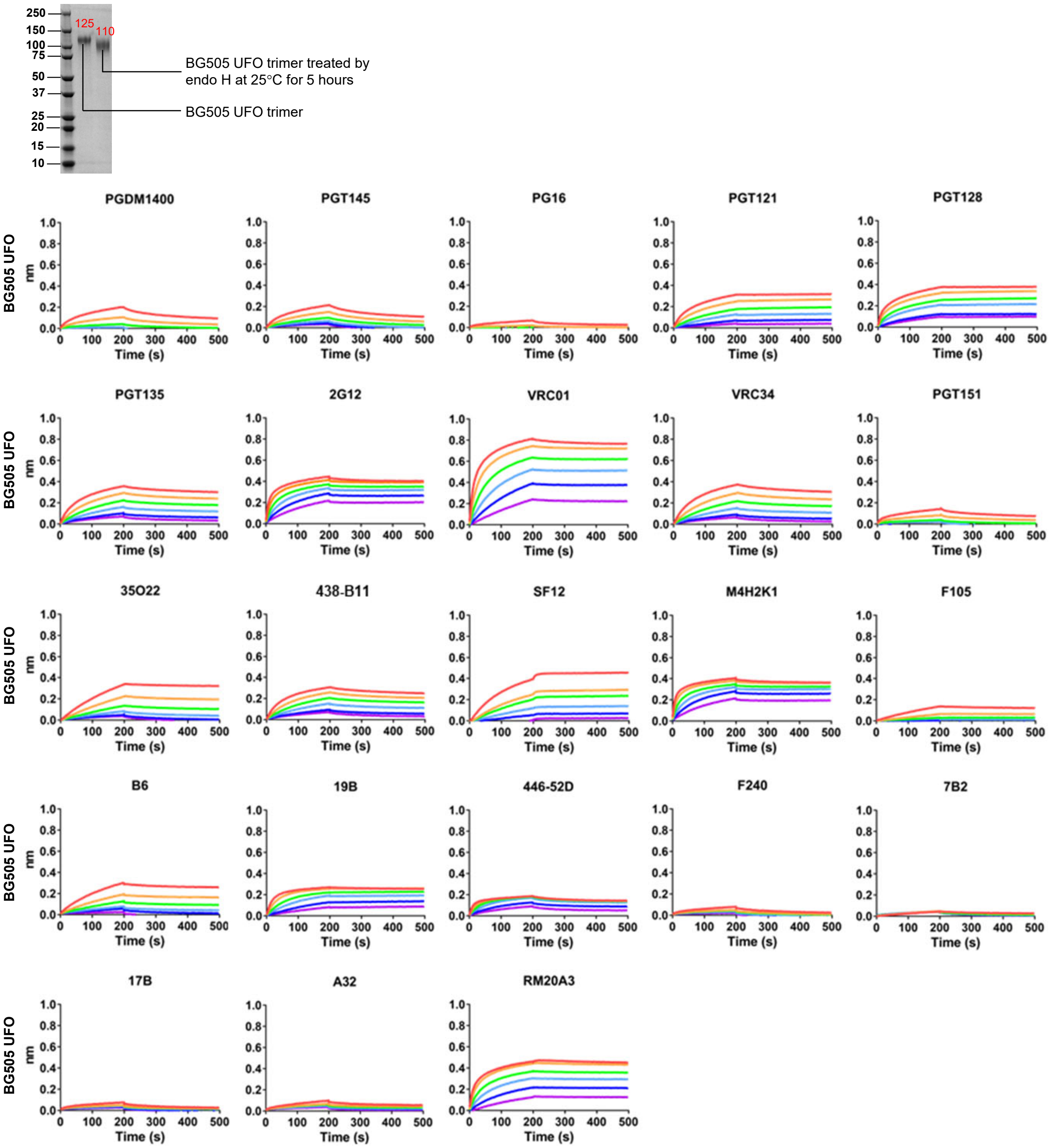

D WT BG505 UFO trimer (AHQ)

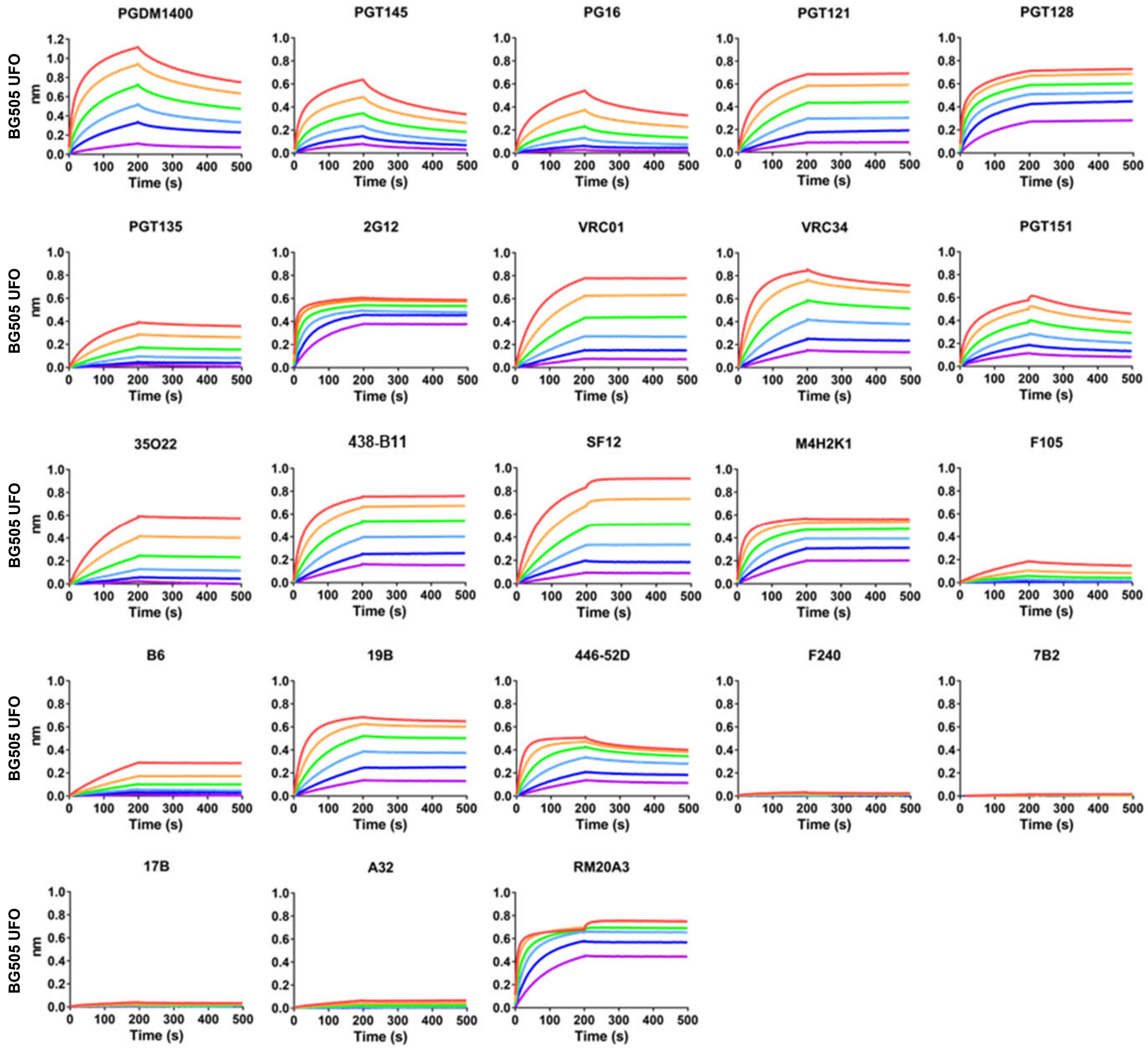

E **WT BG505 UFO trimer-presenting FR SApNP (AHQ)**

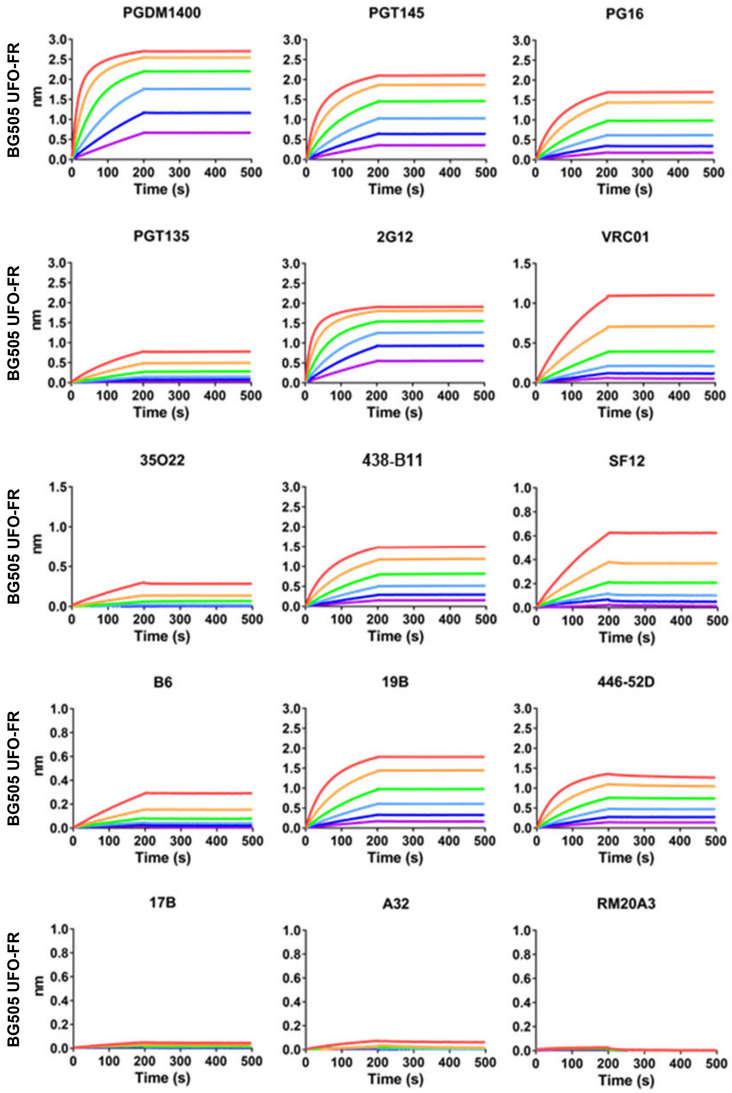

F WT BG505 UFO trimer-presenting E2p-L4P SApNP (AHQ)

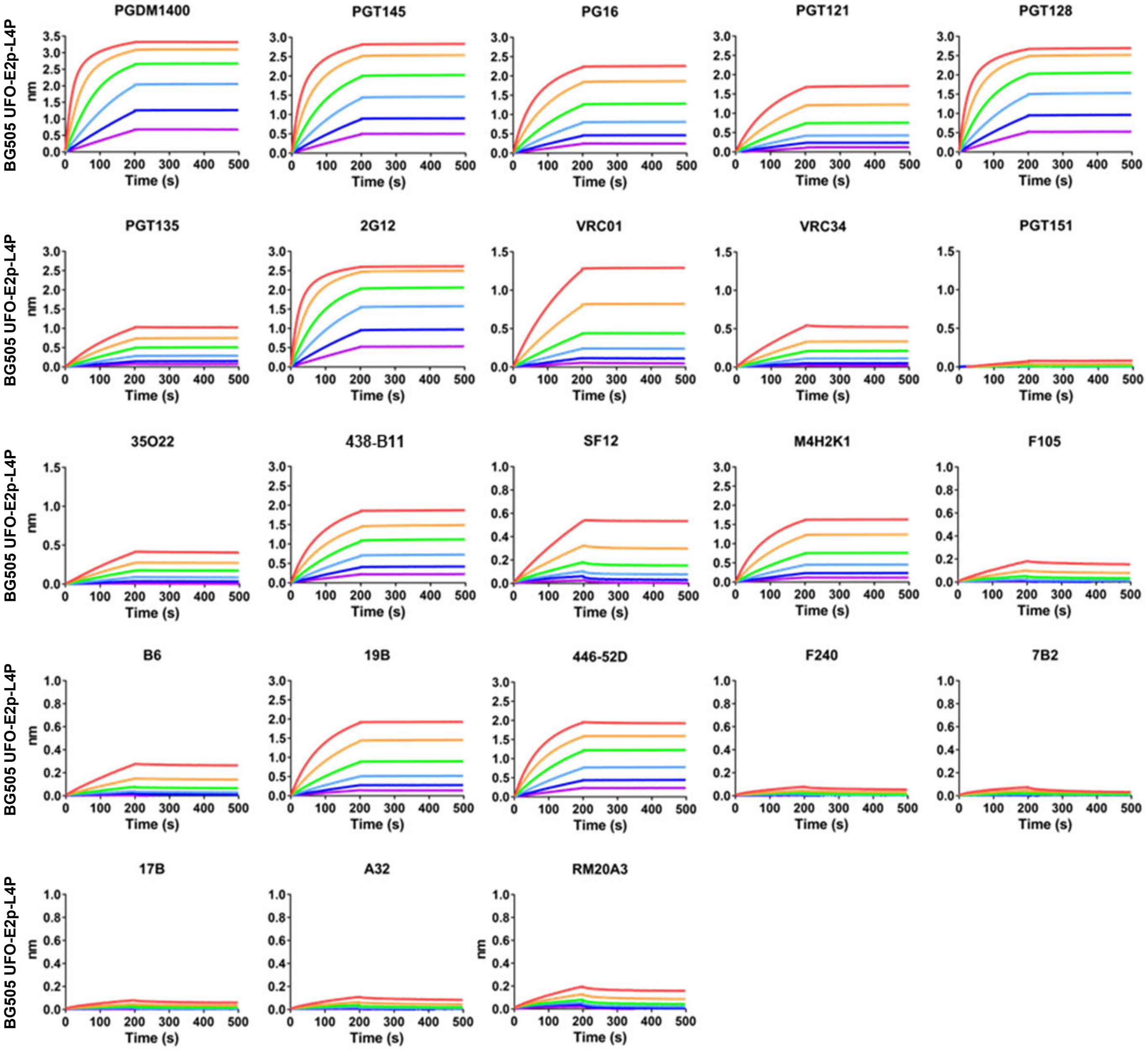

G **WT BG505 UFO trimer-presenting I3-01v9 SApNP (AHQ)**

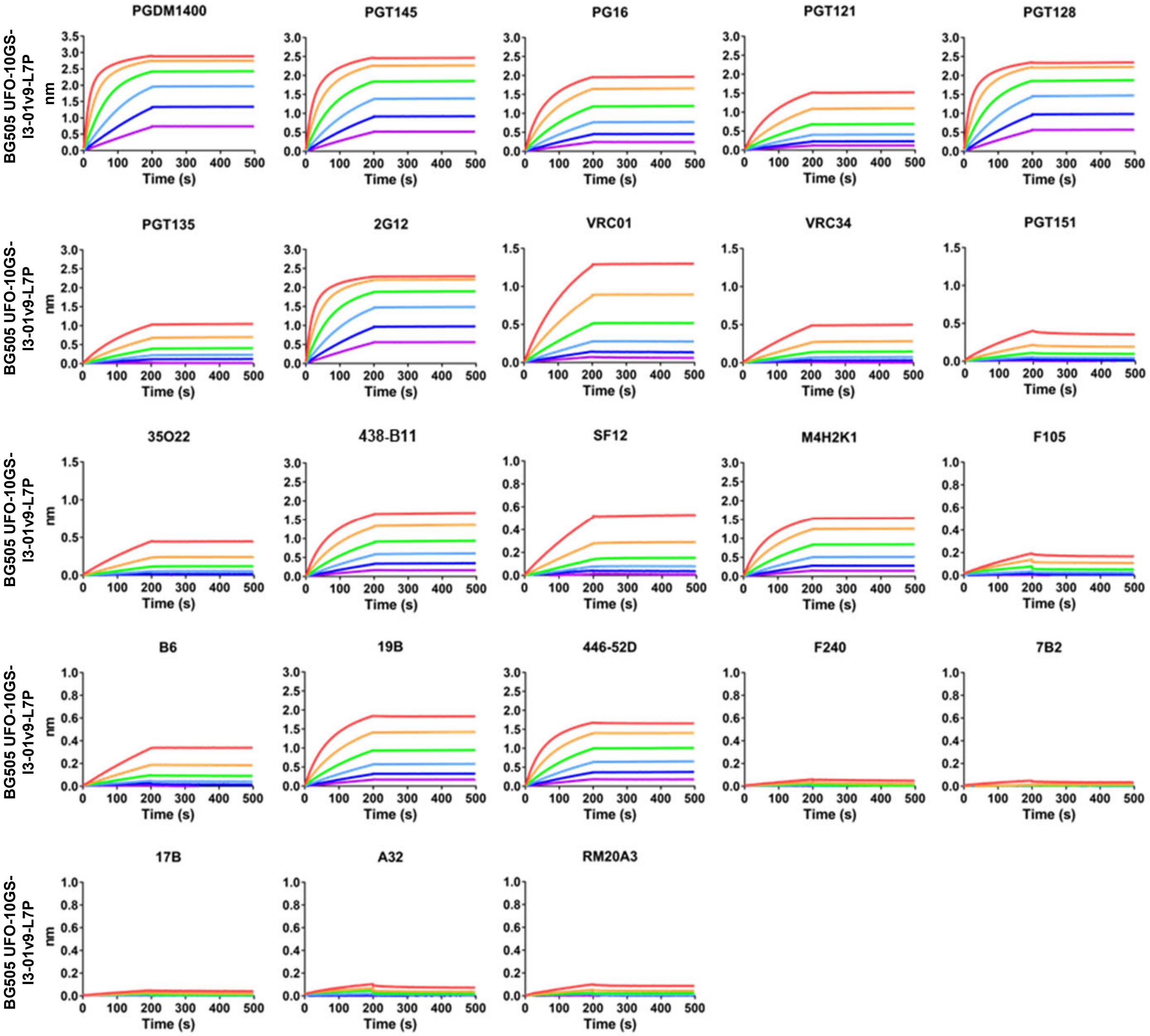

H **BG505 UFO trimer expressed in the presence of kifunensine (AHQ)**

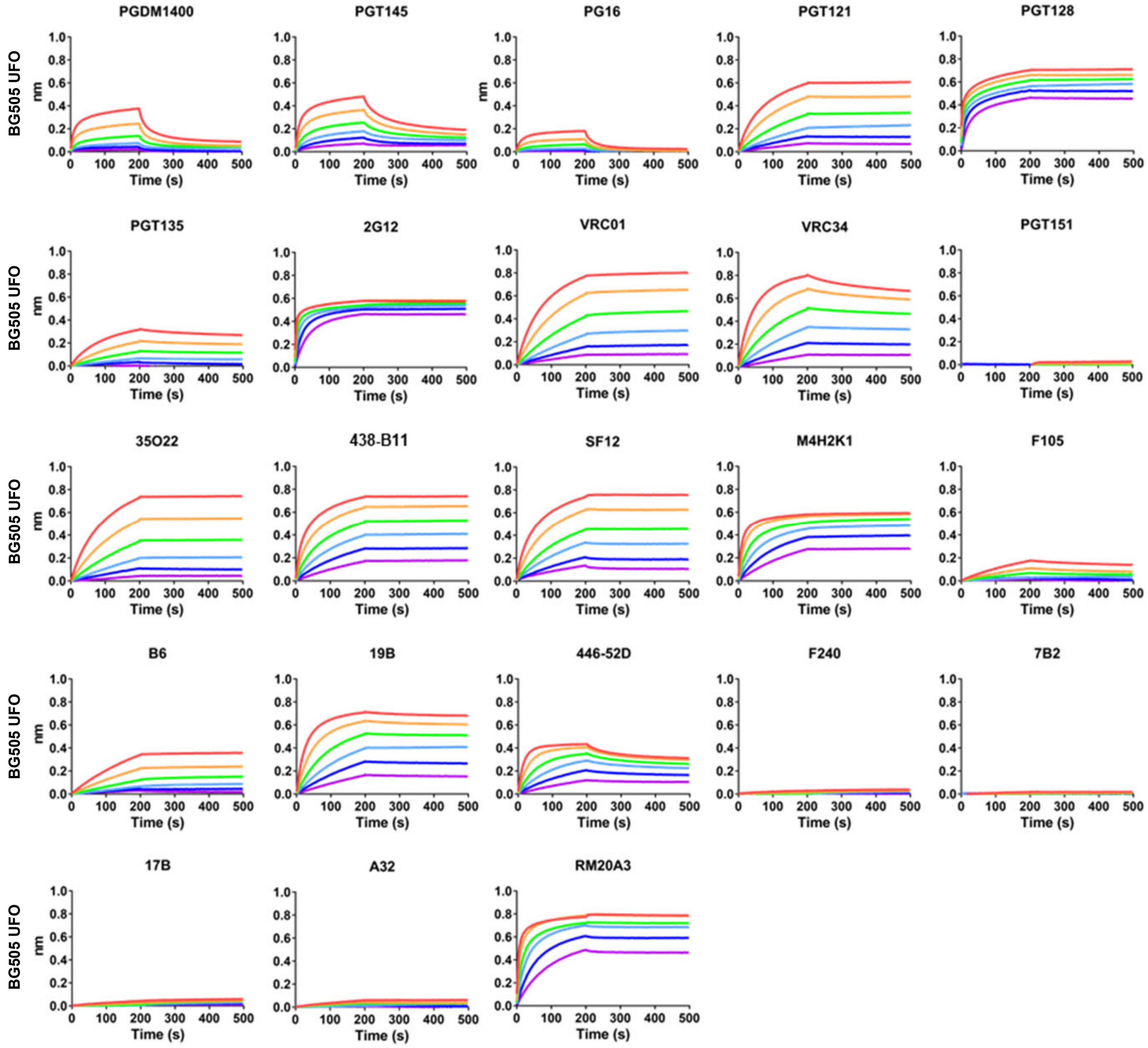

#### fig. S4 (continued 8)

**BG505 UFO trimer-presenting FR SApNP expressed in the presence of kifunensine (AHQ)**

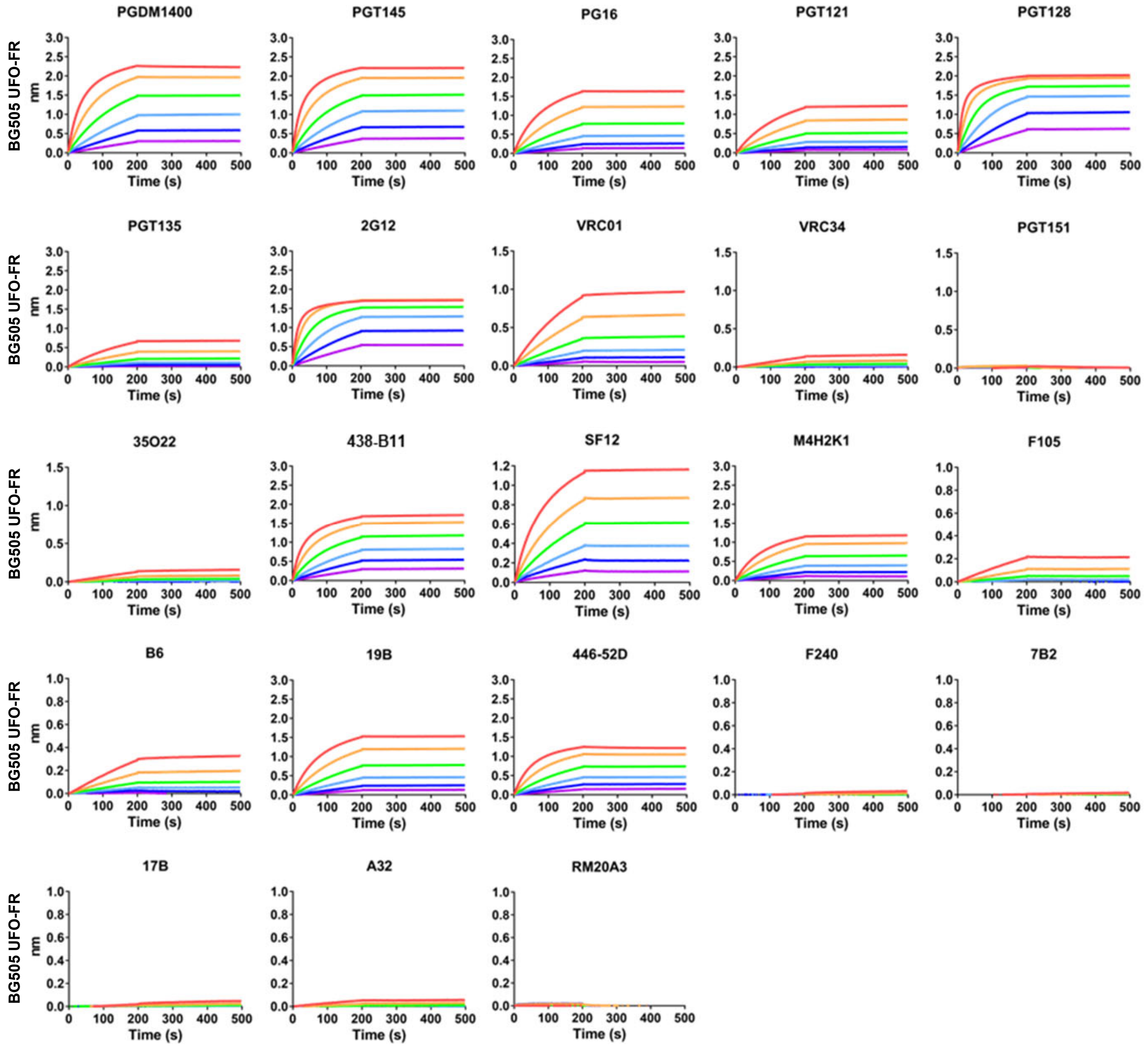

fig. S4 (continued 9)

J BG505 UFO trimer-presenting E2p-L4P SApNP expressed in the presence of kifunensine (AHQ)

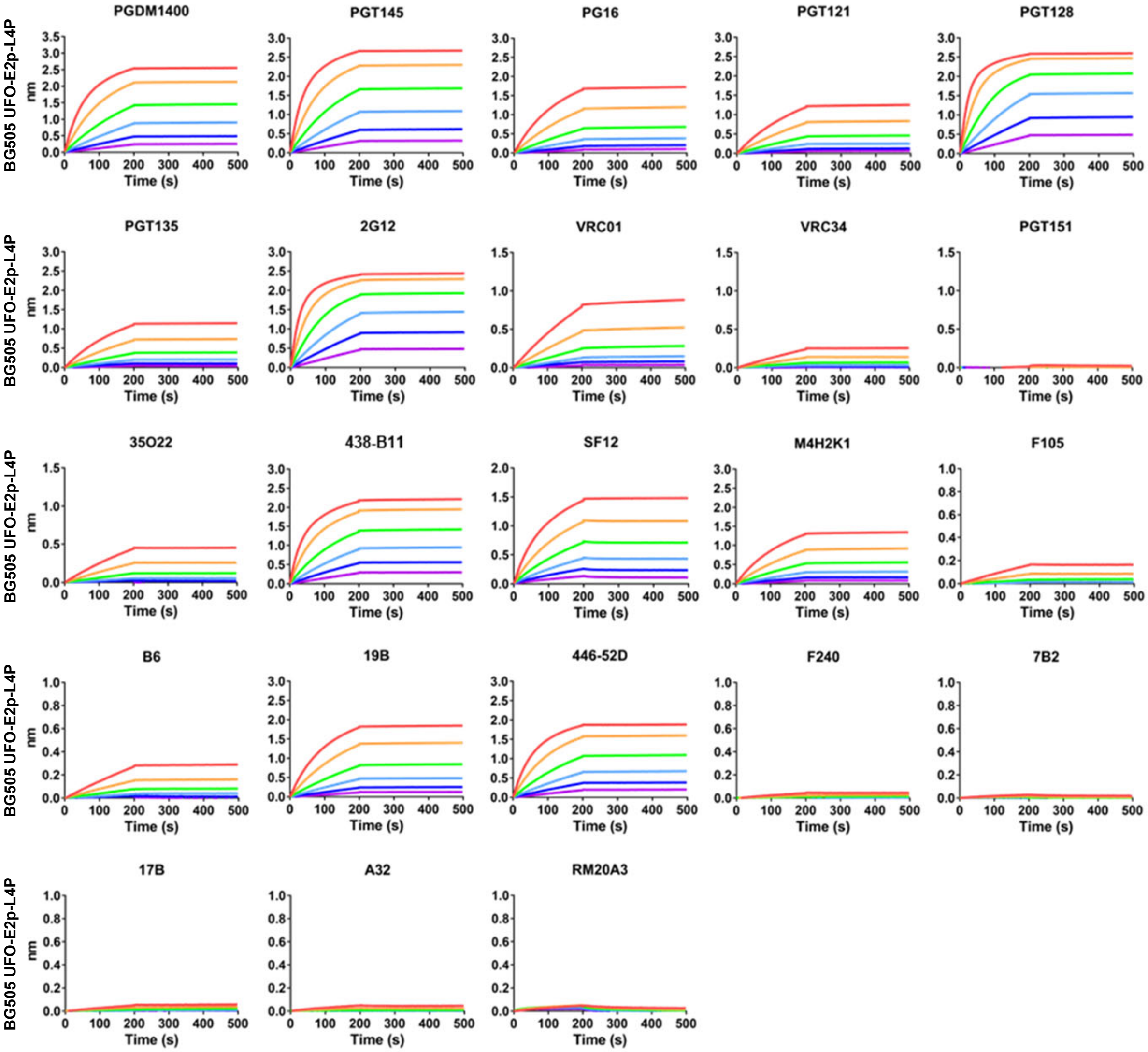

fig. S4 (continued 10)

K BG505 UFO trimer-presenting I3-01v9 SApNP expressed in the presence of kifunensine (AHQ)

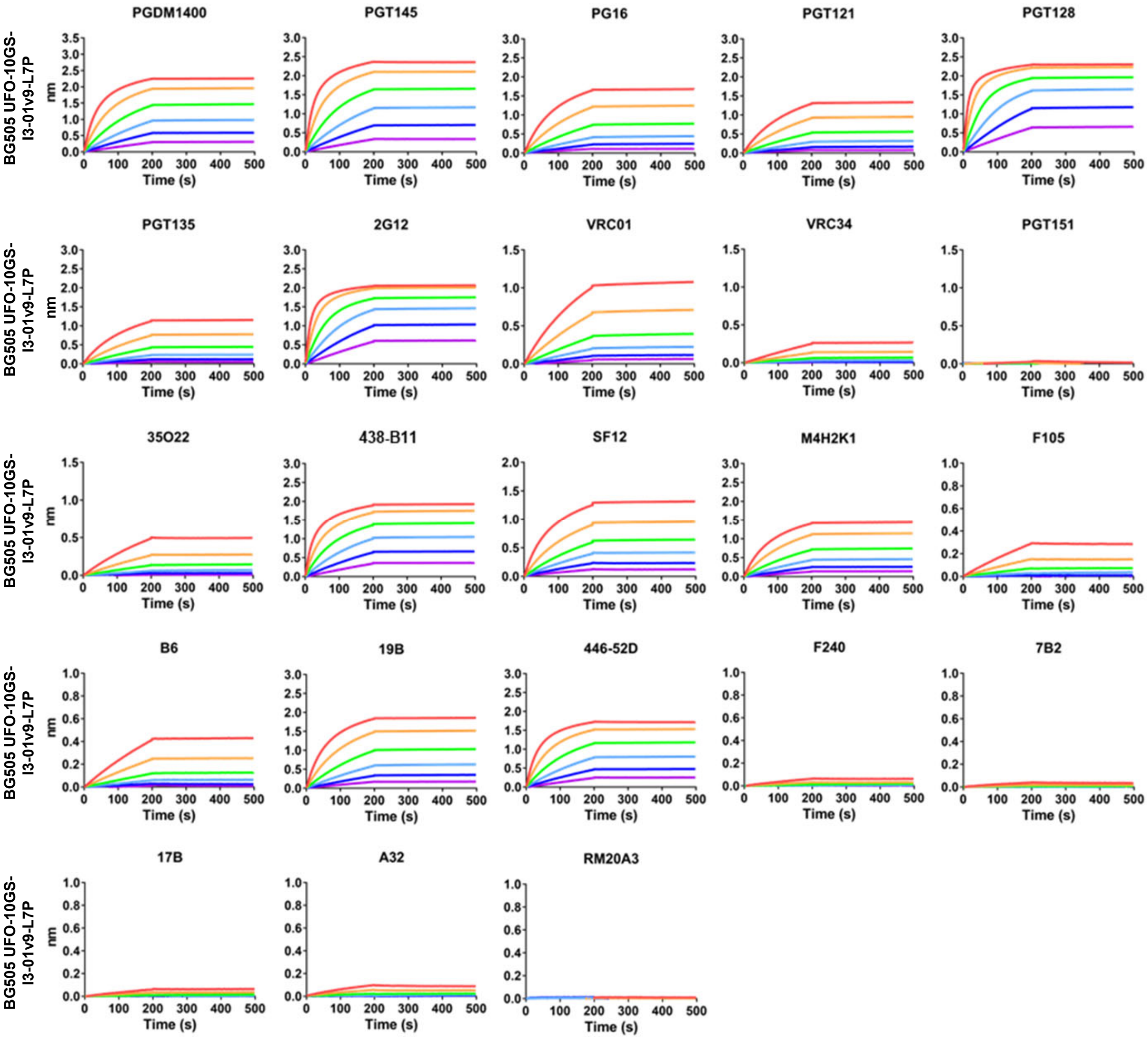

L **BG505 UFO trimer treated by endo H (AHQ)**

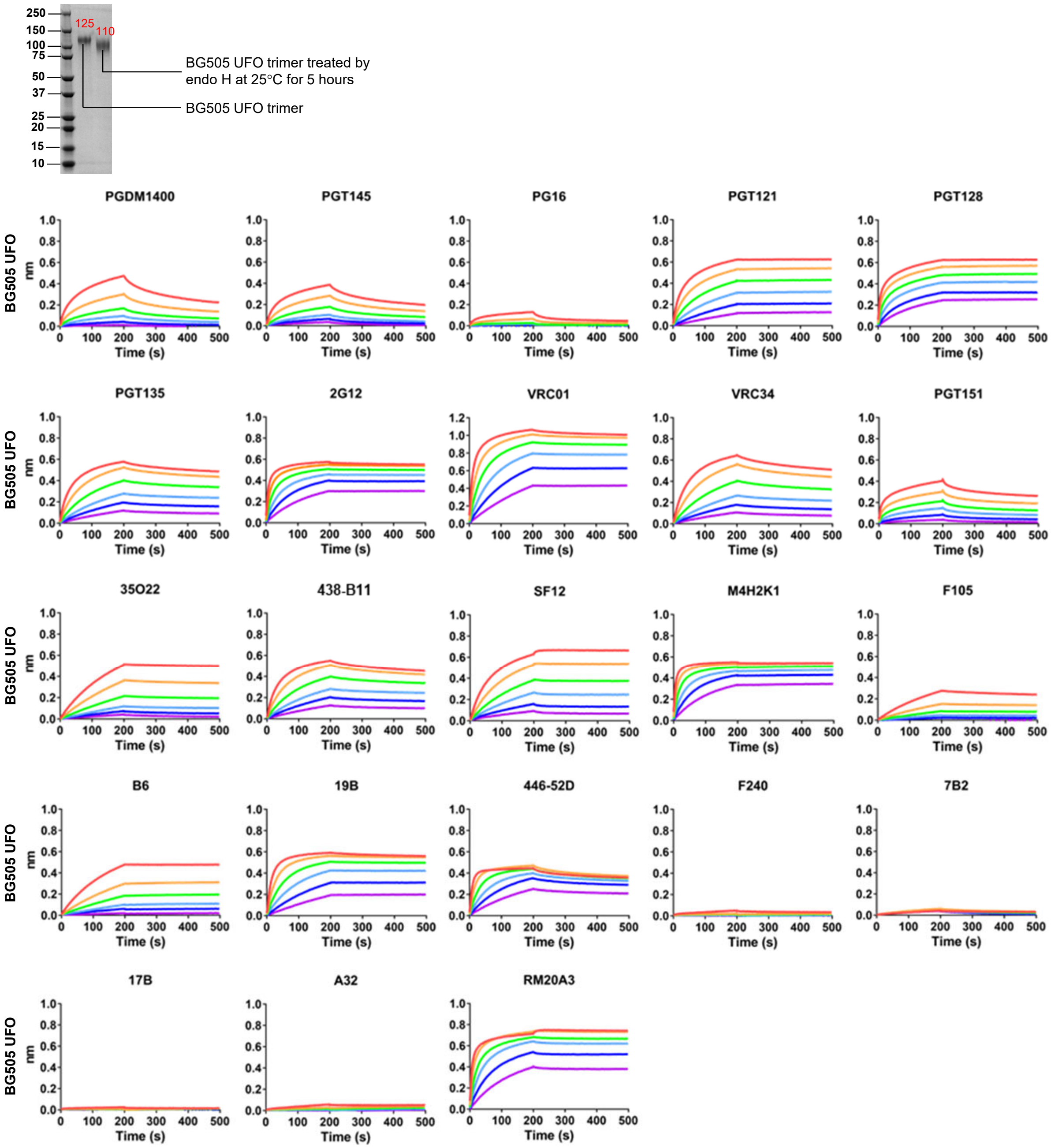

M **BG505 UFO trimer-presenting FR SApNP treated by endo H (AHQ)**

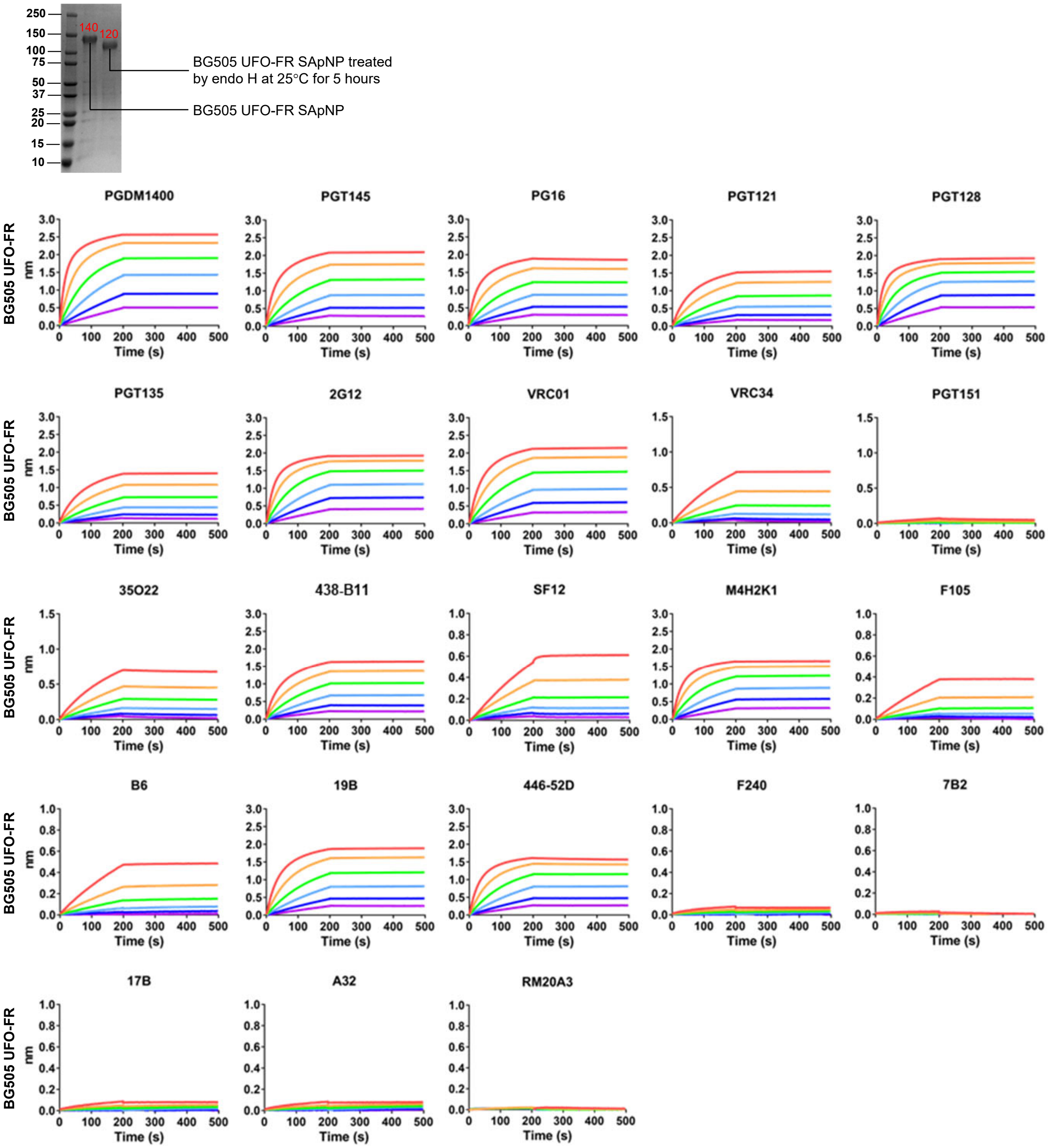

N **BG505 UFO trimer-presenting E2p-L4P SApNP treated by endo H (AHQ)**

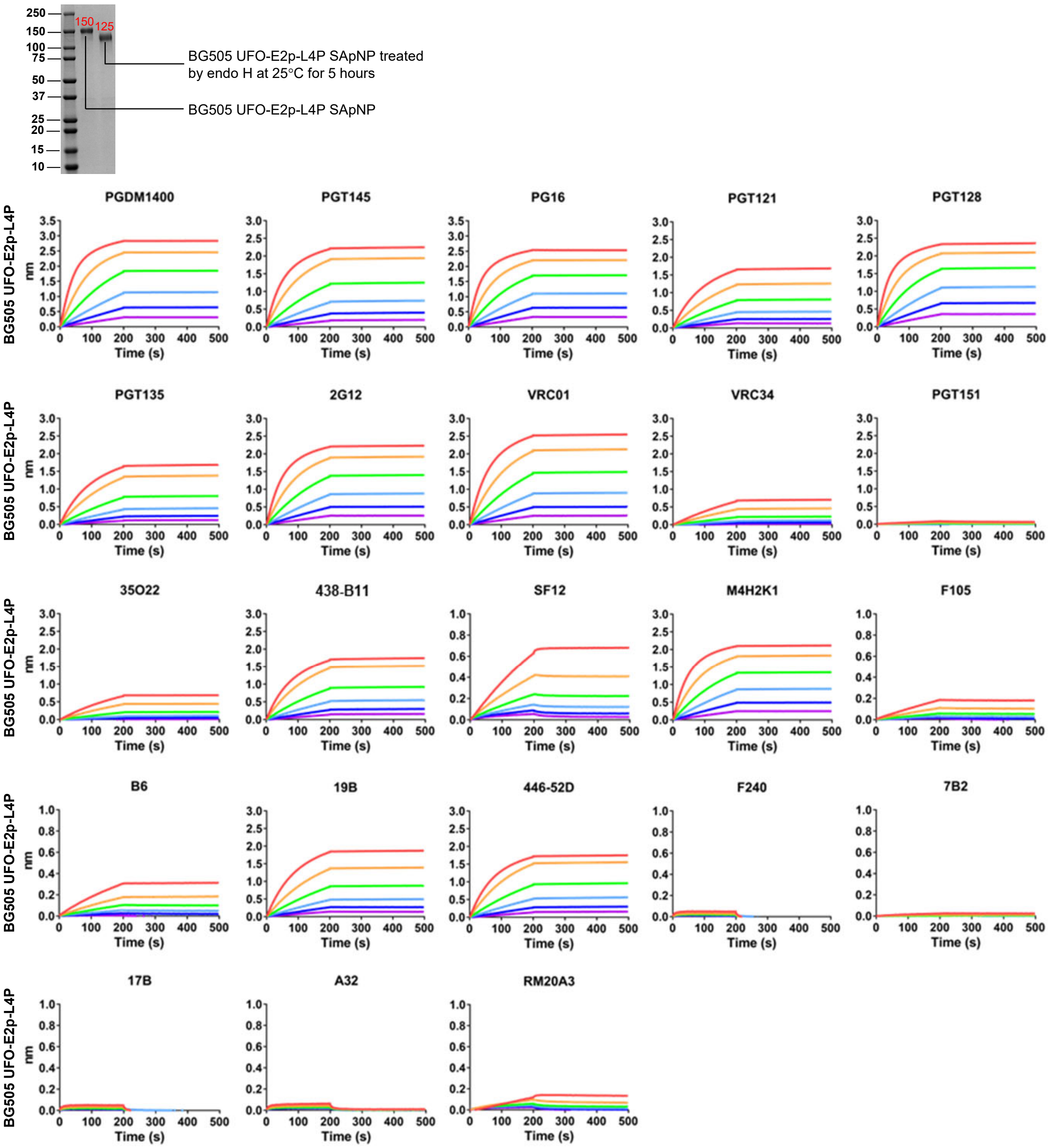

O BG505 UFO trimer-presenting I3-01v9-L7P SApNP treated by endo H (AHQ)

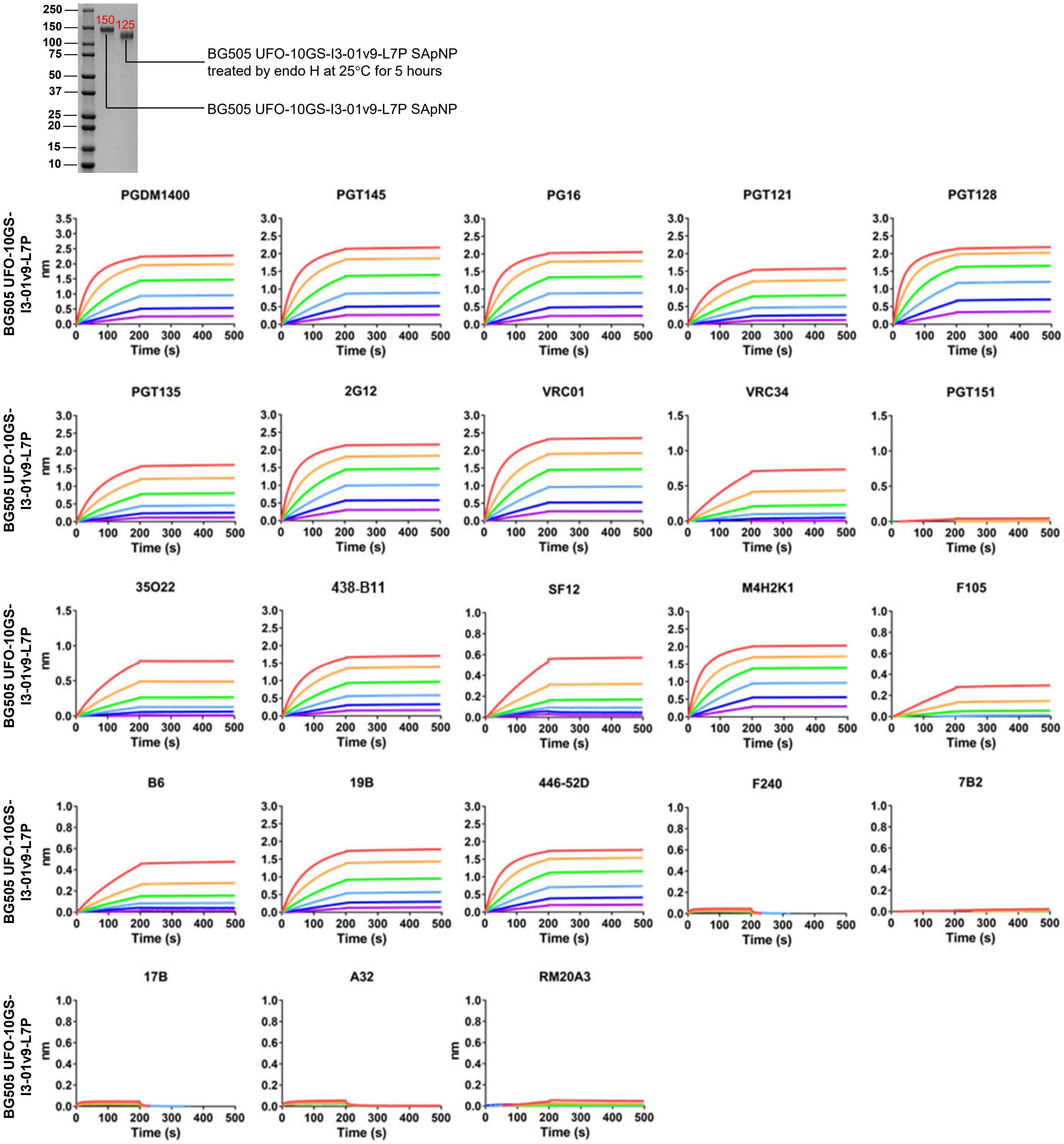

**P** **Signal variation in the BLI analysis of wildtype BG505 UFO-E2p-L4P SApNP**

a. Neutralizing and broadly neutralizing antibodies (NAbs and bNAbs)

| | PGDM 1400 | PGT145 | PG16 | PGT121 | PGT128 | PGT135 | 438-B11 | 2G12 | VRC01 | VRC34 | PGT151 | 35O22 | SF12 | M4H2K1 |
|---|---|---|---|---|---|---|---|---|---|---|---|---|---|---|
| Signal 1 (nm) | 3.117322 | 2.749335 | 2.026484 | 1.642852 | 2.363186 | 1.141498 | 1.959381 | 2.329419 | 1.264801 | 0.517218 | 0.130087 | 0.369583 | 0.550004 | 1.551745 |
| Signal 2 (nm) | 3.196049 | 2.818250 | 2.092079 | 1.700269 | 2.436687 | 1.108669 | 2.010078 | 2.273865 | 1.240167 | 0.509011 | 0.129068 | 0.372469 | 0.595204 | 1.508611 |
| Average (nm) | 3.156686 | 2.783793 | 2.059281 | 1.671561 | 2.399936 | 1.125083 | 1.984730 | 2.301642 | 1.252484 | 0.513114 | 0.129577 | 0.371026 | 0.572604 | 1.530178 |
| STDEV (nm) | 0.055669 | 0.048731 | 0.046383 | 0.040600 | 0.051973 | 0.023214 | 0.035848 | 0.039282 | 0.017419 | 0.005803 | 0.000721 | 0.002041 | 0.031961 | 0.030500 |
| Variation (%) | 1.76 | 1.75 | 2.25 | 2.43 | 2.17 | 2.06 | 1.81 | 1.71 | 1.39 | 1.13 | 0.56 | 0.55 | 5.58 | 1.99 |

b. Non-neutralizing antibodies (nNAbs)

| | F105 | b6 | 19b | 446-52D | F240 | 7B2 | 17b | A32 | RM20A3 |
|---|---|---|---|---|---|---|---|---|---|
| Signal 1 (nm) | 0.197451 | 0.244409 | 1.936790 | 1.969976 | 0.090558 | 0.066060 | 0.100006 | 0.073250 | 0.189037 |
| Signal 2 (nm) | 0.190804 | 0.239480 | 2.004852 | 2.032409 | 0.080123 | 0.075041 | 0.095142 | 0.067766 | 0.199055 |
| Average (nm) | 0.194128 | 0.241945 | 1.970821 | 2.001192 | 0.085341 | 0.070550 | 0.097574 | 0.070508 | 0.194046 |
| STDEV (nm) | 0.004700 | 0.003485 | 0.048127 | 0.044147 | 0.007379 | 0.006351 | 0.003440 | 0.003878 | 0.007084 |
| Variation (%) | 2.42 | 1.44 | 2.44 | 2.21 | 8.65 | 9.00 | 3.53 | 5.50 | 3.65 |

Octet signals, average, standard deviation are shown at the precision of $10^{-6}$, whereas variation (%) is shown at the precision level of $10^{-2}$.

**Q** **Negative EM analysis of glycan-trimmed BG505 UFO trimer in complex with bNAb VRC01**

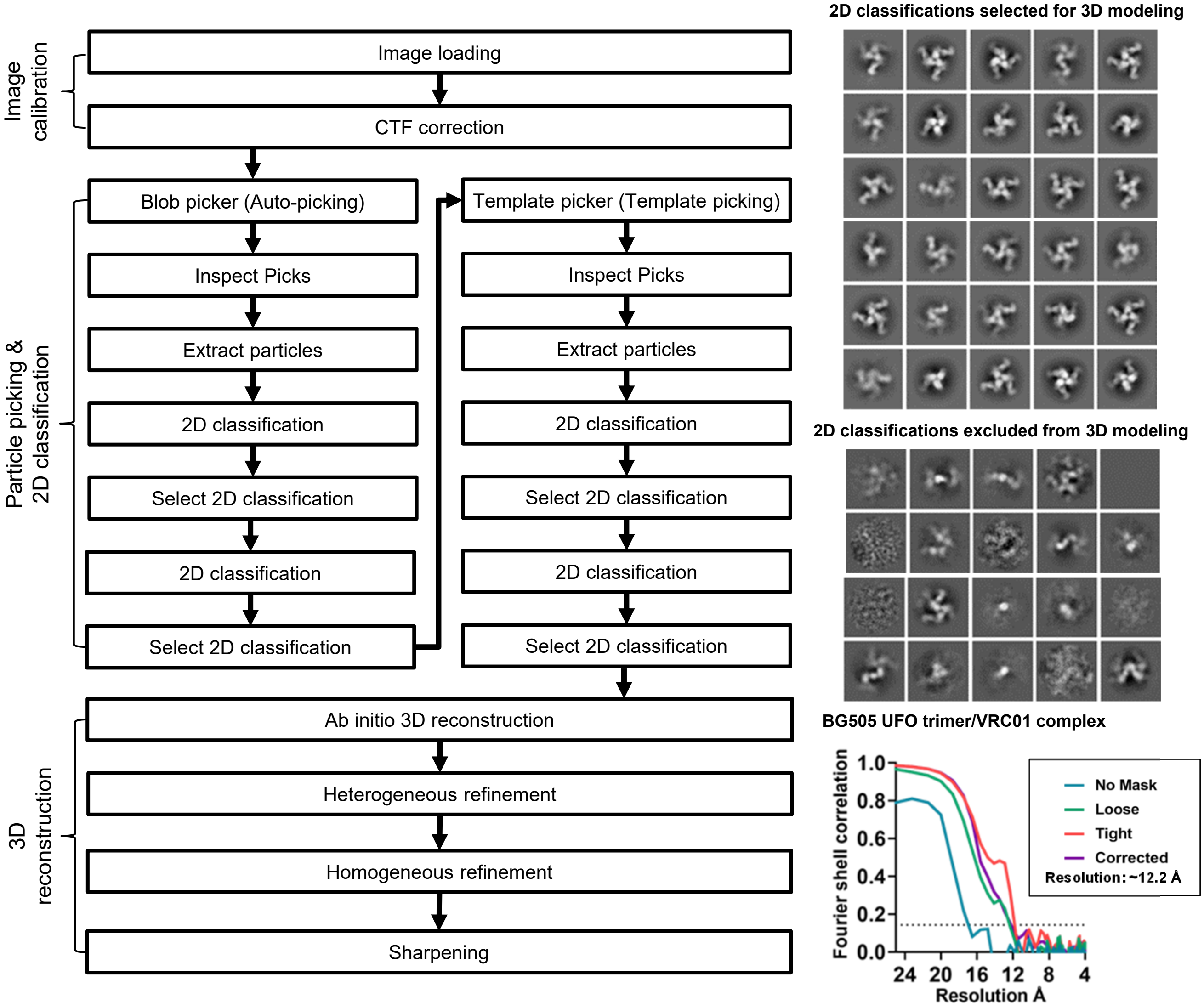

**fig. S4**. **Antigenic characterization of BG505 UFO trimer and trimer-presenting SApNPs with wildtype and modified glycans.** BG505 Env immunogens were tested against a panel of 23 HIV-1 antibodies including NAbs, bNAbs, and nNAbs by BLI. Sensorgrams were obtained from an Octet RED96 instrument using two types of biosensors, AHC and AHQ. The antigens tested here include (**A**) WT BG505 UFO trimer (AHC), (**B**) BG505 UFO trimer expressed in the presence of kifunensine (AHC), (**C**) BG505 UFO trimer treated by endo H (AHC), (**D**) WT BG505 UFO trimer (AHQ), (**E**) WT BG505 UFO trimer-presenting FR SApNP (AHQ), (**F**) WT BG505 UFO trimer-presenting E2p-L4P SApNP (AHQ), (**G**) WT BG505 UFO trimer-presenting I3-01v9-L7P SApNP (AHQ), (**H**) BG505 UFO trimer expressed in the presence of kifunensine (AHQ), (**I**) BG505 UFO trimer-presenting FR SApNP expressed in the presence of kifunensine (AHQ), (**J**) BG505 UFO trimer-presenting E2p-L4P SApNP expressed in the presence of kifunensine (AHQ), (**K**) BG505 UFO trimer-presenting I3-01v9-L7P SApNP expressed in the presence of kifunensine (AHQ), (**L**) BG505 UFO trimer treated by endo H (AHQ), (**M)** BG505 UFO trimer-presenting FR SApNP treated by endo H (AHQ), (**N**) BG505 UFO trimer-presenting E2p-L4P SApNP treated by endo H (AHQ), and (**O**) BG505 UFO trimer-presenting I3-01v9-L7P SApNP treated by endo H (AHQ). A two-fold concentration gradient of antigen, starting at 266.7 nM for the UFO trimer, 14.9 nM for the FR SApNP, and 5.5 nM for the E2p and I3-01v9 SApNPs, was used in a titration series of six. SDS-PAGE was run for (C), (L), (M), (N), and (O) to estimate molecular weight of glycan-trimmed vs. WT BG505 Env immunogens to ensure that molar concentration was comparable in their BLI assays. (**P**) Estimate of signal variation in the BLI analysis within a single Octet run. Wildtype BG505 UFO-E2p-L4P SApNP was tested in duplicate using the highest concentration (5.5 nM). Variation (%) was defined as STDEV/Average×100%. (**Q**) Negative-stain EM analysis of the glycan-trimmed BG505 UFO trimer in complex with bNAb VRC01. Left: flowchart for image processing. Right top: 2D classes selected for 3D reconstruction and excluded from 3D reconstruction. Right bottom: Estimated resolution (12.2 Å) of the 3D reconstruction for the BG505 UFO trimer/VRC01 complex was calculated from the Fourier shell correlation (FSC) using a cut-off of 0.143.

**fig. S5**

A **Week 11 mouse IgG neutralization against HIV-1 BG505.T332N**

| Vaccine | Adjuvant | IC50 titers (µg/ml) | | | | | | | |
|---|---|---|---|---|---|---|---|---|---|
| | | M1 | M2 | M3 | M4 | M5 | M6 | M7 | M8 |
| BG505 E2p SApNP | No adjuvant | >300 | >300 | >300 | >300 | No IgG | >300 | >300 | >300 |
| | AddaVax | >300 | >300 | >300 | 200 | >300 | >300 | No IgG | >300 |
| | Aluminum phosphate | >300 | 90 | No IgG | >300 | >300 | >300 | >300 | >300 |
| | Aluminum hydroxide | >300 | >300 | >300 | >300 | >300 | >300 | >300 | >300 |
| | AddaVax/Aluminum phosphate | >300 | >300 | >300 | >300 | >300 | >300 | >300 | >300 |
| BG505 I3-01v9 SApNP | No adjuvant | >300 | >300 | >300 | >300 | >300 | >300 | >300 | >300 |
| | AddaVax | >300 | >300 | >300 | >300 | >300 | >300 | >300 | >300 |
| | Aluminum phosphate | 21 | >300 | >300 | >300 | >300 | >300 | >300 | >300 |
| | Aluminum hydroxide | No IgG | >300 | >300 | >300 | >300 | 36 | >300 | >300 |
| | AddaVax/Aluminum phosphate | >300 | >300 | >300 | >300 | >300 | >300 | >300 | >300 |

No IgG: indicates no IgG available to test.

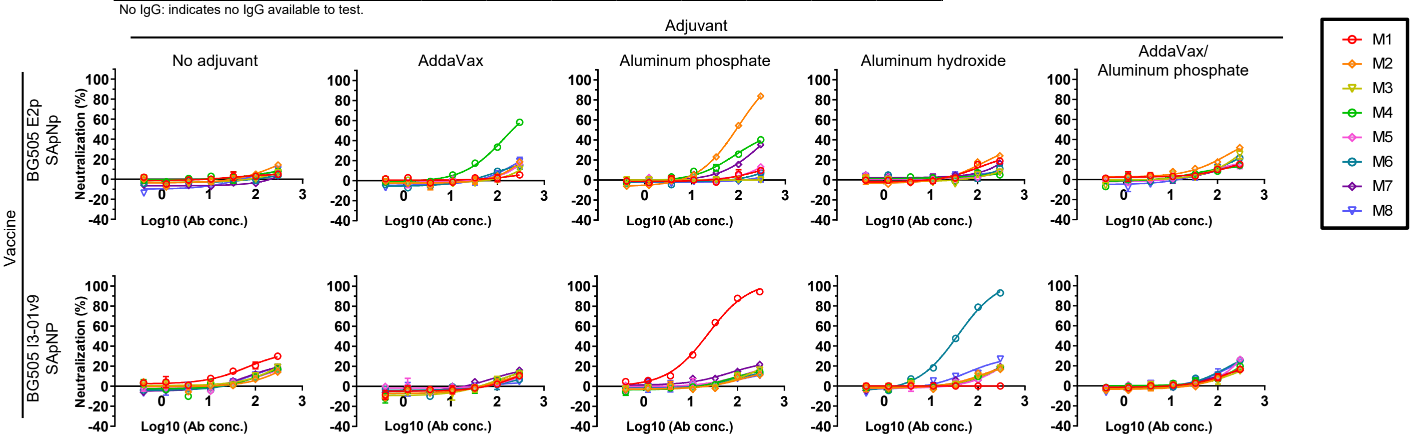


**Week 11 mouse IgG neutralization against HIV SF162**

| Vaccine | Adjuvant | IC50 titers (µg/ml) | | | | | | | |
|---|---|---|---|---|---|---|---|---|---|
| | | M1 | M2 | M3 | M4 | M5 | M6 | M7 | M8 |
| BG505 E2p SApNP | No adjuvant | >300 | >300 | >300 | >300 | - | >300 | >300 | >300 |
| | AddaVax | >300 | 26 | 18 | >300 | 250 | 13 | No IgG | >300 |
| | Aluminum phosphate | 4 | 34 | No IgG | 39 | 62 | >300 | 1 | 2 |
| | Aluminum hydroxide | 14 | 2 | 7 | 29 | 104 | 9 | >300 | 14 |
| | AddaVax/Aluminum phosphate | 77 | 83 | 20 | 287 | 4 | 1 | >300 | 35 |
| BG505 I3-01v9 SApNP | No adjuvant | >300 | >300 | >300 | 16 | >300 | >300 | >300 | 12 |
| | AddaVax | 146 | >300 | 11 | 2 | 79 | 26 | 11 | 89 |
| | Aluminum phosphate | 5 | 34 | 91 | >300 | 126 | 9 | 86 | 5 |
| | Aluminum hydroxide | No IgG | 9 | 41 | 4 | 45 | 28 | 15 | >300 |
| | AddaVax/Aluminum phosphate | 1 | 88 | 13 | 285 | >300 | 14 | >300 | 102 |

No IgG: indicates no IgG available to test.

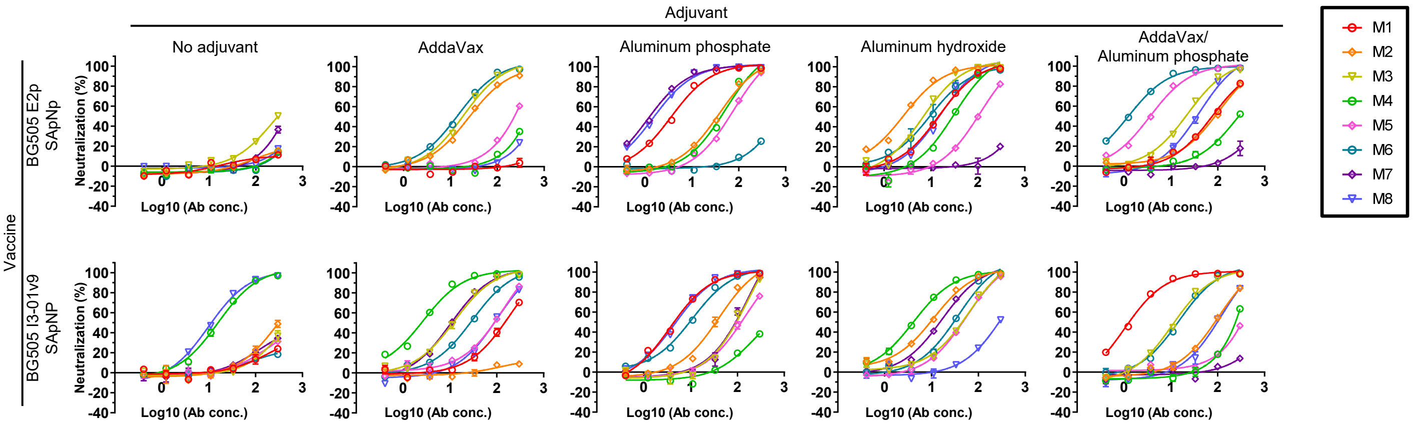


**Week 11 mouse IgG neutralization against MLV (negative control)**

| Vaccine | Adjuvant | Animal ID | IC50 titers (µg/ml) |
|---|---|---|---|
| BG505 E2p SApNP | Aluminum phosphate | D-S19 G3-M2 | >300 |
| BG505 I3-01v9 SApNP | Aluminum phosphate | D-S20 G3-M1 | >300 |
| | Aluminum hydroxide | D-S20 G4-M6 | >300 |

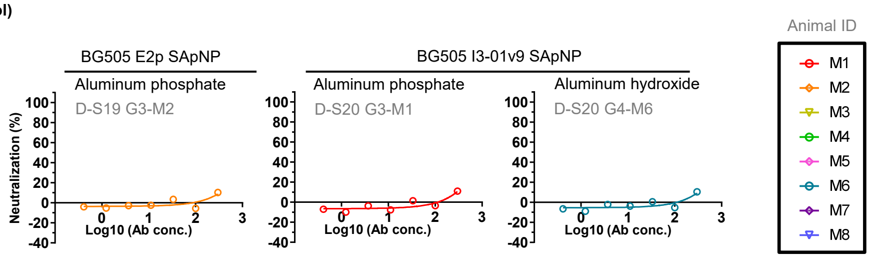

#### B Week 11 mouse IgG neutralization $IC_{50}$ titers against HIV-1 BG505.T332N and BG505.T332N.I396R mutant

| Vaccine | Adjuvant | Animal ID | $IC_{50}$ titers (µg/ml) | |
|---|---|---|---|---|
| | | | BG505. T332N | BG505. T332N. I396R |
| BG505 E2p SApNP | AddaVax | D-S19 G2-M4* | 200 | >300 |
| | Aluminum phosphate | D-S19 G3-M2 | 90 | >300 |
| BG505 I3-01v9 SApNP | Aluminum phosphate | D-S20 G3-M1 | 21 | 293 |
| | Aluminum hydroxide | D-S20 G4-M6 | 36 | >300 |

* starting concentration for this sample was at 95 µg/ml against the BG505.T332N I396R as inadequate sample left to start at 300 µg/ml

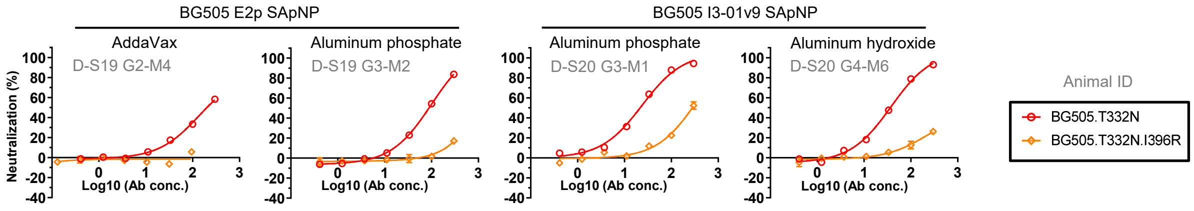


#### C Week 11 mouse IgG neutralization $IC_{50}$ and $IC_{30}$ titers against HIV-1 12 virus panel, BG505.T332N, SF162 & MLV (control)

| Vaccine | Adjuvant | Animal ID | $IC_{50}$ titers (µg/ml) for each virus (clade) | | | | | | | | | | | | | | |
|---|---|---|---|---|---|---|---|---|---|---|---|---|---|---|---|---|---|
| | | | TRO11 (B) | 25710 (C) | 398F1 (A) | CNE8 (AE) | X2278 (B) | BJOX-2000 (BC) | X1632 (G) | CE1176 (C) | 246F3 (AC) | CH119 (BC) | CE0217 (C) | CNE55 (AE) | BG505. T332N (A) | SF162 (B) | MLV (control) |
| BG505 E2p SApNP | Aluminum phosphate | D-S19 G3-M2 | >300 | >300 | >300 | >300 | >300 | >300 | >300 | >300 | >300 | >300 | >300 | >300 | 90 | 34 | >300 |
| BG505 I3-01v9 SApNP | Aluminum phosphate | D-S20 G3-M1 | >300 | >300 | 269 | >300 | >300 | >300 | >300 | >300 | >300 | >300 | >300 | >300 | 21 | 5 | >300 |
| | Aluminum hydroxide | D-S20 G4-M6 | >300 | >300 | >300 | >300 | >300 | >300 | >300 | >300 | >300 | >300 | >300 | >300 | 36 | 28 | >300 |

| Vaccine | Adjuvant | Animal ID | $IC_{30}$ titers (µg/ml) for each virus (clade) | | | | | | | | | | | | | | |
|---|---|---|---|---|---|---|---|---|---|---|---|---|---|---|---|---|---|
| | | | TRO11 (B) | 25710 (C) | 398F1 (A) | CNE8 (AE) | X2278 (B) | BJOX-2000 (BC) | X1632 (G) | CE1176 (C) | 246F3 (AC) | CH119 (BC) | CE0217 (C) | CNE55 (AE) | BG505. T332N (A) | SF162 (B) | MLV (control) |
| BG505 E2p SApNP | Aluminum phosphate | D-S19 G3-M2 | >300 | >300 | 254 | >300 | >300 | >300 | >300 | >300 | >300 | >300 | >300 | >300 | 39 | 14 | >300 |
| BG505 I3-01v9 SApNP | Aluminum phosphate | D-S20 G3-M1 | >300 | >300 | 115 | >300 | >300 | 255 | >300 | >300 | >300 | >300 | >300 | >300 | 9 | 2 | >300 |
| | Aluminum hydroxide | D-S20 G4-M6 | >300 | >300 | 197 | >300 | >300 | >300 | >300 | 221 | >300 | >300 | >300 | >300 | 16 | 12 | >300 |

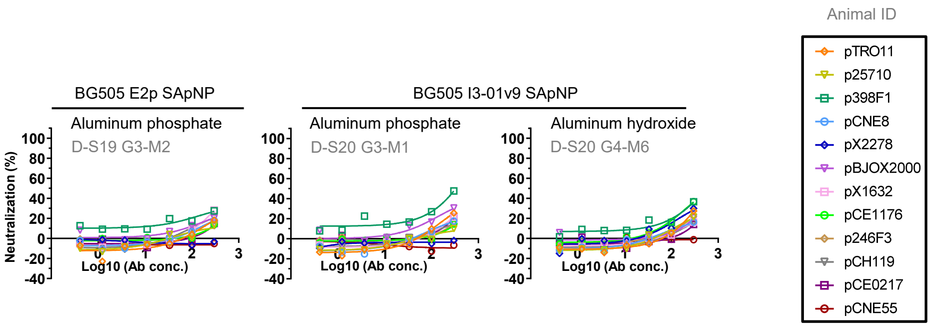

fig. S5

D Weeks 0-28 rabbit plasma neutralization against HIV BG505.T332N, $ID_{50}$ and $ID_{30}$ titers

| Vaccine | Adjuvant | Animal ID | $ID_{50}$ titers | | | | | | | | | | | |
|---|---|---|---|---|---|---|---|---|---|---|---|---|---|---|
| | | | w0 | w2 | w4 | w6 | w8 | w10 | w12 | w16 | w20 | w24 | w26 | w28 |
| BG505 UFO trimer | Addavax | 119 | <100 | <100 | <100 | <100 | <100 | <100 | <100 | <100 | <100 | <100 | <100 | <100 |
| | | 120 | <100 | <100 | <100 | <100 | <100 | <100 | <100 | <100 | <100 | <100 | <100 | <100 |
| | | 121 | <100 | <100 | <100 | <100 | <100 | 136 | <100 | <100 | <100 | <100 | 191 | <100 |
| | | 122 | <100 | <100 | <100 | <100 | <100 | 116 | <100 | <100 | <100 | <100 | 381 | 139 |
| | | 123 | <100 | <100 | <100 | <100 | <100 | <100 | <100 | <100 | <100 | <100 | 150 | <100 |
| | | 124 | <100 | <100 | <100 | 501 | 134 | 178 | <100 | <100 | <100 | <100 | 996 | 194 |
| BG505 FR SApNP | Addavax | 125 | <100 | <100 | <100 | <100 | <100 | <100 | <100 | <100 | <100 | <100 | <100 | <100 |
| | | 126 | <100 | <100 | <100 | <100 | <100 | 737 | 485 | 179 | 113 | <100 | 688 | 745 |
| | | 127 | <100 | <100 | <100 | <100 | <100 | <100 | <100 | <100 | <100 | <100 | 601 | 511 |
| | | 128 | <100 | <100 | <100 | <100 | <100 | <100 | <100 | <100 | <100 | <100 | <100 | <100 |
| | | 129 | <100 | <100 | <100 | <100 | <100 | <100 | <100 | <100 | <100 | <100 | <100 | <100 |
| | | 130 | <100 | <100 | <100 | <100 | <100 | 105 | <100 | <100 | <100 | <100 | 760 | 454 |
| Env Mix FR SApNP | Addavax | 131 | <100 | <100 | <100 | <100 | <100 | <100 | <100 | <100 | <100 | <100 | <100 | <100 |
| | | 132 | <100 | <100 | <100 | <100 | <100 | <100 | <100 | <100 | <100 | <100 | <100 | 113 |
| | | 133 | <100 | <100 | <100 | <100 | <100 | <100 | <100 | <100 | <100 | <100 | <100 | <100 |
| | | 134 | <100 | <100 | <100 | <100 | <100 | <100 | <100 | <100 | <100 | <100 | 151 | 125 |
| | | 135 | <100 | <100 | <100 | <100 | <100 | <100 | <100 | <100 | <100 | <100 | <100 | <100 |
| | | 136 | <100 | <100 | <100 | <100 | <100 | <100 | <100 | <100 | <100 | <100 | <100 | <100 |
| BG505 E2p SApNP | Addavax | 155 | <100 | <100 | <100 | <100 | <100 | 269 | 126 | <100 | <100 | <100 | 563 | 331 |
| | | 156 | <100 | <100 | <100 | <100 | <100 | <100 | <100 | <100 | <100 | <100 | <100 | <100 |
| | | 157 | <100 | <100 | <100 | <100 | <100 | <100 | <100 | <100 | <100 | <100 | <100 | <100 |
| | | 158 | <100 | <100 | <100 | <100 | <100 | <100 | <100 | <100 | <100 | <100 | <100 | <100 |
| | | 159 | <100 | <100 | <100 | <100 | <100 | <100 | <100 | <100 | <100 | <100 | <100 | <100 |
| | | 160 | <100 | <100 | <100 | <100 | <100 | 521 | 208 | <100 | <100 | <100 | 587 | 364 |
| Env Mix E2p SApNP | Addavax | 137 | <100 | <100 | <100 | <100 | <100 | <100 | <100 | <100 | <100 | <100 | <100 | <100 |
| | | 138 | <100 | <100 | <100 | <100 | <100 | <100 | <100 | <100 | <100 | <100 | <100 | <100 |
| | | 139 | <100 | <100 | <100 | <100 | <100 | <100 | <100 | <100 | <100 | <100 | <100 | <100 |
| | | 140 | <100 | <100 | <100 | 220 | <100 | <100 | <100 | <100 | <100 | <100 | 210 | <100 |
| | | 141 | <100 | <100 | <100 | <100 | <100 | <100 | <100 | <100 | <100 | <100 | 247 | 132 |
| | | 142 | <100 | <100 | <100 | <100 | <100 | <100 | <100 | <100 | <100 | <100 | <100 | <100 |
| BG505 I3-01v9 SApNP | Aluminum phosphate | 143 | <100 | <100 | <100 | <100 | <100 | 164 | 240 | <100 | <100 | <100 | 659 | 492 |
| | | 144 | <100 | <100 | <100 | <100 | <100 | <100 | <100 | <100 | <100 | <100 | <100 | 153 |
| | | 145 | <100 | <100 | <100 | <100 | <100 | <100 | <100 | <100 | <100 | <100 | 124 | <100 |
| | | 146 | <100 | <100 | <100 | <100 | <100 | <100 | <100 | <100 | <100 | <100 | 247 | 116 |
| | | 147 | <100 | <100 | <100 | <100 | <100 | 439 | 220 | 141 | 120 | 130 | 958 | 807 |
| | | 148 | <100 | <100 | <100 | <100 | <100 | <100 | <100 | <100 | <100 | <100 | <100 | <100 |
| Env Mix I3-01v9 SApNP | Aluminum phosphate | 149 | <100 | <100 | <100 | <100 | <100 | 402 | 252 | <100 | <100 | <100 | 334 | 382 |
| | | 150 | <100 | <100 | <100 | <100 | <100 | <100 | <100 | <100 | <100 | <100 | 271 | 155 |
| | | 151 | <100 | <100 | <100 | <100 | <100 | 656 | 330 | 241 | 142 | 111 | 1170 | 813 |
| | | 152 | <100 | <100 | <100 | <100 | <100 | <100 | <100 | <100 | <100 | <100 | 128 | <100 |
| | | 153 | <100 | <100 | <100 | <100 | <100 | <100 | <100 | <100 | <100 | <100 | <100 | <100 |
| | | 154 | <100 | <100 | <100 | <100 | <100 | <100 | <100 | <100 | <100 | <100 | <100 | <100 |

| Vaccine | Adjuvant | Animal ID | $ID_{30}$ titers | | | | | | | | | | | |
|---|---|---|---|---|---|---|---|---|---|---|---|---|---|---|
| | | | w0 | w2 | w4 | w6 | w8 | w10 | w12 | w16 | w20 | w24 | w26 | w28 |
| BG505 UFO trimer | Addavax | 119 | <100 | <100 | <100 | <100 | <100 | <100 | <100 | <100 | <100 | <100 | 120 | <100 |
| | | 120 | <100 | <100 | <100 | <100 | <100 | <100 | <100 | <100 | <100 | <100 | <100 | <100 |
| | | 121 | <100 | <100 | <100 | <100 | <100 | 316 | 185 | <100 | <100 | <100 | 446 | 179 |
| | | 122 | <100 | <100 | <100 | <100 | <100 | 270 | 164 | <100 | <100 | <100 | 889 | 324 |
| | | 123 | <100 | <100 | <100 | <100 | <100 | <100 | <100 | <100 | <100 | <100 | 351 | 197 |
| | | 124 | <100 | <100 | <100 | 1168 | 312 | 415 | 213 | 133 | <100 | <100 | 2324 | 454 |
| BG505 FR SApNP | Addavax | 125 | <100 | <100 | <100 | <100 | <100 | <100 | <100 | <100 | <100 | <100 | <100 | 210 |
| | | 126 | <100 | <100 | <100 | <100 | <100 | 1720 | 1131 | 417 | 263 | 140 | 1605 | 1738 |
| | | 127 | <100 | <100 | <100 | <100 | <100 | <100 | <100 | <100 | <100 | <100 | 1401 | 1193 |
| | | 128 | <100 | <100 | <100 | <100 | <100 | 109 | <100 | <100 | <100 | <100 | 186 | 140 |
| | | 129 | <100 | <100 | <100 | <100 | <100 | <100 | <100 | <100 | <100 | <100 | <100 | 124 |
| | | 130 | <100 | <100 | <100 | <100 | <100 | 245 | <100 | <100 | <100 | <100 | 1774 | 1060 |
| Env Mix FR SApNP | Addavax | 131 | <100 | <100 | <100 | <100 | <100 | <100 | <100 | <100 | <100 | <100 | <100 | 115 |
| | | 132 | <100 | <100 | <100 | <100 | <100 | <100 | <100 | <100 | <100 | <100 | 105 | 265 |
| | | 133 | <100 | <100 | <100 | <100 | <100 | <100 | <100 | <100 | <100 | <100 | <100 | <100 |
| | | 134 | <100 | <100 | <100 | <100 | <100 | <100 | <100 | <100 | <100 | <100 | 351 | 291 |
| | | 135 | <100 | <100 | <100 | <100 | <100 | <100 | <100 | <100 | <100 | <100 | <100 | <100 |
| | | 136 | <100 | <100 | <100 | <100 | <100 | <100 | <100 | <100 | <100 | <100 | <100 | 190 |
| BG505 E2p SApNP | Addavax | 155 | <100 | <100 | <100 | <100 | <100 | 627 | 293 | 212 | 147 | 115 | 1313 | 773 |
| | | 156 | <100 | <100 | <100 | <100 | <100 | 122 | <100 | <100 | <100 | <100 | 154 | <100 |
| | | 157 | <100 | <100 | <100 | <100 | <100 | 175 | <100 | <100 | 133 | <100 | 107 | <100 |
| | | 158 | <100 | <100 | <100 | <100 | <100 | 170 | <100 | <100 | <100 | <100 | 149 | <100 |
| | | 159 | <100 | <100 | <100 | <100 | <100 | <100 | <100 | 170 | <100 | <100 | 198 | <100 |
| | | 160 | <100 | <100 | <100 | <100 | <100 | 1215 | 485 | <100 | <100 | 109 | 1369 | 849 |
| Env Mix E2p SApNP | Addavax | 137 | <100 | <100 | <100 | <100 | <100 | <100 | <100 | <100 | <100 | <100 | <100 | <100 |
| | | 138 | <100 | <100 | <100 | <100 | <100 | 117 | <100 | <100 | <100 | <100 | <100 | 116 |
| | | 139 | <100 | <100 | <100 | <100 | <100 | <100 | <100 | <100 | <100 | <100 | <100 | 113 |
| | | 140 | <100 | <100 | <100 | 513 | <100 | <100 | <100 | <100 | <100 | <100 | 490 | 180 |
| | | 141 | <100 | <100 | <100 | <100 | <100 | 118 | <100 | <100 | <100 | <100 | 576 | 308 |
| | | 142 | <100 | <100 | <100 | <100 | <100 | <100 | <100 | <100 | <100 | <100 | <100 | 141 |
| BG505 I3-01v9 SApNP | Aluminum phosphate | 143 | <100 | <100 | <100 | <100 | <100 | 382 | 561 | 198 | 208 | 114 | 1537 | 1148 |
| | | 144 | <100 | <100 | <100 | <100 | <100 | <100 | <100 | <100 | <100 | <100 | <100 | 357 |
| | | 145 | <100 | <100 | <100 | <100 | <100 | <100 | <100 | <100 | <100 | <100 | 289 | 149 |
| | | 146 | <100 | <100 | <100 | <100 | <100 | 104 | <100 | <100 | <100 | <100 | 576 | 271 |
| | | 147 | <100 | <100 | <100 | <100 | <100 | 1025 | 514 | 328 | 280 | 302 | 2235 | 1883 |
| | | 148 | <100 | <100 | <100 | <100 | <100 | <100 | <100 | <100 | <100 | <100 | 129 | 139 |
| Env Mix I3-01v9 SApNP | Aluminum phosphate | 149 | <100 | <100 | <100 | <100 | 106 | 939 | 589 | 218 | 161 | 157 | 780 | 892 |
| | | 150 | <100 | <100 | <100 | <100 | <100 | 123 | <100 | <100 | <100 | <100 | 631 | 363 |
| | | 151 | <100 | <100 | <100 | <100 | <100 | 1530 | 770 | 562 | 331 | 260 | 2731 | 1897 |
| | | 152 | <100 | <100 | <100 | <100 | <100 | 164 | <100 | <100 | <100 | <100 | 298 | 122 |
| | | 153 | <100 | <100 | <100 | <100 | <100 | <100 | <100 | <100 | <100 | <100 | <100 | <100 |
| | | 154 | <100 | <100 | <100 | <100 | <100 | 114 | <100 | <100 | <100 | <100 | 145 | <100 |

fig. S5

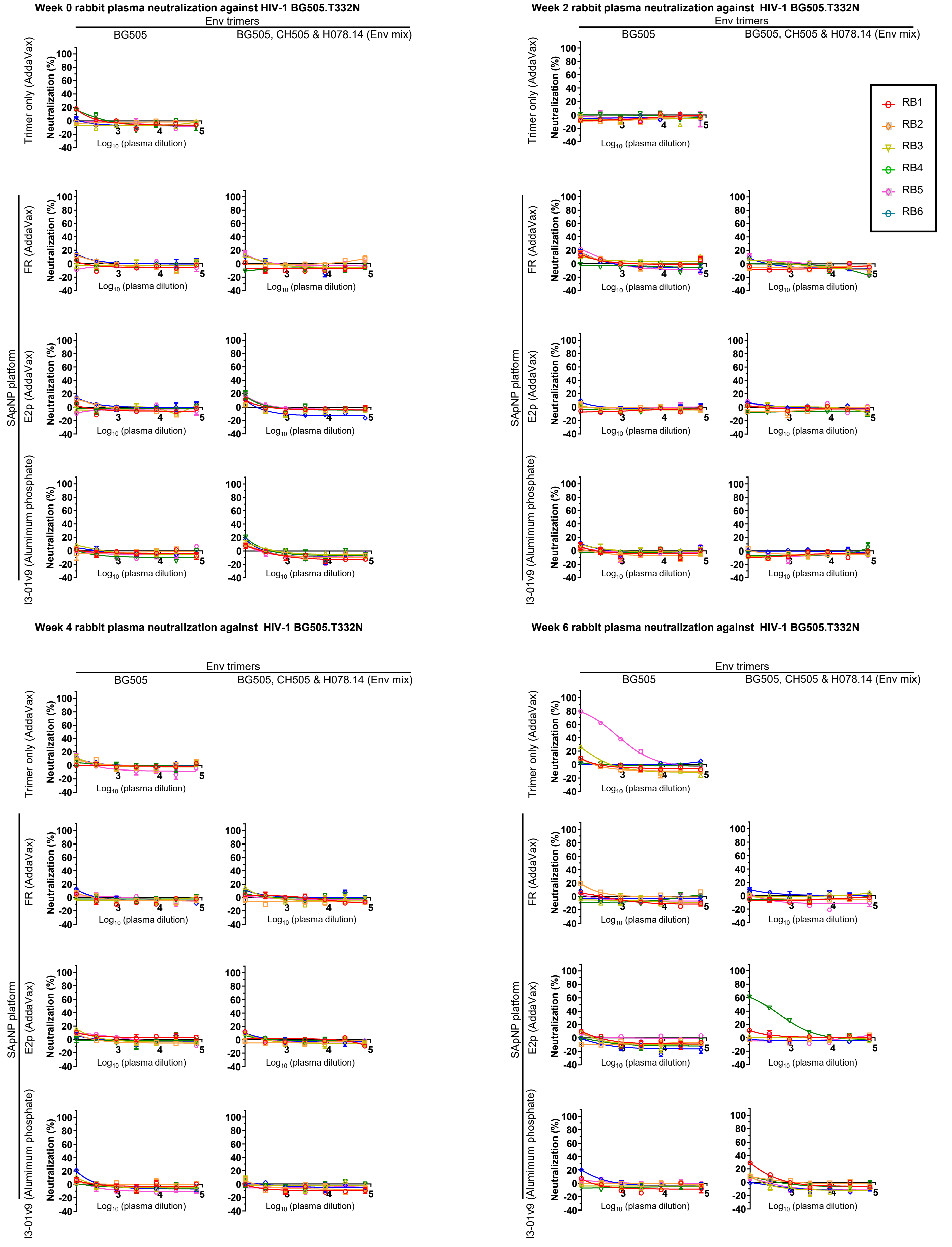

Week 0 rabbit plasma neutralization against HIV-1 BG505.T332N
Week 2 rabbit plasma neutralization against HIV-1 BG505.T332N
Week 4 rabbit plasma neutralization against HIV-1 BG505.T332N
Week 6 rabbit plasma neutralization against HIV-1 BG505.T332N
Env trimers
BG505
BG505, CH505 & H078.14 (Env mix)
Trimer only (AddaVax)
SApNP platform
FR (AddaVax)
E2p (AddaVax)
I3-01v9 (Alumimum phosphate)
Neutralization (%)
Log10 (plasma dilution)
RB1
RB2
RB3
RB4
RB5
RB6

fig. S5

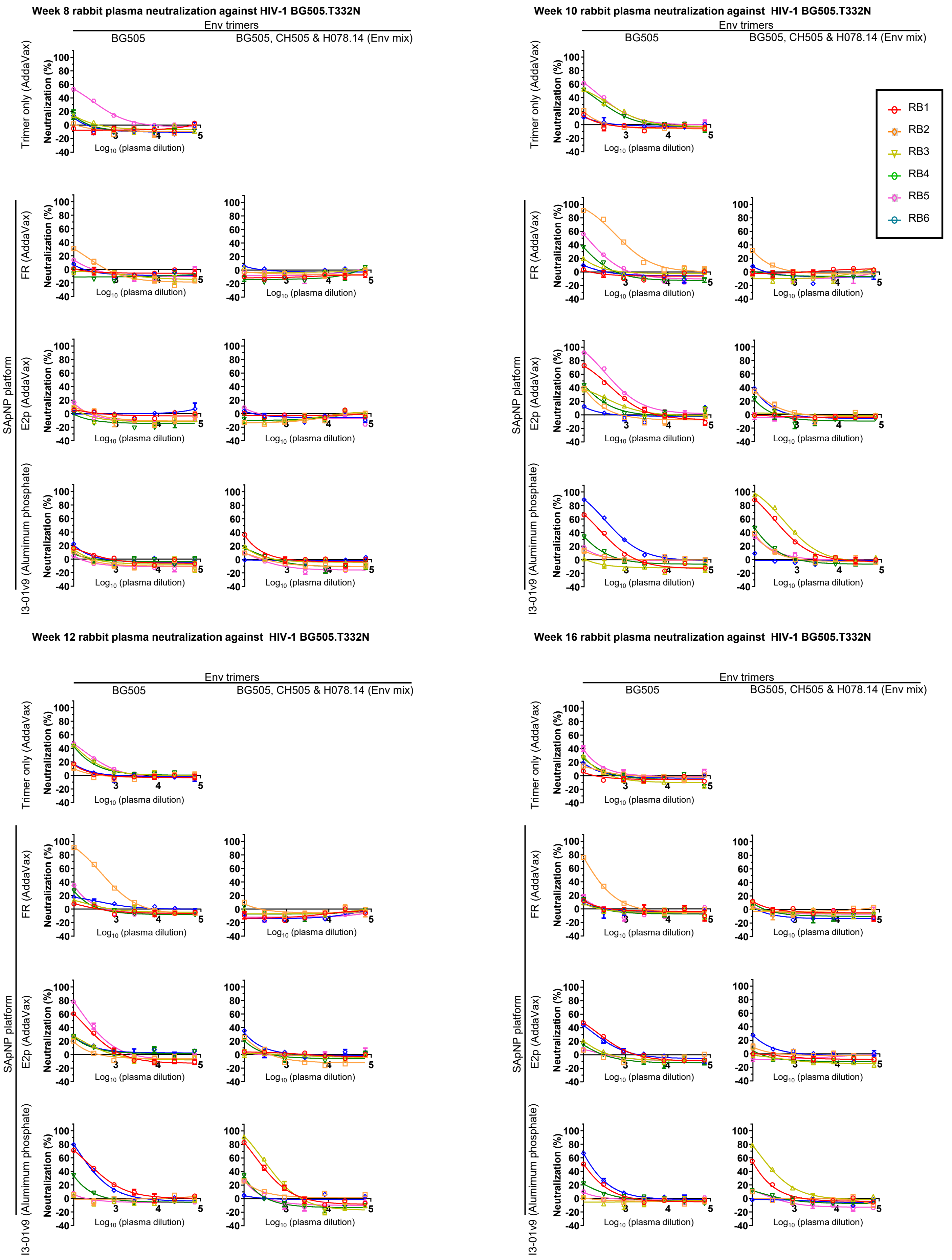
Week 8 rabbit plasma neutralization against HIV-1 BG505.T332N
Week 10 rabbit plasma neutralization against HIV-1 BG505.T332N
Week 12 rabbit plasma neutralization against HIV-1 BG505.T332N
Week 16 rabbit plasma neutralization against HIV-1 BG505.T332N
Env trimers
BG505
BG505, CH505 & H078.14 (Env mix)
Trimer only (AddaVax)
SApNP platform
FR (AddaVax)
E2p (AddaVax)
I3-01v9 (Alumimum phosphate)
Neutralization (%)
Log10 (plasma dilution)
RB1
RB2
RB3
RB4
RB5
RB6

fig. S5

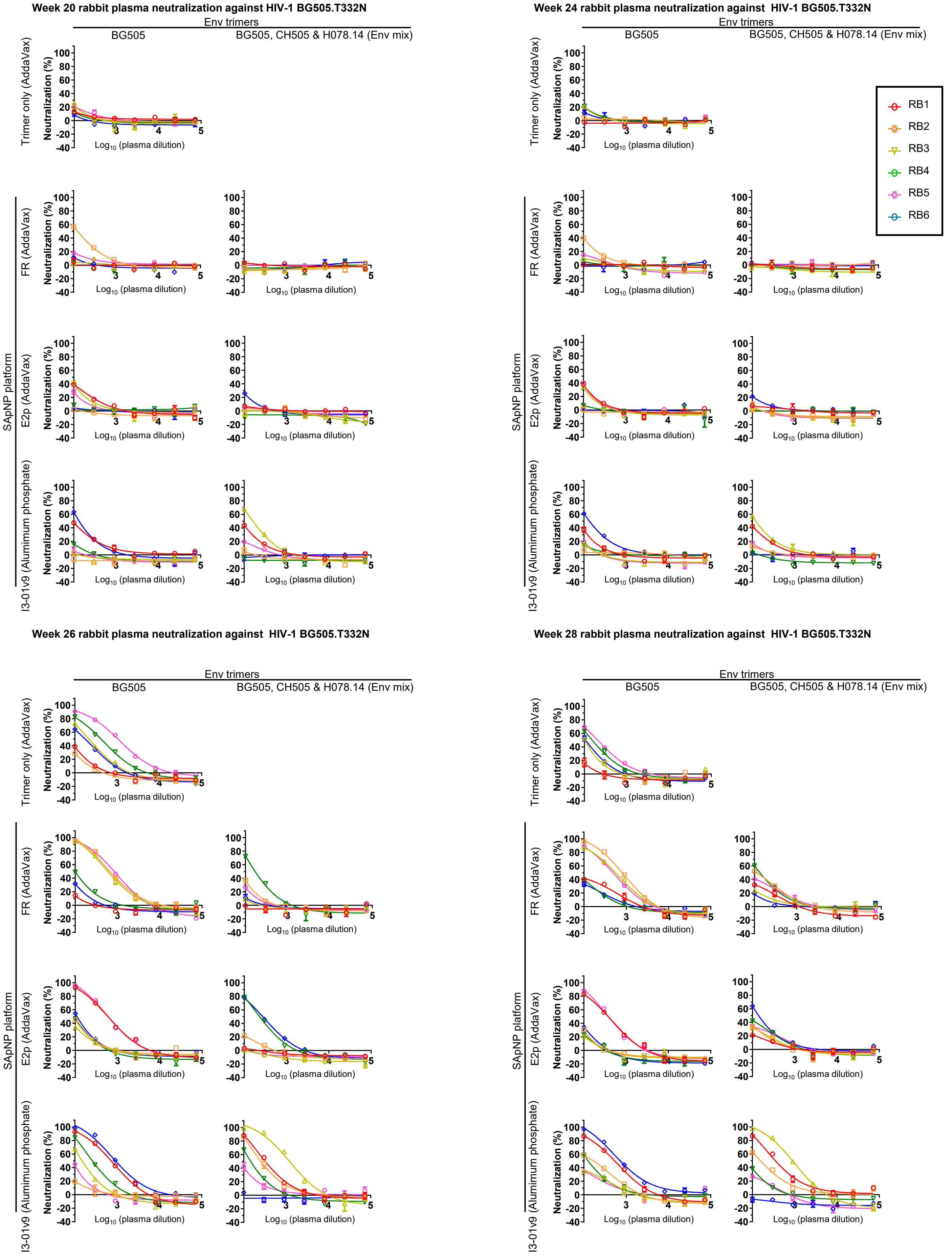

Week 20 rabbit plasma neutralization against HIV-1 BG505.T332N
Week 24 rabbit plasma neutralization against HIV-1 BG505.T332N
Week 26 rabbit plasma neutralization against HIV-1 BG505.T332N
Week 28 rabbit plasma neutralization against HIV-1 BG505.T332N
Env trimers
BG505
BG505, CH505 & H078.14 (Env mix)
Trimer only (AddaVax)
SApNP platform
FR (AddaVax)
E2p (AddaVax)
I3-01v9 (Alumimum phosphate)
Neutralization (%)
Log10 (plasma dilution)
RB1
RB2
RB3
RB4
RB5
RB6

**fig. S5**

E **Weeks 0-28 rabbit plasma neutralization against HIV SF162, $ID_{50}$ titers**

| Vaccine | Adjuvant | Animal ID | $ID_{50}$ titers | | | | | | | | | | | |
|---|---|---|---|---|---|---|---|---|---|---|---|---|---|---|
| | | | w0 | w2 | w4 | w6 | w8 | w10 | w12 | w16 | w20 | w24 | w26 | w28 |
| BG505 UFO trimer | Addavax | 119 | <100 | <100 | <100 | <100 | <100 | 3429 | 920 | 375 | <100 | <100 | 1953 | 565 |
| | | 120 | <100 | <100 | <100 | <100 | <100 | 310 | 107 | <100 | <100 | <100 | 1120 | 452 |
| | | 121 | <100 | <100 | <100 | <100 | <100 | 1926 | 821 | 362 | 108 | 101 | 1190 | 627 |
| | | 122 | <100 | <100 | <100 | <100 | <100 | 840 | 174 | <100 | <100 | <100 | 938 | 280 |
| | | 123 | <100 | <100 | <100 | <100 | <100 | 661 | 135 | <100 | <100 | <100 | 673 | 313 |
| | | 124 | <100 | <100 | <100 | 405 | <100 | 641 | 153 | <100 | <100 | <100 | 2015 | 689 |
| BG505 FR SApNP | Addavax | 125 | <100 | <100 | <100 | <100 | <100 | 2056 | 894 | 423 | <100 | <100 | 3716 | 882 |
| | | 126 | <100 | <100 | 102 | 219 | 164 | 2745 | 975 | 425 | 115 | <100 | 1534 | 754 |
| | | 127 | <100 | <100 | <100 | 200 | 362 | 4060 | 1343 | 554 | 240 | 236 | 5866 | 1759 |
| | | 128 | <100 | <100 | <100 | <100 | 113 | 4055 | 1069 | 265 | 129 | 103 | 4408 | 968 |
| | | 129 | <100 | <100 | <100 | <100 | <100 | 1553 | 521 | 228 | <100 | 133 | 2736 | 829 |
| | | 130 | <100 | <100 | <100 | <100 | <100 | 927 | 256 | 106 | <100 | <100 | 2552 | 973 |
| Env Mix FR SApNP | Addavax | 131 | <100 | <100 | <100 | <100 | 359 | 1025 | 284 | 191 | <100 | <100 | 1541 | 562 |
| | | 132 | <100 | <100 | <100 | 103 | 111 | 1222 | 485 | 218 | <100 | <100 | 1534 | 210 |
| | | 133 | <100 | <100 | <100 | <100 | <100 | 566 | 180 | 104 | <100 | <100 | 1273 | 355 |
| | | 134 | <100 | <100 | <100 | 123 | 346 | 2494 | 1150 | 665 | 214 | 221 | 4272 | 641 |
| | | 135 | <100 | <100 | <100 | 129 | 215 | 2301 | 706 | 252 | <100 | 122 | 3695 | 515 |
| | | 136 | <100 | <100 | <100 | <100 | <100 | 942 | 272 | 152 | <100 | <100 | 4795 | 225 |
| BG505 E2p SApNP | Addavax | 155 | <100 | <100 | <100 | <100 | <100 | 474 | 162 | <100 | <100 | <100 | 1424 | 606 |
| | | 156 | <100 | <100 | <100 | <100 | <100 | 1003 | 323 | <100 | <100 | <100 | 1833 | 257 |
| | | 157 | <100 | <100 | <100 | <100 | <100 | 5656 | 1819 | 390 | 294 | 190 | 2726 | 820 |
| | | 158 | <100 | <100 | <100 | <100 | <100 | 1405 | 428 | <100 | <100 | <100 | 866 | 317 |
| | | 159 | <100 | <100 | <100 | <100 | <100 | 2241 | 666 | 106 | 149 | 209 | 2712 | 1131 |
| | | 160 | <100 | <100 | <100 | <100 | <100 | 1063 | 311 | 183 | <100 | 108 | 610 | 293 |
| Env Mix E2p SApNP | Addavax | 137 | <100 | <100 | <100 | <100 | <100 | 912 | 329 | 167 | <100 | 107 | 1061 | 275 |
| | | 138 | <100 | <100 | <100 | <100 | <100 | 1220 | 360 | 115 | <100 | <100 | 3029 | 660 |
| | | 139 | <100 | <100 | <100 | 202 | 309 | 1557 | 1018 | 643 | 395 | 334 | 7500 | 946 |
| | | 140 | <100 | <100 | <100 | 138 | <100 | 2666 | 833 | 280 | <100 | <100 | 3422 | 517 |
| | | 141 | <100 | <100 | <100 | <100 | <100 | 1484 | 461 | 234 | <100 | <100 | 895 | 230 |
| | | 142 | <100 | <100 | <100 | <100 | <100 | 1949 | 836 | 467 | 230 | 244 | 3779 | 993 |
| BG505 I3-01v9 SApNP | Aluminum phosphate | 143 | <100 | <100 | <100 | <100 | 132 | 1101 | 342 | 173 | <100 | <100 | 2028 | 216 |
| | | 144 | <100 | <100 | <100 | 146 | 317 | 1641 | 833 | 464 | 297 | 359 | 5417 | 1031 |
| | | 145 | <100 | <100 | 112 | 121 | 174 | 1084 | 415 | 223 | 114 | 139 | 5341 | 1272 |
| | | 146 | <100 | <100 | <100 | 102 | 262 | 1469 | 1250 | 693 | 233 | 245 | 2477 | 579 |
| | | 147 | <100 | <100 | 287 | 1378 | 2413 | 3366 | 2188 | 1287 | 895 | 924 | 6167 | 1551 |
| | | 148 | <100 | <100 | <100 | 255 | 532 | 3835 | 1642 | 717 | 320 | 256 | 2790 | 514 |
| Env Mix I3-01v9 SApNP | Aluminum phosphate | 149 | <100 | <100 | <100 | 291 | 449 | 996 | 418 | 124 | <100 | 104 | 1396 | 363 |
| | | 150 | <100 | <100 | <100 | 210 | 370 | 2981 | 2027 | 275 | 147 | 158 | 5805 | 1113 |
| | | 151 | <100 | <100 | <100 | <100 | 179 | 6838 | 1663 | 566 | 355 | 325 | 7193 | 3074 |
| | | 152 | <100 | <100 | <100 | 220 | 276 | 2487 | 726 | 230 | 110 | 114 | 5597 | 1552 |
| | | 153 | <100 | <100 | 116 | 333 | 349 | 4624 | 1252 | 392 | 159 | 171 | 7687 | 3191 |
| | | 154 | <100 | <100 | <100 | 145 | 223 | 2588 | 737 | 250 | 163 | 215 | 3045 | 844 |

fig. S5

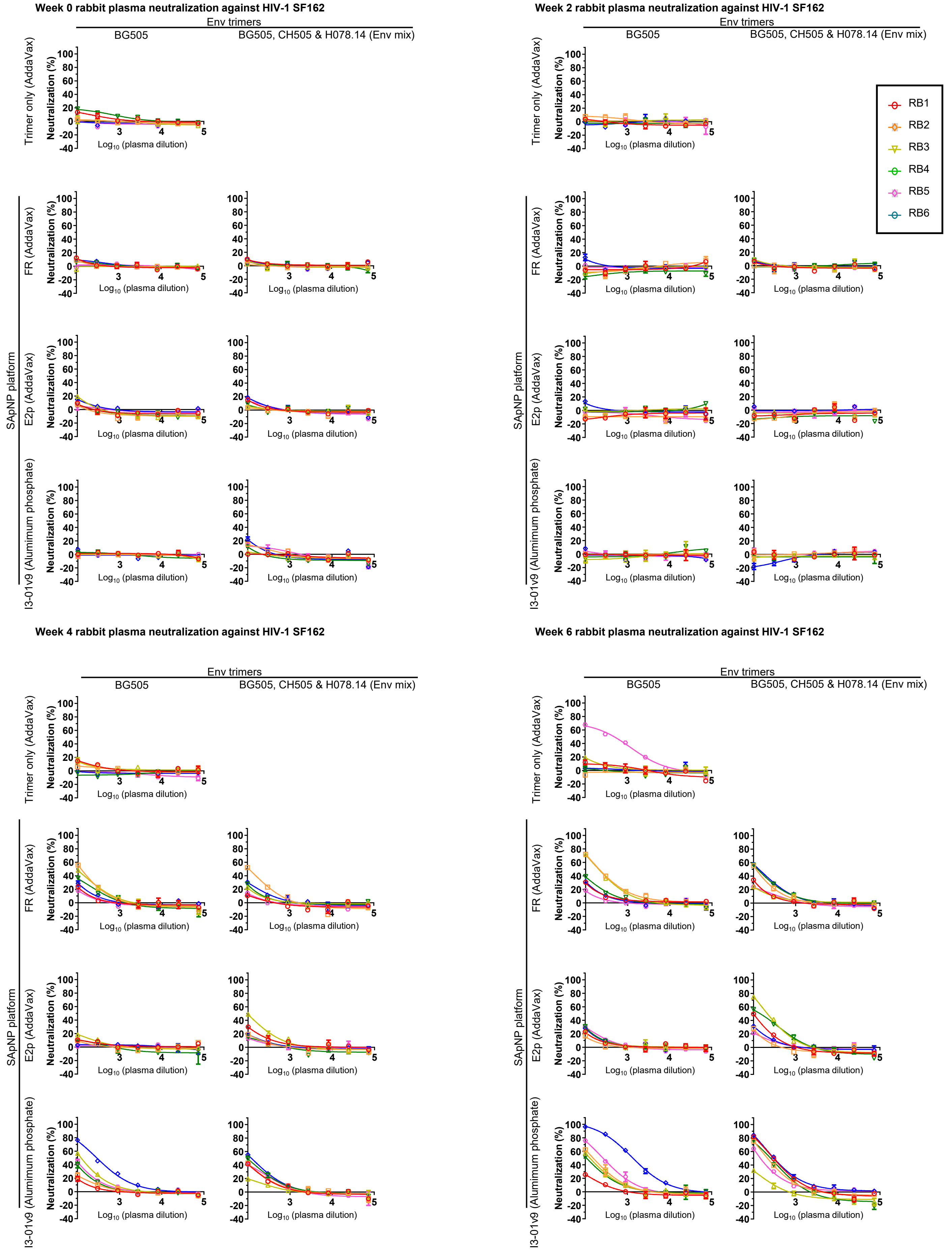

Week 0 rabbit plasma neutralization against HIV-1 SF162
Week 2 rabbit plasma neutralization against HIV-1 SF162
Week 4 rabbit plasma neutralization against HIV-1 SF162
Week 6 rabbit plasma neutralization against HIV-1 SF162
Env trimers
BG505
BG505, CH505 & H078.14 (Env mix)
Trimer only (AddaVax)
SApNP platform
FR (AddaVax)
E2p (AddaVax)
I3-01v9 (Alumimum phosphate)
Neutralization (%)
Log10 (plasma dilution)
RB1
RB2
RB3
RB4
RB5
RB6

#### fig. S5

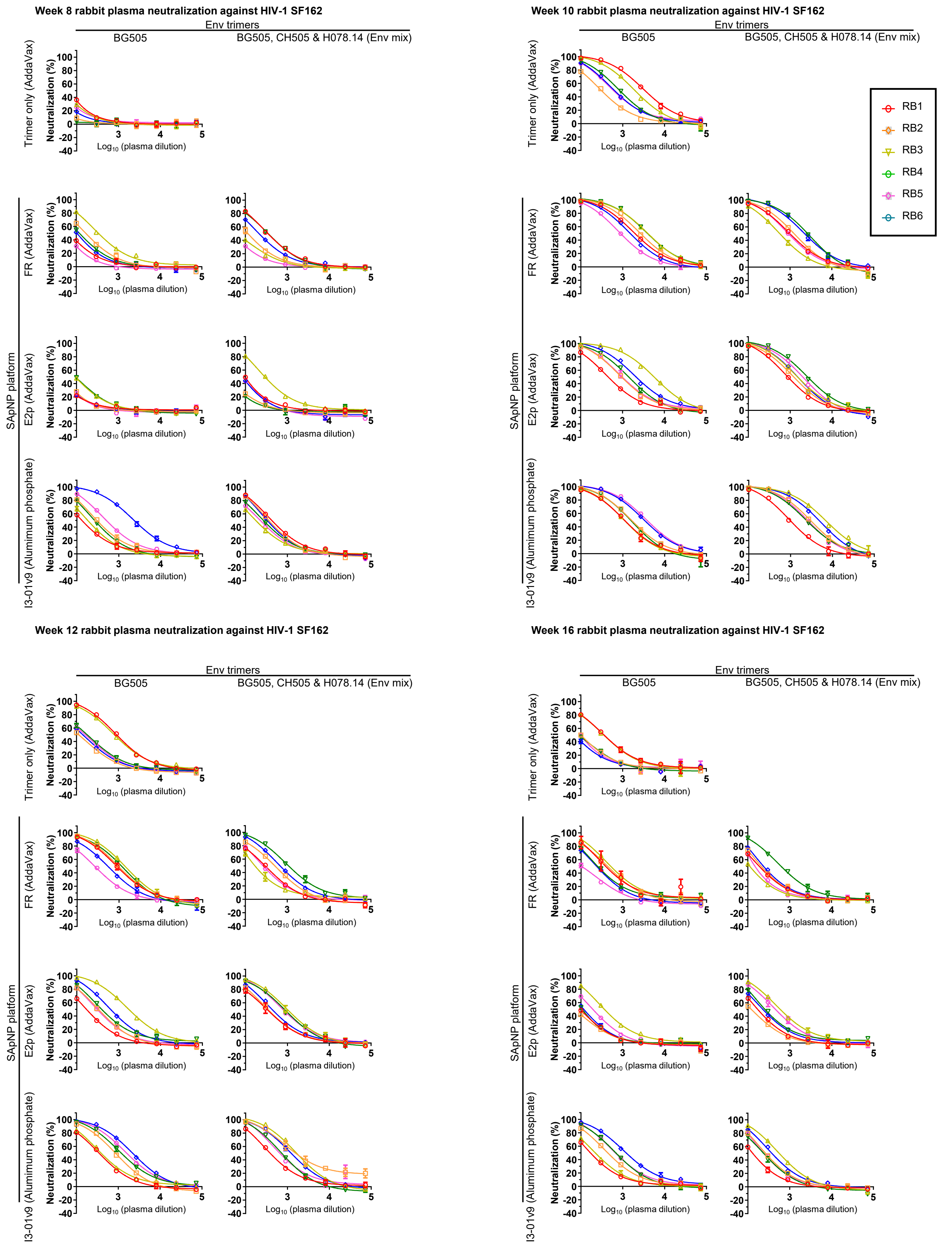

Week 8 rabbit plasma neutralization against HIV-1 SF162
Week 10 rabbit plasma neutralization against HIV-1 SF162
Week 12 rabbit plasma neutralization against HIV-1 SF162
Week 16 rabbit plasma neutralization against HIV-1 SF162
Env trimers
BG505
BG505, CH505 & H078.14 (Env mix)
Trimer only (AddaVax)
SApNP platform
FR (AddaVax)
E2p (AddaVax)
I3-01v9 (Alumimum phosphate)
I3-01v9 (Aluminum phosphate)
Neutralization (%)
$Log_{10}$ (plasma dilution)
RB1
RB2
RB3
RB4
RB5
RB6

fig. S5

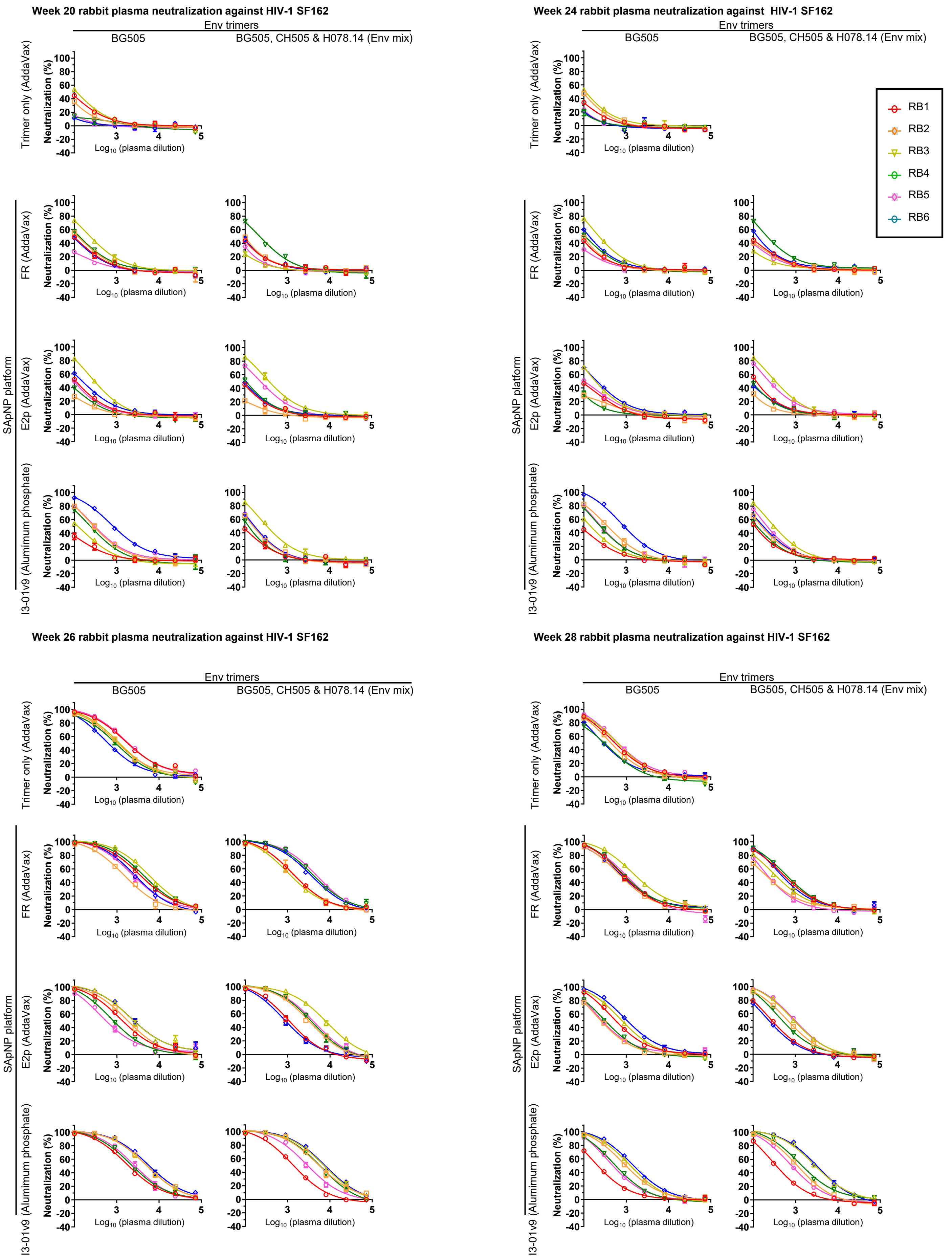
Week 20 rabbit plasma neutralization against HIV-1 SF162
Week 24 rabbit plasma neutralization against HIV-1 SF162
Week 26 rabbit plasma neutralization against HIV-1 SF162
Week 28 rabbit plasma neutralization against HIV-1 SF162
Env trimers
BG505
BG505, CH505 & H078.14 (Env mix)
Trimer only (AddaVax)
SApNP platform
FR (AddaVax)
E2p (AddaVax)
I3-01v9 (Alumimum phosphate)
Neutralization (%)
Log10 (plasma dilution)
RB1
RB2
RB3
RB4
RB5
RB6

fig. S5

F Weeks 0, 2, 10 & 26 rabbit plasma neutralization against MLV, $ID_{50}$ titers

| Vaccine | Adjuvant | Animal ID | $ID_{50}$ titers | | | |
|---|---|---|---|---|---|---|
| | | | w0 | w2 | w10 | w26 |
| BG505 UFO trimer | Addavax | 119 | <100 | <100 | <100 | <100 |
| | | 120 | <100 | <100 | <100 | <100 |
| | | 121 | <100 | <100 | <100 | <100 |
| | | 122 | <100 | <100 | <100 | <100 |
| | | 123 | <100 | <100 | <100 | <100 |
| | | 124 | <100 | <100 | <100 | <100 |
| BG505 FR SApNP | Addavax | 125 | <100 | <100 | <100 | <100 |
| | | 126 | <100 | <100 | <100 | <100 |
| | | 127 | <100 | <100 | <100 | <100 |
| | | 128 | <100 | <100 | <100 | <100 |
| | | 129 | <100 | <100 | <100 | <100 |
| | | 130 | <100 | <100 | <100 | <100 |
| Env Mix FR SApNP | Addavax | 131 | <100 | <100 | <100 | <100 |
| | | 132 | <100 | <100 | <100 | <100 |
| | | 133 | <100 | <100 | <100 | <100 |
| | | 134 | <100 | <100 | <100 | <100 |
| | | 135 | <100 | <100 | <100 | <100 |
| | | 136 | <100 | <100 | <100 | <100 |
| BG505 E2p SApNP | Addavax | 155 | <100 | <100 | <100 | <100 |
| | | 156 | <100 | <100 | <100 | <100 |
| | | 157 | <100 | <100 | <100 | <100 |
| | | 158 | <100 | <100 | <100 | <100 |
| | | 159 | <100 | <100 | <100 | <100 |
| | | 160 | <100 | <100 | <100 | <100 |
| Env Mix E2p SApNP | Addavax | 137 | <100 | <100 | <100 | <100 |
| | | 138 | <100 | <100 | <100 | <100 |
| | | 139 | <100 | <100 | <100 | <100 |
| | | 140 | <100 | <100 | <100 | <100 |
| | | 141 | <100 | <100 | <100 | <100 |
| | | 142 | <100 | <100 | <100 | <100 |
| BG505 I3-01v9 SApNP | Aluminum phosphate | 143 | <100 | <100 | <100 | <100 |
| | | 144 | <100 | <100 | <100 | <100 |
| | | 145 | <100 | <100 | <100 | <100 |
| | | 146 | <100 | <100 | <100 | <100 |
| | | 147 | <100 | <100 | <100 | <100 |
| | | 148 | <100 | <100 | <100 | <100 |
| Env Mix I3-01v9 SApNP | Aluminum phosphate | 149 | <100 | <100 | <100 | <100 |
| | | 150 | <100 | <100 | <100 | <100 |
| | | 151 | <100 | <100 | <100 | <100 |
| | | 152 | <100 | <100 | <100 | <100 |
| | | 153 | <100 | <100 | <100 | <100 |
| | | 154 | <100 | <100 | <100 | <100 |

fig. S5

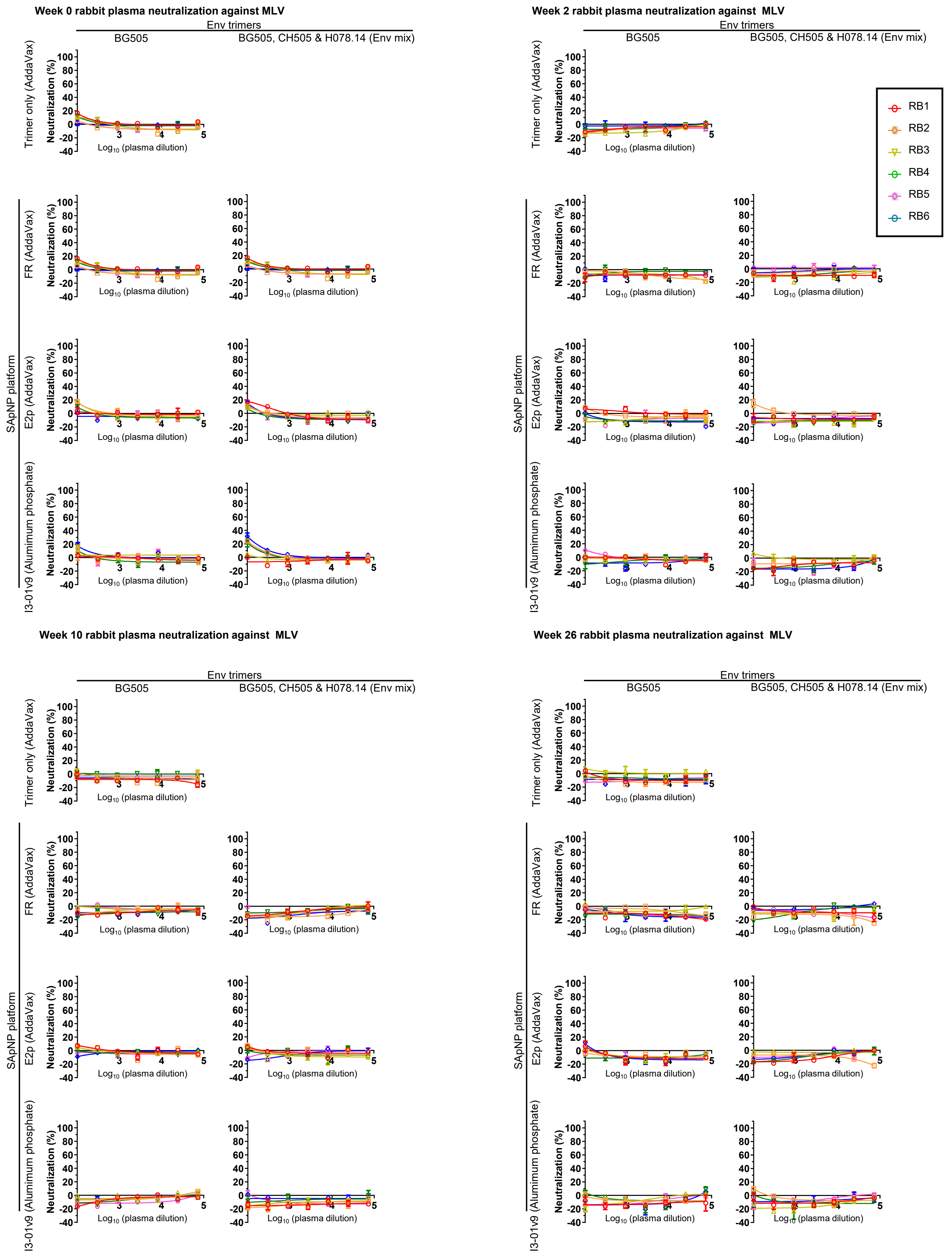
Week 0 rabbit plasma neutralization against MLV
Week 2 rabbit plasma neutralization against MLV
Week 10 rabbit plasma neutralization against MLV
Week 26 rabbit plasma neutralization against MLV
Env trimers
BG505
BG505, CH505 & H078.14 (Env mix)
Trimer only (AddaVax)
SApNP platform
FR (AddaVax)
E2p (AddaVax)
I3-01v9 (Alumimum phosphate)
Neutralization (%)
Log10 (plasma dilution)
RB1
RB2
RB3
RB4
RB5
RB6

**fig. S5**

**G** **Week 26 rabbit plasma neutralization against the 12-virus global panel, $ID_{50}$ and $ID_{30}$ titers**

| Vaccine | Adjuvant | Rabbit ID | $ID_{50}$ titers for each virus (clade) | | | | | | | | | | | | | | |
|---|---|---|---|---|---|---|---|---|---|---|---|---|---|---|---|
| | | | TRO11 (B) | 25710 (C) | 398F1 (A) | CNE8 (AE) | X2278 (B) | BJOX-2000 (BC) | X1632 (G) | CE1176 (C) | 246F3 (AC) | CH119 (BC) | CE0217 (C) | CNE55 (AE) | BG505.T332N (A) | SF162 (B) | MLV (control) |
| BG505 UFO trimer | Addavax | 119 | <40 | <40 | 135 | <40 | <40 | <40 | <40 | <40 | <40 | <40 | <40 | <40 | <100 | 1953 | <40 |
| | | 120 | <40 | <40 | 80 | <40 | <40 | <40 | <40 | <40 | <40 | <40 | <40 | <40 | <100 | 1120 | <40 |
| | | 121 | <40 | <40 | 93 | <40 | <40 | <40 | <40 | <40 | <40 | <40 | <40 | <40 | 191 | 1190 | <40 |
| | | 122 | <40 | <40 | 144 | <40 | <40 | <40 | <40 | <40 | <40 | <40 | <40 | <40 | 381 | 938 | <40 |
| | | 123 | 57 | 54 | 258 | 48 | <40 | <40 | <40 | <40 | 43 | <40 | <40 | <40 | 150 | 673 | <40 |
| | | 124 | <40 | <40 | 116 | <40 | <40 | <40 | <40 | <40 | <40 | <40 | <40 | <40 | 510 | 2015 | <40 |
| BG505 FR SApNP | Addavax | 125 | <40 | <40 | 221 | <40 | <40 | <40 | <40 | <40 | <40 | <40 | <40 | <40 | <100 | 3716 | <40 |
| | | 126 | <40 | <40 | 94 | <40 | <40 | <40 | <40 | <40 | <40 | <40 | <40 | <40 | 688 | 1534 | <40 |
| | | 127 | <40 | <40 | 127 | <40 | <40 | <40 | <40 | <40 | <40 | <40 | <40 | <40 | 601 | 5866 | <40 |
| | | 128 | <40 | <40 | 154 | <40 | <40 | <40 | <40 | <40 | <40 | <40 | <40 | <40 | <100 | 4408 | <40 |
| | | 129 | <40 | 40 | 148 | <40 | <40 | <40 | <40 | <40 | <40 | <40 | <40 | <40 | <100 | 2736 | <40 |
| | | 130 | <40 | <40 | 100 | <40 | <40 | <40 | <40 | <40 | <40 | <40 | <40 | <40 | 760 | 2552 | <40 |
| Env Mix FR SApNP | Addavax | 131 | <40 | 53 | 174 | <40 | <40 | <40 | <40 | <40 | <40 | <40 | <40 | <40 | <100 | 1541 | <40 |
| | | 132 | <40 | <40 | 106 | <40 | <40 | <40 | <40 | <40 | <40 | <40 | <40 | <40 | <100 | 1534 | <40 |
| | | 133 | <40 | 71 | 179 | 48 | <40 | 58 | <40 | 47 | 41 | <40 | <40 | <40 | <100 | 1273 | <40 |
| | | 134 | <40 | <40 | 74 | <40 | <40 | <40 | <40 | <40 | <40 | <40 | <40 | <40 | 151 | 4272 | <40 |
| | | 135 | <40 | <40 | 100 | <40 | <40 | <40 | <40 | <40 | <40 | <40 | <40 | <40 | <100 | 3695 | <40 |
| | | 136 | <40 | <40 | 106 | <40 | <40 | <40 | <40 | <40 | <40 | <40 | <40 | <40 | <100 | 4795 | <40 |
| BG505 E2p SApNP | Addavax | 155 | <40 | <40 | 63 | <40 | <40 | <40 | <40 | <40 | <40 | <40 | <40 | <40 | 563 | 1424 | <40 |
| | | 156 | <40 | <40 | 55 | <40 | <40 | <40 | <40 | <40 | <40 | <40 | <40 | <40 | <100 | 1833 | <40 |
| | | 157 | <40 | 42 | 108 | <40 | <40 | <40 | <40 | <40 | <40 | <40 | <40 | <40 | <100 | 2726 | <40 |
| | | 158 | <40 | <40 | 103 | <40 | <40 | <40 | <40 | <40 | <40 | <40 | <40 | <40 | <100 | 866 | <40 |
| | | 159 | 54 | <40 | 417 | 88 | <40 | <40 | <40 | 59 | 76 | 44 | <40 | 43 | <100 | 2712 | <40 |
| | | 160 | <40 | <40 | 167 | <40 | <40 | <40 | <40 | <40 | <40 | <40 | <40 | <40 | 587 | 610 | <40 |
| Env Mix E2p SApNP | Addavax | 137 | <40 | <40 | 157 | <40 | <40 | <40 | <40 | <40 | <40 | <40 | <40 | <40 | <100 | 1061 | <40 |
| | | 138 | <40 | 44 | 108 | <40 | <40 | <40 | <40 | <40 | <40 | <40 | <40 | <40 | <100 | 3029 | <40 |
| | | 139 | <40 | 58 | 386 | <40 | <40 | <40 | <40 | <40 | <40 | <40 | <40 | <40 | <100 | 7500 | <40 |
| | | 140 | <40 | <40 | 171 | <40 | <40 | <40 | <40 | <40 | <40 | <40 | <40 | <40 | <100 | 3422 | <40 |
| | | 141 | <40 | 44 | 202 | <40 | <40 | <40 | <40 | <40 | <40 | <40 | <40 | <40 | 247 | 895 | <40 |
| | | 142 | <40 | 51 | 228 | <40 | <40 | <40 | <40 | <40 | <40 | <40 | <40 | <40 | <100 | 3779 | <40 |
| BG505 I3-01v9 SApNP | Aluminum phosphate | 143 | <40 | 51 | 363 | <40 | <40 | <40 | <40 | <40 | <40 | <40 | <40 | <40 | 973 | 2028 | <40 |
| | | 144 | <40 | <40 | 72 | <40 | <40 | <40 | <40 | <40 | <40 | <40 | <40 | <40 | <100 | 5417 | <40 |
| | | 145 | <40 | <40 | 174 | <40 | <40 | <40 | <40 | <40 | <40 | <40 | <40 | <40 | <100 | 5341 | <40 |
| | | 146 | <40 | <40 | 179 | <40 | <40 | <40 | <40 | <40 | <40 | 42 | <40 | <40 | 247 | 2477 | <40 |
| | | 147 | <40 | 71 | 205 | 48 | 50 | 47 | <40 | <40 | 42 | <40 | <40 | <40 | 958 | 6167 | <40 |
| | | 148 | <40 | 80 | 172 | <40 | <40 | <40 | <40 | <40 | <40 | <40 | <40 | 43 | <100 | 2790 | <40 |
| Env Mix I3-01v9 SApNP | Aluminum phosphate | 149 | <40 | <40 | 257 | <40 | <40 | <40 | <40 | <40 | <40 | <40 | <40 | <40 | 334 | 1396 | <40 |
| | | 150 | <40 | 59 | 406 | 45 | <40 | <40 | <40 | <40 | <40 | <40 | <40 | <40 | 271 | 5805 | <40 |
| | | 151 | <40 | 53 | 259 | <40 | <40 | <40 | <40 | <40 | <40 | <40 | <40 | <40 | 1170 | 7193 | <40 |
| | | 152 | <40 | <40 | 178 | <40 | <40 | <40 | <40 | <40 | <40 | <40 | <40 | <40 | 128 | 5597 | <40 |
| | | 153 | <40 | 81 | 883 | <40 | <40 | <40 | <40 | <40 | <40 | 52 | <40 | <40 | <100 | 7687 | <40 |
| | | 154 | <40 | 61 | 225 | <40 | <40 | 42 | <40 | <40 | <40 | <40 | <40 | <40 | <100 | 3045 | <40 |

| Vaccine | Adjuvant | Rabbit ID | $ID_{30}$ titers for each virus (clade) | | | | | | | | | | | | | | |
|---|---|---|---|---|---|---|---|---|---|---|---|---|---|---|---|
| | | | TRO11 (B) | 25710 (C) | 398F1 (A) | CNE8 (AE) | X2278 (B) | BJOX-2000 (BC) | X1632 (G) | CE1176 (C) | 246F3 (AC) | CH119 (BC) | CE0217 (C) | CNE55 (AE) | BG505.T332N (A) | SF162 (B) | MLV (control) |
| BG505 UFO trimer | Addavax | 119 | 62 | 90 | 309 | 59 | 49 | 45 | <40 | 53 | 68 | 58 | <40 | <40 | 120 | 4558 | <40 |
| | | 120 | <40 | 58 | 199 | <40 | <40 | <40 | <40 | <40 | <40 | <40 | <40 | <40 | <100 | 2614 | <40 |
| | | 121 | <40 | 54 | 233 | <40 | <40 | <40 | <40 | <40 | <40 | <40 | <40 | <40 | 446 | 2778 | <40 |
| | | 122 | 45 | 66 | 351 | <40 | <40 | <40 | <40 | <40 | <40 | <40 | <40 | <40 | 889 | 2188 | <40 |
| | | 123 | 282 | 198 | 694 | 128 | 52 | 63 | <40 | 61 | 126 | 103 | <40 | 99 | 351 | 1571 | <40 |
| | | 124 | <40 | 43 | 326 | <40 | <40 | <40 | <40 | <40 | <40 | <40 | <40 | <40 | 1190 | 4701 | <40 |
| BG505 FR SApNP | Addavax | 125 | <40 | 94 | 758 | 59 | <40 | 43 | <40 | 52 | 43 | <40 | <40 | <40 | <100 | 8670 | <40 |
| | | 126 | <40 | 77 | 250 | 43 | <40 | <40 | <40 | 41 | 42 | <40 | <40 | <40 | 1605 | 3579 | <40 |
| | | 127 | <40 | 82 | 426 | <40 | <40 | <40 | <40 | <40 | <40 | <40 | <40 | <40 | 1401 | 13687 | <40 |
| | | 128 | <40 | 96 | 565 | <40 | <40 | <40 | <40 | <40 | <40 | <40 | <40 | <40 | 186 | 10286 | <40 |
| | | 129 | 52 | 107 | 575 | 93 | 59 | 98 | <40 | 52 | 59 | 42 | <40 | <40 | <100 | 6385 | <40 |
| | | 130 | <40 | 79 | 513 | <40 | <40 | <40 | <40 | <40 | <40 | <40 | <40 | <40 | 1774 | 5955 | <40 |
| Env Mix FR SApNP | Addavax | 131 | <40 | 136 | 1093 | 72 | <40 | 73 | <40 | <40 | <40 | <40 | <40 | <40 | <100 | 3596 | <40 |
| | | 132 | <40 | 82 | 450 | 47 | <40 | 54 | <40 | <40 | <40 | <40 | <40 | <40 | 105 | 3580 | <40 |
| | | 133 | 88 | 278 | 680 | 119 | 67 | 137 | 57 | 116 | 100 | 80 | 56 | 100 | <100 | 2971 | <40 |
| | | 134 | <40 | 55 | 249 | <40 | <40 | <40 | <40 | <40 | <40 | <40 | <40 | <40 | 351 | 9968 | <40 |
| | | 135 | <40 | 69 | 296 | <40 | <40 | 51 | <40 | 57 | 57 | <40 | <40 | <40 | <100 | 8622 | <40 |
| | | 136 | <40 | 50 | 418 | <40 | <40 | <40 | <40 | <40 | <40 | <40 | <40 | <40 | <100 | 11188 | <40 |
| BG505 E2p SApNP | Addavax | 155 | 41 | 73 | 171 | 62 | <40 | 66 | <40 | 43 | 55 | 59 | <40 | <40 | 1313 | 3323 | <40 |
| | | 156 | <40 | 40 | 175 | <40 | <40 | 45 | <40 | <40 | <40 | <40 | <40 | <40 | 154 | 4276 | <40 |
| | | 157 | <40 | 105 | 434 | 78 | <40 | 88 | <40 | <40 | <40 | 52 | <40 | <40 | 107 | 6360 | <40 |
| | | 158 | <40 | 70 | 336 | 49 | <40 | 53 | <40 | <40 | <40 | <40 | <40 | <40 | 149 | 2020 | <40 |
| | | 159 | 172 | 86 | 1808 | 271 | <40 | 83 | <40 | 167 | 262 | 97 | 47 | 99 | 198 | 6328 | <40 |
| | | 160 | <40 | <40 | 827 | <40 | <40 | <40 | <40 | <40 | 94 | <40 | <40 | <40 | 1369 | 1422 | <40 |
| Env Mix E2p SApNP | Addavax | 137 | <40 | 77 | 783 | <40 | <40 | 51 | <40 | <40 | <40 | <40 | <40 | <40 | <100 | 2475 | <40 |
| | | 138 | <40 | 121 | 592 | <40 | <40 | <40 | <40 | <40 | <40 | <40 | <40 | <40 | <100 | 7067 | <40 |
| | | 139 | 51 | 161 | 1509 | 51 | 65 | <40 | 40 | <40 | <40 | 41 | <40 | <40 | <100 | 17500 | <40 |
| | | 140 | <40 | 63 | 457 | <40 | <40 | <40 | <40 | <40 | <40 | <40 | <40 | <40 | 144 | 7986 | <40 |
| | | 141 | <40 | 110 | 508 | <40 | <40 | <40 | <40 | <40 | <40 | <40 | <40 | <40 | 576 | 2088 | <40 |
| | | 142 | <40 | 144 | 587 | 44 | <40 | <40 | <40 | <40 | <40 | <40 | <40 | <40 | <100 | 8818 | <40 |
| BG505 I3-01v9 SApNP | Aluminum phosphate | 143 | <40 | 192 | 1717 | <40 | <40 | <40 | <40 | <40 | <40 | <40 | <40 | <40 | 2269 | 4732 | <40 |
| | | 144 | <40 | <40 | 197 | <40 | <40 | 44 | <40 | <40 | <40 | <40 | <40 | <40 | <100 | 12639 | <40 |
| | | 145 | <40 | 94 | 647 | <40 | <40 | 54 | <40 | <40 | <40 | <40 | <40 | <40 | <100 | 12463 | <40 |
| | | 146 | <40 | 95 | 507 | <40 | <40 | 47 | <40 | <40 | 45 | 106 | <40 | <40 | 576 | 5780 | <40 |
| | | 147 | 65 | 202 | 660 | 112 | 130 | 111 | 40 | 80 | 105 | 53 | 51 | 80 | 2235 | 14390 | <40 |
| | | 148 | 61 | 275 | 521 | 79 | 57 | <40 | <40 | 82 | 80 | 62 | <40 | 156 | 108 | 6510 | <40 |
| Env Mix I3-01v9 SApNP | Aluminum phosphate | 149 | <40 | 80 | 1114 | 69 | <40 | <40 | <40 | <40 | <40 | <40 | <40 | <40 | 780 | 3257 | <40 |
| | | 150 | 53 | 154 | 1575 | 116 | 48 | 51 | 44 | 44 | 75 | 81 | <40 | 54 | 631 | 13545 | <40 |
| | | 151 | <40 | 139 | 1675 | 84 | <40 | <40 | 42 | <40 | 58 | <40 | <40 | <40 | 2731 | 16783 | <40 |
| | | 152 | <40 | 68 | 747 | 78 | <40 | <40 | <40 | <40 | 41 | <40 | <40 | <40 | 298 | 13060 | <40 |
| | | 153 | <40 | 288 | 7874 | 45 | <40 | <40 | <40 | 52 | <40 | 417 | <40 | <40 | <100 | 17936 | <40 |
| | | 154 | 51 | 158 | 981 | 69 | 52 | 113 | <40 | 50 | 50 | 74 | <40 | <40 | 145 | 7105 | <40 |

**fig. S5. Neutralization data from the evaluation of Env immunogens with wildtype glycans in mice and rabbits.** (**A**) Mouse IgG neutralization from ten groups of mice (8 per group) that were immunized with BG505 UFO trimer-presenting E2p and I3-01v9 SApNPs. In the first study, five groups of mice were immunized with BG505 UFO trimer-presenting E2p SApNP, each group with a different adjuvant condition: no adjuvant, AddaVax (AV), Aluminum phosphate (AP), Aluminum hydroxide or AVx2/APx2. In a second study, five groups were immunized with BG505 UFO trimer-presenting I3-01v9 SApNP, with the same five adjuvant conditions. Data are shown for neutralization of tier 2 clade A BG505.T332N, tier 1 clade B SF162, and MLV pseudoviruses (MLV-pps) by purified mouse IgG from the last time point, week 11. $IC_{50}$ titers, and neutralization curves are shown. (**B**) Positive samples (>30% autologous neutralization) from (A) were tested against the BG505.T332N variant with the C3/V4 knockout mutation (I396R) and against **(C)** a 12-virus global panel of HIV-1 strains. For (A-C) an IgG concentration of 300 μg/ml was used as the starting point and subjected to a 3-times dilution series in the TZM-bl neutralization assay. Samples were tested in duplicate in the TZM-bl assay, except in the MLV, I396R mutant and the 12-virus global panel assays, where samples were tested in singlet due to a low quantity of remaining IgG. IgG samples no longer available at the time of an assay are marked '-' in the $IC_{50}/IC_{30}$ titer tables. (**D**)-(**G**) Rabbit plasma neutralization from seven groups of rabbits that were immunized with BG505 or mixed Env vaccines. All antigens were mixed with AV except for I3-01v9 SApNPs, which were formulated with AP. BG505 UFO trimer and its three SApNPs (FR, E2p, and I3-01v9) were tested in groups 1, 2, 4, and 6, respectively, with a mixed Env group added for each SApNP platform and tested in groups 3, 5, and 7. The mixed Env vaccine is a cocktail containing equal amounts of SApNPs that present clade A BG505 UFO, clade B H078.14 UFO-BG, and clade C CH505 UFO-BG trimers. Data are shown for (**D**) Neutralization of tier 2 clade A BG505.T332N by rabbit plasma from all groups at every time point between week 0 (w0) and week 28 (w28). The heat-inactivated rabbit plasma was diluted 100 times as the starting point and subjected to a 3-times dilution series in the TZM-bl assay. Samples were tested in duplicate against all viruses. $ID_{50}$ and $ID_{30}$ titers and neutralization curves are graphed for each rabbit at each time point. (**E**) As (D), except neutralization is against tier 1 Clade B SF162. (**F**) As (D) and (E), except neutralization is against MLV-pps as a negative control, and only for key weeks 0, 2, 10 and 26. (**G**) $ID_{50}$ and $ID_{30}$ titers obtained from a large-scale TZM-bl neutralization assay against a 12-virus global panel with rabbit plasma diluted 40 times as the starting point for all 12 isolates. Of note, BG505.T332N, SF162, and MLV data from (D)-(F) were included in this table to facilitate comparison with the 12-virus panel.

**fig. S6**

A

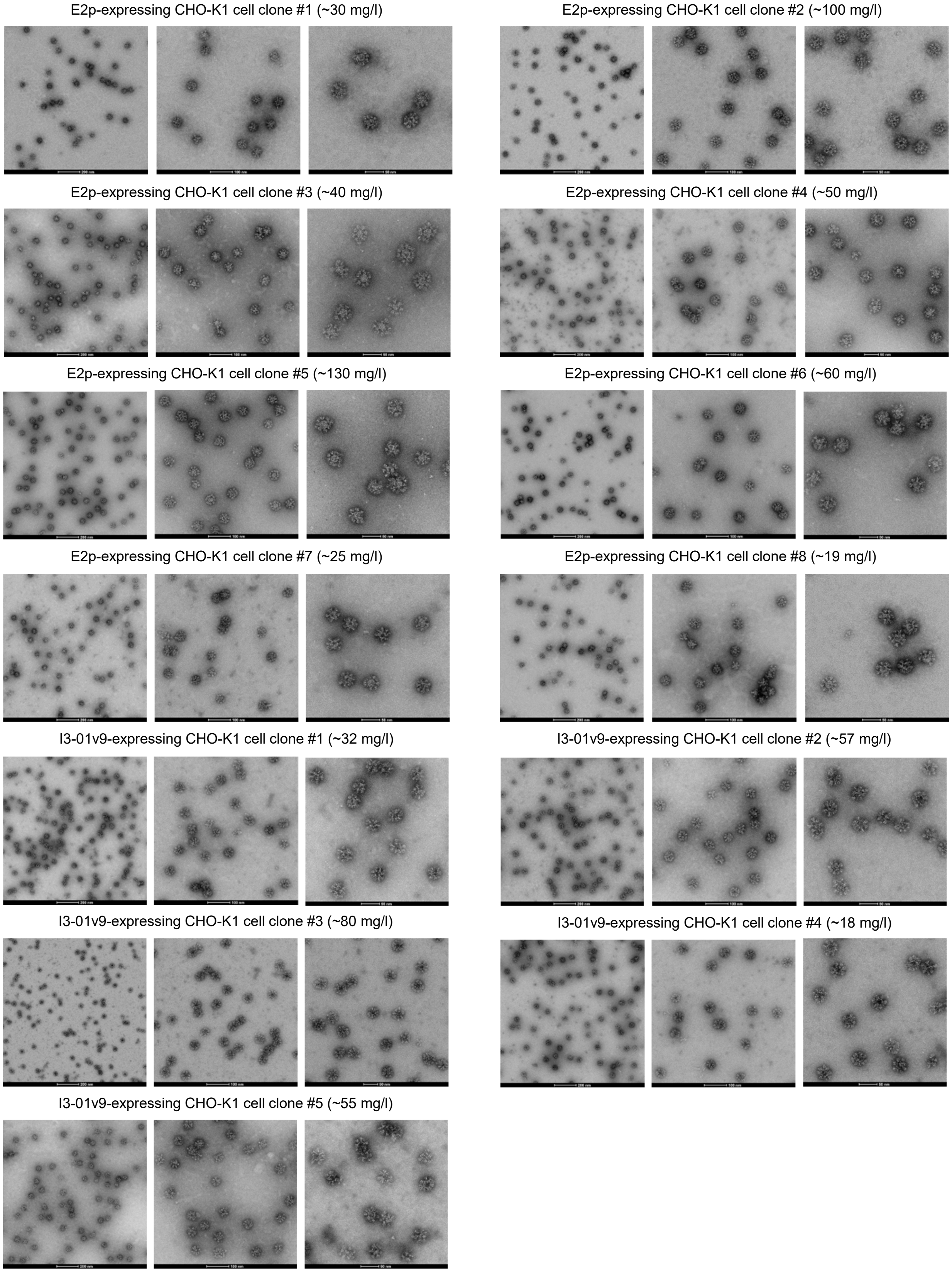
E2p-expressing CHO-K1 cell clone #1 (~30 mg/l)
E2p-expressing CHO-K1 cell clone #2 (~100 mg/l)
E2p-expressing CHO-K1 cell clone #3 (~40 mg/l)
E2p-expressing CHO-K1 cell clone #4 (~50 mg/l)
E2p-expressing CHO-K1 cell clone #5 (~130 mg/l)
E2p-expressing CHO-K1 cell clone #6 (~60 mg/l)
E2p-expressing CHO-K1 cell clone #7 (~25 mg/l)
E2p-expressing CHO-K1 cell clone #8 (~19 mg/l)
I3-01v9-expressing CHO-K1 cell clone #1 (~32 mg/l)
I3-01v9-expressing CHO-K1 cell clone #2 (~57 mg/l)
I3-01v9-expressing CHO-K1 cell clone #3 (~80 mg/l)
I3-01v9-expressing CHO-K1 cell clone #4 (~18 mg/l)
I3-01v9-expressing CHO-K1 cell clone #5 (~55 mg/l)

**fig. S6**

**B** **Week 11 mouse IgG neutralization against HIV-1 BG505.T332N. Immunization with glycan trimmed BG505 UFO trimers.**

| Vaccine | Adjuvant | Glycan trimming regimen | $IC_{50}$ titers (µg/ml) | | | | | | | | |
|---|---|---|---|---|---|---|---|---|---|---|---|
| | | | M1 | M2 | M3 | M4 | M5 | M6 | M7 | M8 | |
| BG505 UFO trimer | AddaVax | Regimen 1 | >300 | >300 | 68 | 20 | >300 | 35 | >300 | 22 | |
| | | Regimen 2 | 1 | 254 | >300 | >300 | 17 | >300 | 93 | 3 | |
| | Aluminum phosphate | Regimen 1 | 17 | 155 | No IgG | 44 | 2 | 8 | 158 | >300 | ★ |
| | | Regimen 2 | 118 | 13 | 5 | 48 | 10 | >300 | >300 | 66 | ★ |

No IgG: indicates no IgG available to test ★ indicates samples from this group further tested against other HIV-1 viruses & MLV

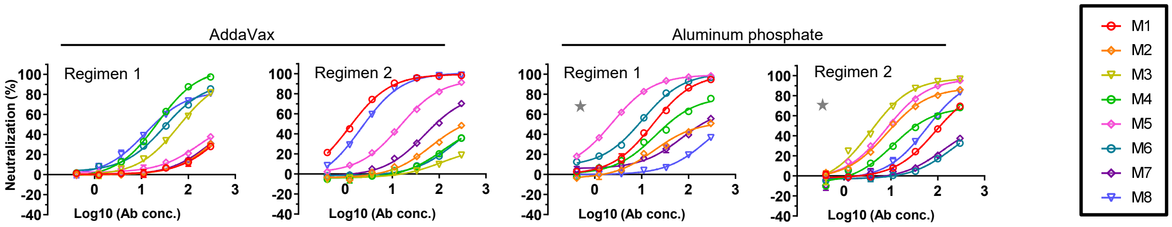


**Week 11 mouse IgG neutralization against HIV SF162**

| Vaccine | Adjuvant | Glycan trimming regimen | $IC_{50}$ titers (µg/ml) | | | | | | | |
|---|---|---|---|---|---|---|---|---|---|---|
| | | | M1 | M2 | M3 | M4 | M5 | M6 | M7 | M8 |
| BG505 UFO trimer | Aluminum phosphate | Regimen 1 | 53 | 15 | No IgG | >300 | 19 | 6 | 5 | 1 |
| | | Regimen 2 | 11 | 34 | 62 | 7 | 15 | 11 | 14 | 1 |

No IgG: indicates no IgG available to test

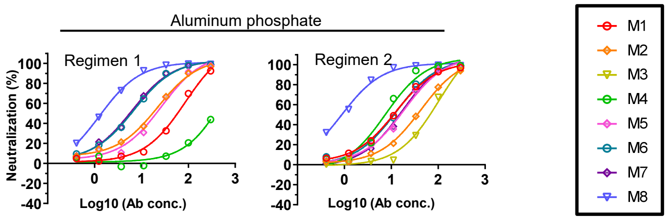


**Week 11 mouse IgG neutralization against MLV (control)**

| Vaccine | Adjuvant | Glycan trimming regimen | $IC_{50}$ titers (µg/ml) | | | | | | | |
|---|---|---|---|---|---|---|---|---|---|---|
| | | | M1 | M2 | M3 | M4 | M5 | M6 | M7 | M8 |
| BG505 UFO trimer | AddaVax | Regimen 1 | >300 | >300 | >300 | >300 | >300 | >300 | >300 | >300 |
| | | Regimen 2 | >300 | >300 | >300 | >300 | >300 | >300 | >300 | >300 |
| | Aluminum phosphate | Regimen 1 | >300 | >300 | No IgG | >300 | >300 | >300 | >300 | >300 |
| | | Regimen 2 | >300 | >300 | >300 | >300 | >300 | >300 | >300 | >300 |

No IgG: indicates no IgG available to test

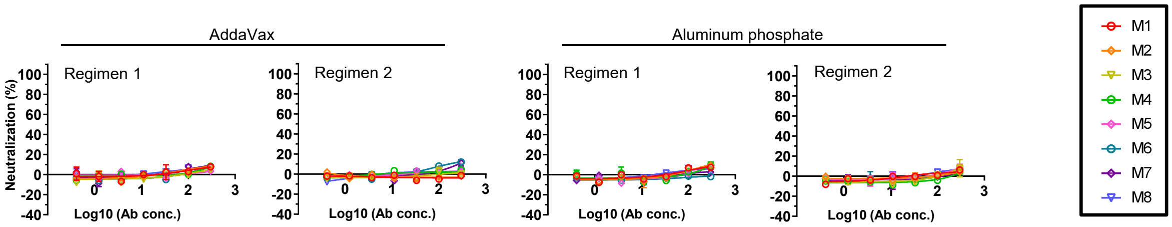

**C** Week 11 mouse IgG neutralization against HIV-1 BG505.T332N. Immunization with glycan trimmed BG505 E2p SApNPs, with repeats.

| Vaccine | Adjuvant | Glycan trimming regimen | Repeat | IC₅₀ titers (µg/ml) | | | | | | | | |
|---|---|---|---|---|---|---|---|---|---|---|---|---|
| | | | | M1 | M2 | M3 | M4 | M5 | M6 | M7 | M8 | |
| BG505 E2p SApNP | AddaVax | Regimen 1 | 1st | >300 | >300 | >300 | >300 | 46 | >300 | >300 | >300 | ★ |
| | | Regimen 2 | 1st | >300 | >300 | >300 | >300 | >300 | >300 | >300 | 70 | ★ |
| | Aluminum phosphate | Regimen 1 | 1st | 293 | 76 | 75 | 108 | 17 | >300 | 16 | >300 | ★ |
| | | Regimen 2 | 1st | 28 | 194 | 11 | >300 | >300 | 27 | >300 | 32 | |
| | | | 2nd | 45 | 65 | 71 | 22 | 20 | 264 | >300 | 220 | |
| | | | 3rd | >300 | 23 | 4 | 199 | >300 | 42 | 8 | >300 | ★ |

★ indicates samples from this group further tested against other HIV-1 viruses & MLV

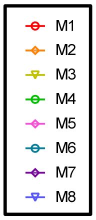


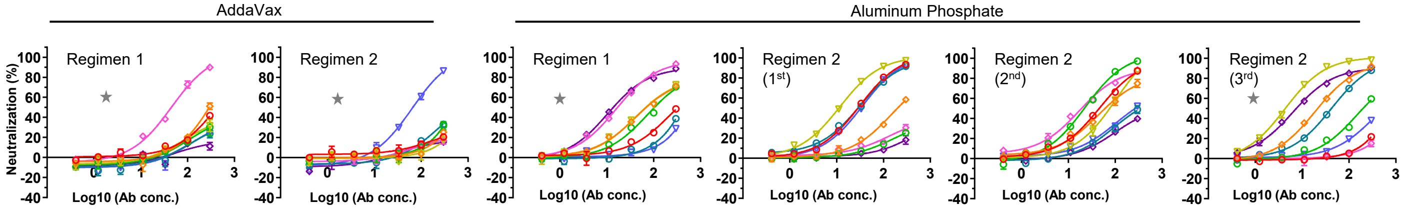


Week 11 mouse IgG neutralization against HIV-1 SF162. Immunization with glycan trimmed BG505 E2p SApNPs.

| Vaccine | Adjuvant | Glycan trimming regimen | IC₅₀ titers (µg/ml) | | | | | | | |
|---|---|---|---|---|---|---|---|---|---|---|
| | | | M1 | M2 | M3 | M4 | M5 | M6 | M7 | M8 |
| BG505 E2p SApNP | AddaVax | Regimen 1 | 5 | 3 | 6 | 6 | 22 | 35 | 121 | 3 |
| | | Regimen 2 | 37 | 9 | 65 | 13 | 46 | 3 | 6 | 3 |
| | Aluminum phosphate | Regimen 1 | 7 | 25 | 14 | 12 | 6 | 24 | 39 | 12 |
| | | Regimen 2 | 8 | 1 | 4 | 29 | 6 | 8 | 18 | 3 |

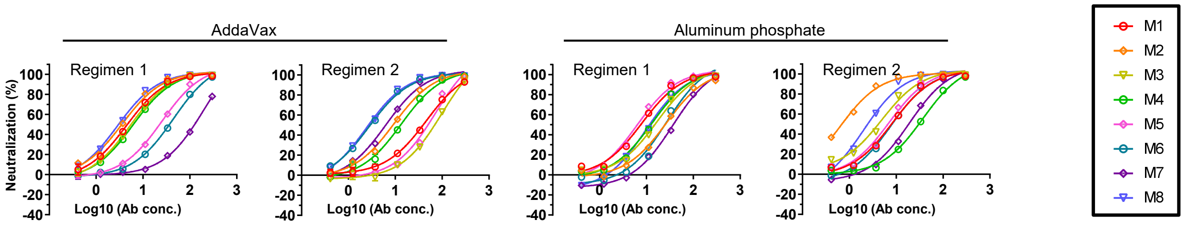


Week 11 mouse IgG neutralization against MLV (control). Immunization with glycan trimmed BG505 E2p SApNPs.

| Vaccine | Adjuvant | Glycan trimming regimen | IC₅₀ titers (µg/ml) | | | | | | | |
|---|---|---|---|---|---|---|---|---|---|---|
| | | | M1 | M2 | M3 | M4 | M5 | M6 | M7 | M8 |
| BG505 E2p SApNP | AddaVax | Regimen 1 | >300 | - | >300 | >300 | >300 | >300 | >300 | >300 |
| | | Regimen 2 | >300 | >300 | >300 | >300 | >300 | >300 | >300 | >300 |
| | Aluminum phosphate | Regimen 1 | >300 | >300 | >300 | >300 | >300 | >300 | >300 | >300 |
| | | Regimen 2 | >300 | >300 | >300 | >300 | >300 | >300 | >300 | >300 |

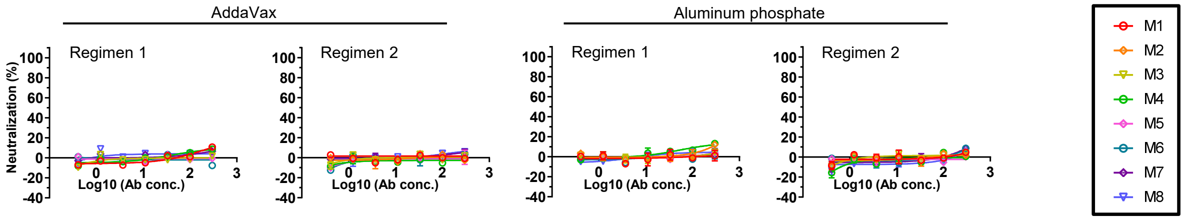

**fig. S6**

**D** **Week 11 mouse IgG neutralization against HIV-1 BG505.T332N. Immunization with glycan trimmed BG505 I3-01v9 SApNPs, with repeats.**

| Vaccine | Adjuvant | Glycan trimming regimen | Repeat | IC$_{50}$ titers (µg/ml) | | | | | | | | |
|---|---|---|---|---|---|---|---|---|---|---|---|---|
| | | | | M1 | M2 | M3 | M4 | M5 | M6 | M7 | M8 | |
| BG505 I3-01v9 SApNP | Aluminum phosphate | Regimen 1 | 1st | 86 | >300 | 275 | >300 | >300 | No IgG | No IgG | 48 | ★ |
| | | | 2nd | >300 | >300 | >300 | >300 | 23 | >300 | >300 | 212 | |
| | | | 3rd | >300 | >300 | >300 | >300 | >300 | 26 | >300 | 203 | |
| | | Regimen 2 | 1st | >300 | 17 | 85 | 37 | 39 | >300 | >300 | >300 | ★ |
| | | | 2nd | 59 | >300 | >300 | >300 | 113 | 148 | 41 | >300 | |
| | | | 3rd | 109 | >300 | >300 | >300 | 40 | 9 | >300 | >300 | |
| | | | 4th | >300 | 57 | 290 | >300 | 16 | >300 | >300 | 14 | |

No IgG: indicates no IgG available to test ★ indicates samples from this group further tested against other HIV-1 viruses & MLV

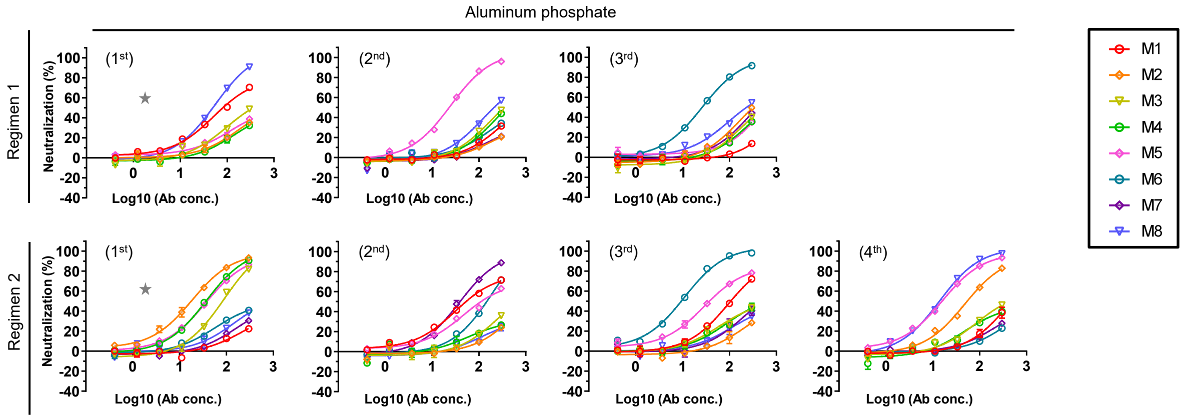


**Week 11 mouse IgG neutralization against HIV-1 SF162. Immunization with glycan trimmed BG505 I3-01v9 SApNPs.**

| Vaccine | Adjuvant | Glycan trimming regimen | IC$_{50}$ titers (µg/ml) | | | | | | | |
|---|---|---|---|---|---|---|---|---|---|---|
| | | | M1 | M2 | M3 | M4 | M5 | M6 | M7 | M8 |
| BG505 I3-01v9 SApNP | Aluminum phosphate | Regimen 1 | 11 | 15 | 5 | 125 | 7 | No IgG | No IgG | 8 |
| | | Regimen 2 | 2 | 16 | 3 | 12 | 8 | 12 | 11 | 11 |

No IgG: indicates no IgG available to test

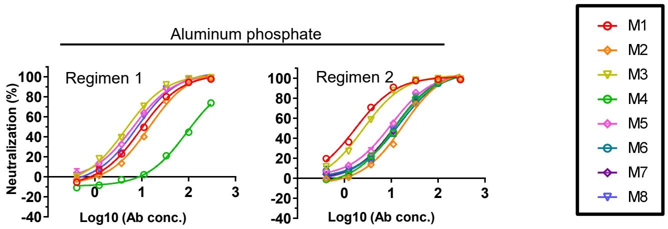


**Week 11 mouse IgG neutralization against MLV (control). Immunization with glycan trimmed BG505 I3-01v9 SApNPs.**

| Vaccine | Adjuvant | Glycan trimming regimen | IC$_{50}$ titers (µg/ml) | | | | | | | |
|---|---|---|---|---|---|---|---|---|---|---|
| | | | M1 | M2 | M3 | M4 | M5 | M6 | M7 | M8 |
| BG505 I3-01v9 SApNP | Aluminum phosphate | Regimen 1 | >300 | >300 | >300 | >300 | >300 | No IgG | No IgG | >300 |
| | | Regimen 2 | >300 | >300 | >300 | >300 | >300 | >300 | >300 | >300 |

No IgG: indicates no IgG available to test

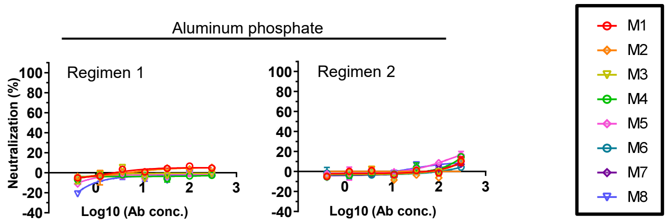

**E** **Week 11 mouse IgG neutralization $IC_{50}$ titers against HIV-1 BG505.T332N and BG505.T332N.I396R mutant**

| Vaccine | Adjuvant | Glycan trimming regimen | Animal ID | $IC_{50}$ titers (µg/ml) BG505.T332N | $IC_{50}$ titers (µg/ml) BG505.T332N.I396R |
|---|---|---|---|---|---|
| BG505 UFO trimer | Aluminum phosphate | Regimen 1 | D-S24 G3-M1 | 17 | >300 |
| | | | D-S24 G3-M2 | 155 | >300 |
| | | | D-S24 G3-M4 | 44 | 172 |
| | | | D-S24 G3-M5 | 2 | >300 |
| | | | D-S24 G3-M6 | 8 | >300 |
| | | | D-S24 G3-M7 | 158 | >300 |
| | | | D-S24 G3-M8 | >300 | >300 |
| | | Regimen 2 | D-S24 G4-M1 | 118 | >300 |
| | | | D-S24 G4-M2 | 13 | >300 |
| | | | D-S24 G4-M3 | 5 | >300 |
| | | | D-S24 G4-M4 | 48 | >300 |
| | | | D-S24 G4-M5 | 10 | >300 |
| | | | D-S24 G4-M6 | >300 | >300 |
| | | | D-S24 G4-M7 | >300 | >300 |
| | | | D-S24 G4-M8 | 66 | >300 |
| BG505 E2p SApNP | AddaVax | Regimen 1 | D-S21 G1-M2 | >300 | 31 |
| | | | D-S21 G1-M3 | >300 | >300 |
| | | | D-S21 G1-M5 | 46 | >300 |
| | | Regimen 2 | D-S21 G2-M4 | >300 | >300 |
| | | | D-S21 G2-M8 | 70 | 11 |

| Vaccine | Adjuvant | Glycan trimming regimen | Animal ID | $IC_{50}$ titers (µg/ml) BG505.T332N | $IC_{50}$ titers (µg/ml) BG505.T332N.I396R |
|---|---|---|---|---|---|
| BG505 E2p SApNP | Aluminum phosphate | Regimen 1 | D-S25 G1-M1 | 293 | >300 |
| | | | D-S25 G1-M2 | 76 | 87 |
| | | | D-S25 G1-M3 | 75 | 22 |
| | | | D-S25 G1-M4 | 108 | 89 |
| | | | D-S25 G1-M5 | 17 | 46 |
| | | | D-S25 G1-M6 | >300 | >300 |
| | | | D-S25 G1-M7 | 16 | >300 |
| | | Regimen 2 | D-S25 G2-M2 | 23 | 31 |
| | | | D-S25 G2-M3 | 4 | 202 |
| | | | D-S25 G2-M4 | 199 | 180 |
| | | | D-S25 G2-M6 | 42 | 27 |
| | | | D-S25 G2-M7 | 8 | >300 |
| | | | D-S25 G2-M8 | >300 | >300 |
| BG505 I3-01v9 SApNP | Aluminum phosphate | Regimen 1 | D-S21 G3-M1 | 86 | 154 |
| | | | D-S21 G3-M3 | 275 | >300 |
| | | | D-S21 G3-M8 | 48 | >300 |
| | | Regimen 2 | D-S21 G4-M2 | 17 | 6 |
| | | | D-S21 G4-M3 | 85 | 23 |
| | | | D-S21 G4-M4 | 37 | 17 |
| | | | D-S21 G4-M5 | 39 | 277 |

**Week 11 mouse IgG neutralization against HIV-1 BG505.T332N and BG505.T332N.I396R mutant**

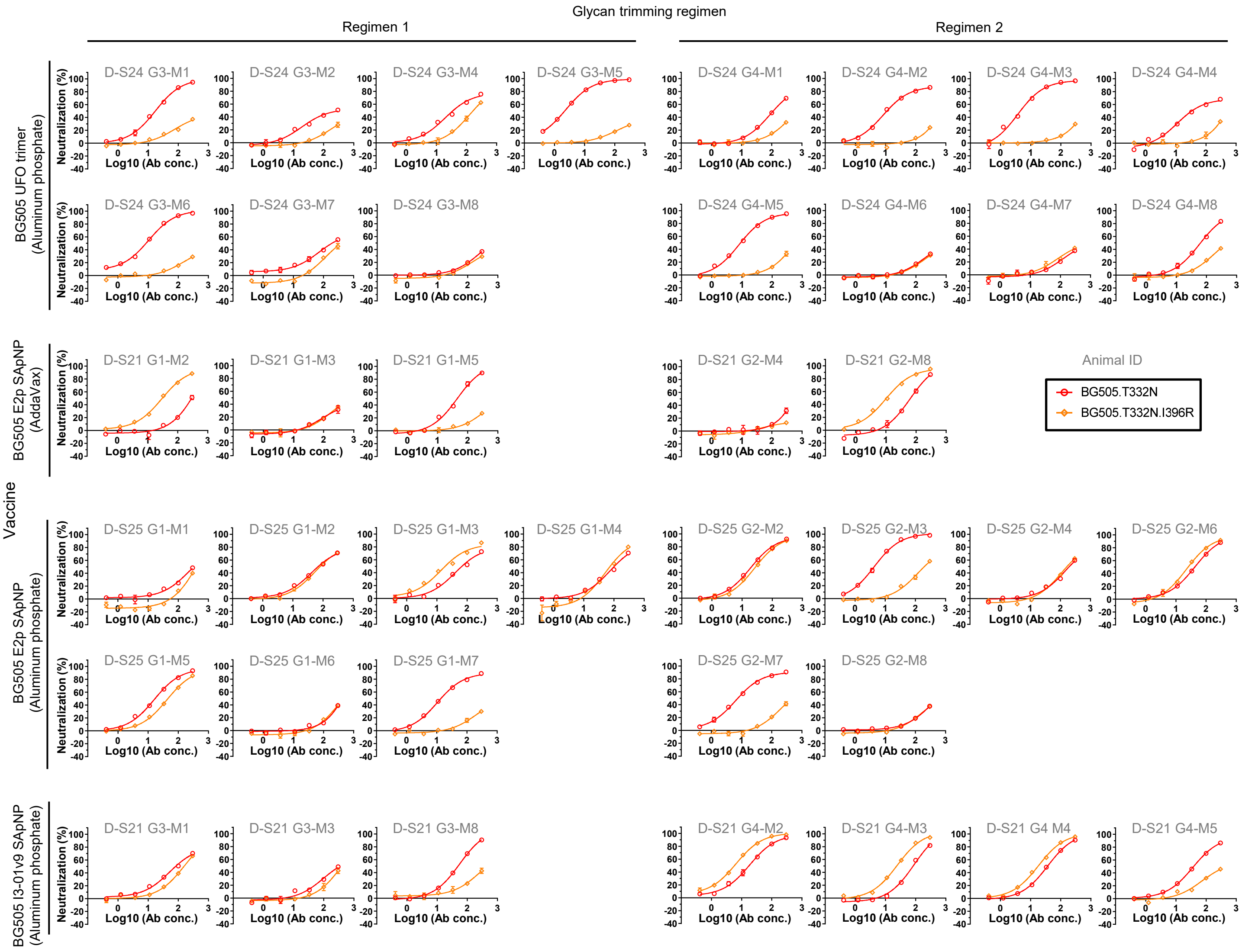

**F** **Week 11 mouse IgG neutralization $IC_{50}$ and $IC_{30}$ titers against HIV-1 12 virus global panel, BG505.T332N, SF162 & MLV (control)**

| Vaccine | Adjuvant | Glycan trimming regimen | Animal ID | $IC_{50}$ titers (μg/ml) for each virus (clade) | | | | | | | | | | | | | | |
|---|---|---|---|---|---|---|---|---|---|---|---|---|---|---|---|---|---|---|
| | | | | TRO11 (B) | 25710 (C) | 398F1 (A) | CNE8 (AE) | X2278 (B) | BJOX-2000 (BC) | X1632 (G) | CE1176 (C) | 246F3 (AC) | CH119 (BC) | CE0217 (C) | CNE55 (AE) | BG505 (A) | SF162 (B) | MLV (control) |
| BG505 UFO Trimer | Aluminum phosphate | Regimen 1 | D-S24 G3-M1 | >300 | >300 | >300 | >300 | >300 | >300 | >300 | >300 | >300 | - | - | - | 17 | 53 | >300 |
| | | | D-S24 G3-M2 | >300 | >300 | >300 | >300 | >300 | >300 | >300 | >300 | >300 | >300 | >300 | >300 | 155 | 15 | >300 |
| | | | D-S24 G3-M4 | >300 | >300 | >300 | >300 | >300 | >300 | >300 | >300 | >300 | >300 | >300 | >300 | 44 | >300 | >300 |
| | | | D-S24 G3-M5 | >300 | >300 | >300 | >300 | >300 | >300 | >300 | >300 | >300 | - | - | - | 2 | 19 | >300 |
| | | | D-S24 G3-M6 | >300 | >300 | >300 | >300 | >300 | >300 | 236 | >300 | >300 | >300 | >300 | >300 | 8 | 6 | >300 |
| | | | D-S24 G3-M7 | >300 | >300 | >300 | >300 | >300 | >300 | >300 | >300 | >300 | >300 | >300 | >300 | 158 | 5 | >300 |
| | | Regimen 2 | D-S24 G4-M1 | - | >300 | >300 | >300 | >300 | >300 | >300 | >300 | >300 | - | - | - | 118 | 11 | >300 |
| | | | D-S24 G4-M2 | - | >300 | >300 | >300 | >300 | >300 | - | >300 | >300 | - | - | - | 13 | 34 | >300 |
| | | | D-S24 G4-M3 | >300 | >300 | 256 | >300 | >300 | >300 | >300 | >300 | >300 | - | - | - | 5 | 62 | >300 |
| | | | D-S24 G4-M4 | >300 | >300 | 256 | >300 | >300 | >300 | >300 | >300 | >300 | >300 | >300 | >300 | 48 | 7 | >300 |
| | | | D-S24 G4-M5 | >300 | >300 | >300 | >300 | >300 | >300 | >300 | >300 | >300 | >300 | >300 | >300 | 10 | 15 | >300 |
| | | | D-S24 G4-M8 | >300 | >300 | 138 | >300 | >300 | >300 | 177 | >300 | >300 | >300 | >300 | >300 | 66 | 1 | >300 |
| BG505 E2p SApNP | AddaVax | Regimen 1 | D-S21 G1-M3 | >300 | >300 | 275 | 151 | 263 | >300 | >300 | 217 | >300 | >300 | >300 | >300 | >300 | 6 | - |
| | | | D-S21 G1-M5 | >300 | 300 | 225 | 195 | - | >300 | >300 | >300 | >300 | >300 | >300 | - | 46 | 22 | >300 |
| | | Regimen 2 | D-S21 G2-M8 | >300 | >300 | >300 | >300 | - | >300 | >300 | >300 | >300 | >300 | >300 | - | 70 | 3 | >300 |
| BG505 E2p SApNP | Aluminum phosphate | Regimen 1 | D-S25 G1-M1 | >300 | >300 | 264 | >300 | >300 | >300 | 247 | >300 | >300 | >300 | >300 | >300 | 293 | 7 | >300 |
| | | | D-S25 G1-M2 | - | - | 238 | >300 | - | - | - | >300 | - | - | - | - | 76 | 25 | >300 |
| | | | D-S25 G1-M3 | >300 | >300 | 285 | >300 | >300 | >300 | >300 | >300 | >300 | >300 | >300 | >300 | 75 | 14 | >300 |
| | | | D-S25 G1-M4 | - | 199 | 221 | >300 | - | >300 | - | >300 | - | - | - | - | 108 | 12 | >300 |
| | | | D-S25 G1-M5 | - | >300 | 251 | >300 | >300 | >300 | - | >300 | >300 | - | - | - | 17 | 6 | >300 |
| | | | D-S25 G1-M7 | >300 | >300 | >300 | >300 | >300 | >300 | >300 | >300 | >300 | >300 | >300 | >300 | 16 | 39 | >300 |
| | | Regimen 2 | D-S25 G2-M2 | >300 | >300 | 200 | >300 | >300 | >300 | 216 | >300 | >300 | - | - | - | 23 | 1 | >300 |
| | | | D-S25 G2-M3 | >300 | >300 | 260 | >300 | >300 | >300 | >300 | >300 | >300 | >300 | - | - | 4 | 4 | >300 |
| | | | D-S25 G2-M4 | >300 | 263 | 153 | >300 | >300 | >300 | >300 | >300 | >300 | >300 | >300 | >300 | 199 | 29 | >300 |
| | | | D-S25 G2-M6 | >300 | >300 | 239 | >300 | >300 | >300 | >300 | >300 | >300 | >300 | >300 | >300 | 42 | 8 | >300 |
| | | | D-S25 G2-M7 | >300 | >300 | 285 | >300 | >300 | >300 | >300 | >300 | >300 | >300 | >300 | >300 | 8 | 18 | >300 |
| BG505 I3-01v9 SApNP | Aluminum phosphate | Regimen 1 | D-S21 G3-M1 | >300 | >300 | 298 | >300 | >300 | >300 | >300 | >300 | >300 | >300 | >300 | >300 | 86 | 11 | >300 |
| | | | D-S21 G3-M3 | >300 | >300 | >300 | >300 | >300 | >300 | >300 | >300 | >300 | >300 | >300 | >300 | 275 | 5 | >300 |
| | | | D-S21 G3-M8 | >300 | >300 | >300 | 224 | >300 | >300 | >300 | >300 | >300 | >300 | >300 | >300 | 48 | 8 | >300 |
| | | Regimen 2 | D-S21 G4-M2 | >300 | >300 | 291 | >300 | >300 | >300 | >300 | >300 | >300 | >300 | >300 | >300 | 17 | 16 | >300 |
| | | | D-S21 G4-M3 | >300 | >300 | 236 | 236 | >300 | >300 | >300 | >300 | >300 | >300 | >300 | >300 | 85 | 3 | >300 |
| | | | D-S21 G4-M4 | >300 | >300 | >300 | >300 | >300 | >300 | >300 | >300 | >300 | >300 | >300 | - | 37 | 12 | >300 |
| | | | D-S21 G4-M5 | >300 | 267 | 284 | >300 | >300 | >300 | >300 | >300 | >300 | >300 | >300 | >300 | 39 | 8 | >300 |

| Vaccine | Adjuvant | Glycan trimming regimen | Animal ID | $IC_{30}$ titers (μg/ml) for each virus (clade) | | | | | | | | | | | | | | |
|---|---|---|---|---|---|---|---|---|---|---|---|---|---|---|---|---|---|---|
| | | | | TRO11 (B) | 25710 (C) | 398F1 (A) | CNE8 (AE) | X2278 (B) | BJOX-2000 (BC) | X1632 (G) | CE1176 (C) | 246F3 (AC) | CH119 (BC) | CE0217 (C) | CNE55 (AE) | BG505 (A) | SF162 (B) | MLV (control) |
| BG505 UFO Trimer | Aluminum phosphate | Regimen 1 | D-S24 G3-M1 | 235 | 167 | 178 | 265 | >300 | 158 | >300 | >300 | >300 | - | - | - | 7 | 23 | >300 |
| | | | D-S24 G3-M2 | >300 | 284 | 295 | >300 | >300 | 269 | 282 | >300 | >300 | >300 | >300 | >300 | 66 | 7 | >300 |
| | | | D-S24 G3-M4 | >300 | 246 | >300 | >300 | >300 | 257 | >300 | >300 | >300 | >300 | >300 | >300 | 19 | 168 | >300 |
| | | | D-S24 G3-M5 | 273 | 196 | 285 | 258 | 180 | 192 | 233 | >300 | 283 | - | - | - | 1 | 8 | >300 |
| | | | D-S24 G3-M6 | >300 | 191 | 275 | 229 | 285 | 213 | 101 | >300 | >300 | 298 | >300 | >300 | 3 | 2 | >300 |
| | | | D-S24 G3-M7 | >300 | >300 | >300 | >300 | 279 | 283 | 276 | >300 | >300 | >300 | 289 | >300 | 68 | 2 | >300 |
| | | Regimen 2 | D-S24 G4-M1 | - | 246 | 275 | 245 | 160 | 204 | 186 | >300 | 283 | - | - | - | 51 | 5 | >300 |
| | | | D-S24 G4-M2 | - | >300 | 226 | 288 | 183 | 270 | - | >300 | >300 | - | - | - | 6 | 14 | >300 |
| | | | D-S24 G4-M3 | 261 | >300 | 110 | 280 | 284 | >300 | >300 | >300 | >300 | - | - | - | 2 | 26 | >300 |
| | | | D-S24 G4-M4 | >300 | >300 | 110 | >300 | >300 | >300 | 296 | >300 | >300 | >300 | >300 | >300 | 21 | 3 | >300 |
| | | | D-S24 G4-M5 | >300 | >300 | 133 | 249 | 244 | 177 | 161 | >300 | >300 | >300 | >300 | >300 | 4 | 6 | >300 |
| | | | D-S24 G4-M8 | >300 | 254 | 59 | 220 | >300 | 132 | 76 | 180 | 173 | 171 | >300 | >300 | 28 | 0 | >300 |
| BG505 E2p SApNP | AddaVax | Regimen 1 | D-S21 G1-M3 | 225 | 149 | 118 | 65 | 113 | >300 | 281 | 93 | 206 | 275 | >300 | >300 | 249 | 2 | - |
| | | | D-S21 G1-M5 | >300 | 129 | 97 | 84 | - | 166 | 208 | 168 | 277 | 277 | >300 | - | 20 | 9 | >300 |
| | | Regimen 2 | D-S21 G2-M8 | >300 | >300 | 209 | 182 | - | >300 | 225 | >300 | >300 | >300 | >300 | - | 30 | 1 | >300 |
| BG505 E2p SApNP | Aluminum phosphate | Regimen 1 | D-S25 G1-M1 | >300 | 194 | 113 | 226 | >300 | 212 | 106 | >300 | >300 | >300 | >300 | >300 | 126 | 3 | >300 |
| | | | D-S25 G1-M2 | - | - | 102 | 207 | - | - | - | >300 | - | - | - | - | 33 | 11 | >300 |
| | | | D-S25 G1-M3 | >300 | >300 | 122 | >300 | >300 | >300 | >300 | >300 | >300 | >300 | >300 | >300 | 32 | 6 | >300 |
| | | | D-S25 G1-M4 | - | 85 | 95 | 269 | - | >300 | - | >300 | - | - | - | - | 46 | 5 | >300 |
| | | | D-S25 G1-M5 | - | >300 | 107 | 180 | >300 | 239 | - | >300 | >300 | - | - | - | 7 | 3 | >300 |
| | | | D-S25 G1-M7 | 249 | >300 | 137 | 256 | >300 | 285 | >300 | >300 | >300 | >300 | >300 | >300 | 7 | 17 | >300 |
| | | Regimen 2 | D-S25 G2-M2 | >300 | >300 | 86 | >300 | >300 | 140 | 93 | >300 | >300 | - | - | - | 10 | 0 | >300 |
| | | | D-S25 G2-M3 | >300 | >300 | 111 | 254 | >300 | 198 | >300 | >300 | >300 | >300 | - | - | 2 | 2 | >300 |
| | | | D-S25 G2-M4 | >300 | 113 | 66 | 129 | 224 | 143 | >300 | >300 | >300 | >300 | >300 | >300 | 85 | 12 | >300 |
| | | | D-S25 G2-M6 | >300 | >300 | 103 | >300 | >300 | >300 | 230 | >300 | >300 | >300 | >300 | >300 | 18 | 4 | >300 |
| | | | D-S25 G2-M7 | 236 | >300 | 122 | 254 | >300 | >300 | 187 | >300 | >300 | >300 | >300 | >300 | 4 | 8 | >300 |
| BG505 I3-01v9 SApNP | Aluminum phosphate | Regimen 1 | D-S21 G3-M1 | >300 | 192 | 128 | >300 | 162 | >300 | >300 | 144 | 268 | >300 | >300 | >300 | 37 | 5 | >300 |
| | | | D-S21 G3-M3 | >300 | 153 | 190 | 129 | >300 | >300 | 204 | 178 | 256 | >300 | >300 | >300 | 118 | 2 | >300 |
| | | | D-S21 G3-M8 | 266 | 137 | 160 | 96 | >300 | 286 | 253 | 187 | 254 | >300 | >300 | >300 | 21 | 3 | >300 |
| | | Regimen 2 | D-S21 G4-M2 | 292 | 176 | 125 | >300 | 274 | >300 | >300 | 208 | >300 | >300 | >300 | >300 | 7 | 7 | >300 |
| | | | D-S21 G4-M3 | 256 | 130 | 101 | 101 | 284 | 175 | 145 | 164 | 188 | 208 | >300 | >300 | 37 | 1 | >300 |
| | | | D-S21 G4-M4 | >300 | 143 | 151 | 153 | >300 | 213 | >300 | 215 | 281 | >300 | >300 | - | 16 | 5 | >300 |
| | | | D-S21 G4-M5 | >300 | 267 | 284 | >300 | >300 | >300 | >300 | >300 | >300 | >300 | >300 | >300 | 39 | 4 | >300 |

**Week 11 mouse IgG neutralization against HIV-1 12 virus global panel**

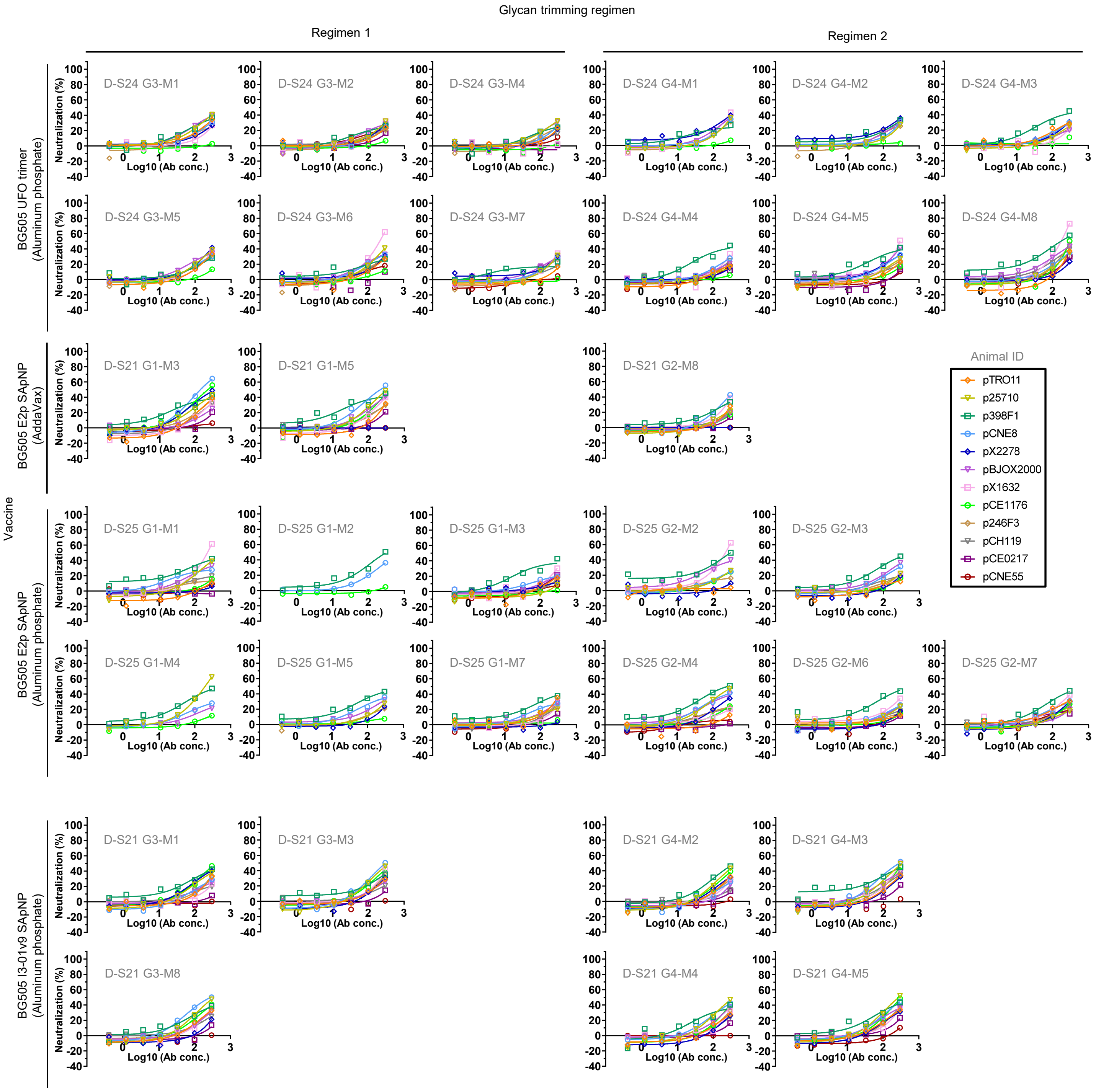

G **Week 0 rabbit IgG neutralization against HIV-1 BG505.T332N**

| Vaccine | Adjuvant | Glycan trimming regimen | IC$_{50}$ titers (µg/ml) | | | | | |
|---|---|---|---|---|---|---|---|---|
| | | | R1 | R2 | R3 | R4 | R5 | R6 |
| BG505 E2p SApNP | AddaVax | Regimen 1 | >300 | >300 | >300 | >300 | >300 | >300 |
| | | Regimen 2 | >300 | >300 | >300 | >300 | >300 | >300 |
| BG505 I3-01v9 SApNP | Aluminum phosphate | Regimen 1 | >300 | >300 | >300 | >300 | >300 | >300 |
| | | Regimen 2 | >300 | >300 | >300 | >300 | >300 | >300 |

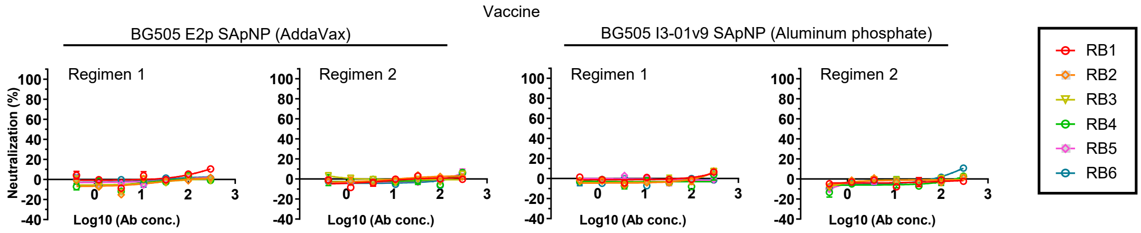


**Week 0 rabbit IgG neutralization against HIV-1 SF162**

| Vaccine | Adjuvant | Glycan trimming regimen | IC$_{50}$ titers (µg/ml) | | | | | |
|---|---|---|---|---|---|---|---|---|
| | | | R1 | R2 | R3 | R4 | R5 | R6 |
| BG505 E2p SApNP | AddaVax | Regimen 1 | >300 | >300 | >300 | >300 | >300 | >300 |
| | | Regimen 2 | >300 | >300 | >300 | >300 | >300 | >300 |
| BG505 I3-01v9 SApNP | Aluminum phosphate | Regimen 1 | >300 | >300 | >300 | >300 | >300 | >300 |
| | | Regimen 2 | >300 | >300 | >300 | >300 | >300 | >300 |

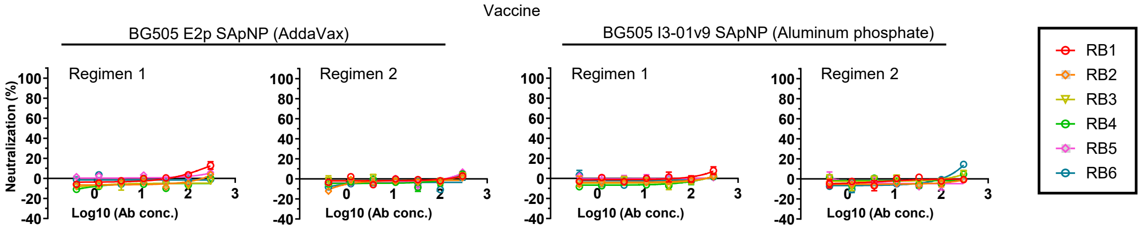


**Week 0 rabbit IgG neutralization against MLV (control)**

| Vaccine | Adjuvant | Glycan trimming regimen | IC$_{50}$ titers (µg/ml) | | | | | |
|---|---|---|---|---|---|---|---|---|
| | | | R1 | R2 | R3 | R4 | R5 | R6 |
| BG505 E2p SApNP | AddaVax | Regimen 1 | >300 | >300 | >300 | >300 | >300 | >300 |
| | | Regimen 2 | >300 | >300 | >300 | >300 | >300 | >300 |
| BG505 I3-01v9 SApNP | Aluminum phosphate | Regimen 1 | >300 | >300 | >300 | >300 | >300 | >300 |
| | | Regimen 2 | >300 | >300 | >300 | >300 | >300 | >300 |

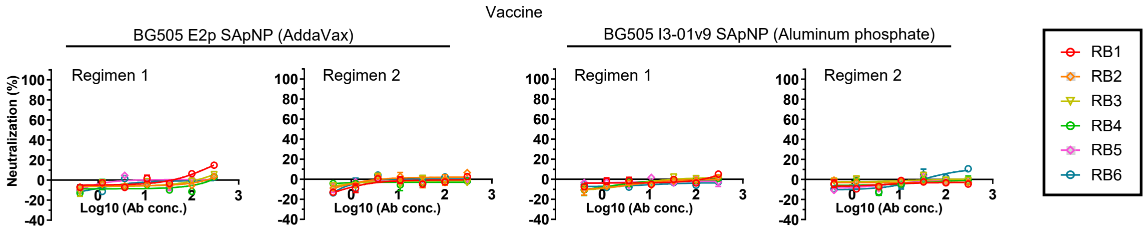

fig. S6

H **Week 11 rabbit IgG neutralization against HIV-1 BG505.T332N**

| Vaccine | Adjuvant | Glycan trimming regimen | $IC_{50}$ titers (μg/ml) | | | | | |
|---|---|---|---|---|---|---|---|---|
| | | | R1 | R2 | R3 | R4 | R5 | R6 |
| BG505 E2p SApNP | AddaVax | Regimen 1 | >300 | 39 | >300 | >300 | >300 | 272 |
| | | Regimen 2 | 233 | >300 | >300 | >300 | >300 | >300 |
| BG505 I3-01v9 SApNP | Aluminum phosphate | Regimen 1 | >300 | 36 | 193 | 79 | >300 | >300 |
| | | Regimen 2 | >300 | 93 | >300 | 44 | 167 | 23 |

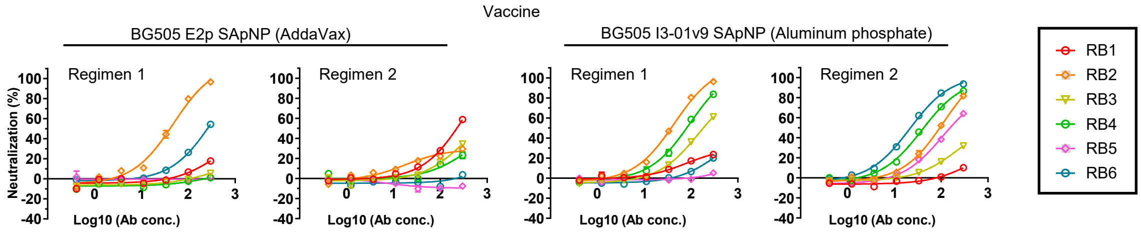


**Week 11 rabbit IgG neutralization against HIV-1 SF162**

| Vaccine | Adjuvant | Glycan trimming regimen | $IC_{50}$ titers (μg/ml) | | | | | |
|---|---|---|---|---|---|---|---|---|
| | | | R1 | R2 | R3 | R4 | R5 | R6 |
| BG505 E2p SApNP | AddaVax | Regimen 1 | 10 | 18 | 11 | 22 | 48 | 4 |
| | | Regimen 2 | 18 | 5 | 3 | 15 | 8 | 26 |
| BG505 I3-01v9 SApNP | Aluminum phosphate | Regimen 1 | 3 | 4 | 4 | 22 | 41 | 9 |
| | | Regimen 2 | 2 | 6 | 6 | 4 | 2 | 11 |

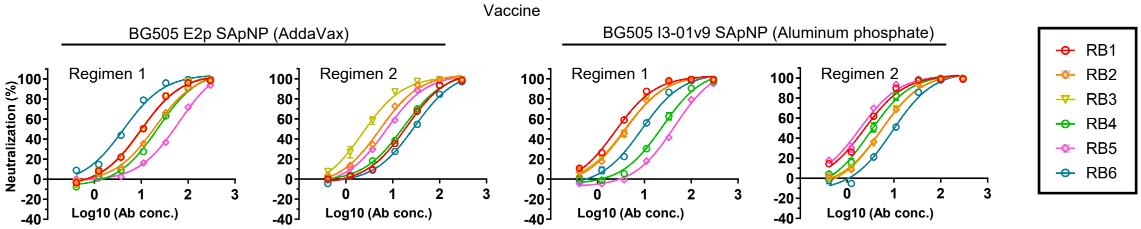


**Week 11 rabbit IgG neutralization against MLV (control)**

| Vaccine | Adjuvant | Glycan trimming regimen | $IC_{50}$ titers (μg/ml) | | | | | |
|---|---|---|---|---|---|---|---|---|
| | | | R1 | R2 | R3 | R4 | R5 | R6 |
| BG505 E2p SApNP | AddaVax | Regimen 1 | >300 | >300 | >300 | >300 | >300 | >300 |
| | | Regimen 2 | >300 | >300 | >300 | >300 | >300 | >300 |
| BG505 I3-01v9 SApNP | Aluminum phosphate | Regimen 1 | >300 | >300 | >300 | >300 | >300 | >300 |
| | | Regimen 2 | >300 | >300 | >300 | >300 | 291 | >300 |

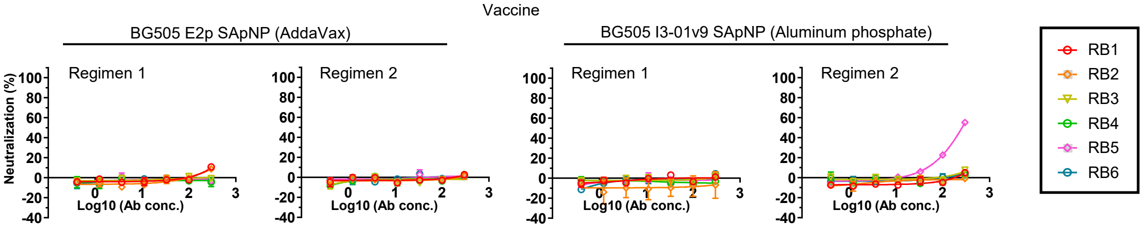

**fig. S6**

**Week 11 rabbit IgG neutralization $IC_{50}$ titers against BG505.T332N and glycan hole mutant viruses**

| Vaccine | Adjuvant | Glycan trimming regimen | Animal ID | IC50 values (µg/ml) BG505. T332N | BG505. T332N. Q130N | BG505. T332N. S241N | BG505. T332N. P291T | BG505. T332N. T465N |
|---|---|---|---|---|---|---|---|---|
| BG505 E2p SApNP | AddaVax | Regimen 1 | G1-2 | 39 | 31 | 15 | 29 | >300 |
| | | | G1-6 | 272 | >300 | >300 | >300 | 231 |
| | | Regimen 2 | G2-1 | 233 | 174 | 85 | 138 | >300 |
| | | | G2-3 | >300 | >300 | >300 | >300 | >300 |
| BG505 I3-01v9 SApNP | Aluminum phosphate | Regimen 1 | G3-2 | 36 | 26 | 17 | 20 | >300 |
| | | | G3-3 | 193 | 227 | >300 | >300 | 186 |
| | | | G3-4 | 79 | 60 | 31 | 52 | >300 |
| | | Regimen 2 | G4-2 | 93 | 73 | 25 | 59 | 169 |
| | | | G4-3 | >300 | >300 | 74 | 214 | 211 |
| | | | G4-4 | 44 | 29 | 12 | 33 | >300 |
| | | | G4-5 | 167 | 112 | 119 | 157 | >300 |
| | | | G4-6 | 23 | 14 | 8 | 12 | 18 |

**Week 11 rabbit IgG neutralization against BG505.T332N and glycan hole mutant viruses**

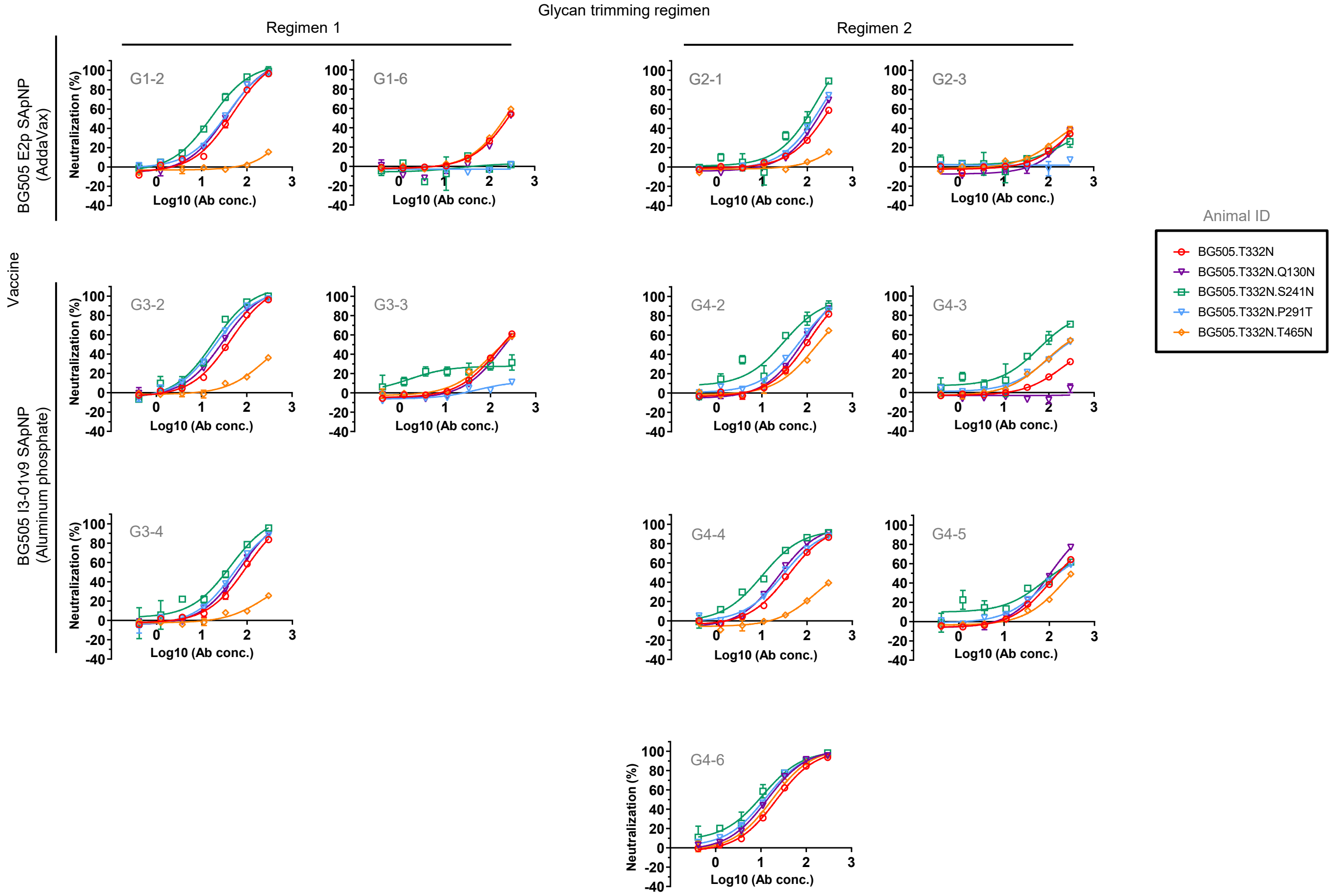

### fig. S6

J Week 11 rabbit IgG neutralization $IC_{50}$ and $IC_{30}$ titers against HIV-1 12 virus panel, BG505.T332N, SF162 & MLV pSG3

| Vaccine | Adjuvant | Glycan trimming regimen | Animal ID | $IC_{50}$ titers (μg/ml) for each virus (clade) | | | | | | | | | | | | | | |
|---|---|---|---|---|---|---|---|---|---|---|---|---|---|---|---|---|---|---|
| | | | | TRO11 (B) | 25710 (C) | 398F1 (A) | CNE8 (AE) | X2278 (B) | BJOX-2000 (BC) | X1632 (G) | CE1176 (C) | 246F3 (AC) | CH119 (BC) | CE0217 (C) | CNE55 (AE) | BG505. T332N (A) | SF162 (B) | MLV (control) |
| BG505 E2p SApNP | AddaVax | Regimen 1 | G1-2 | >300 | >300 | >300 | >300 | >300 | >300 | >300 | >300 | >300 | >300 | >300 | >300 | 39 | 18 | >300 |
| | | | G1-6 | >300 | >300 | >300 | >300 | >300 | >300 | >300 | >300 | >300 | >300 | >300 | >300 | 272 | 4 | >300 |
| | | Regimen 2 | G2-1 | >300 | >300 | >300 | >300 | >300 | >300 | >300 | >300 | >300 | >300 | >300 | >300 | 233 | 18 | >300 |
| | | | G2-3 | >300 | 279 | >300 | 240 | >300 | >300 | >300 | >300 | >300 | >300 | >300 | >300 | >300 | 3 | >300 |
| BG505 I3-01v9 SApNP | Aluminum phosphate | Regimen 1 | G3-2 | >300 | >300 | >300 | >300 | >300 | >300 | >300 | >300 | >300 | >300 | >300 | >300 | 36 | 4 | >300 |
| | | | G3-3 | >300 | >300 | >300 | 277 | >300 | >300 | >300 | >300 | >300 | >300 | >300 | >300 | 193 | 4 | >300 |
| | | | G3-4 | >300 | >300 | >300 | >300 | >300 | >300 | >300 | >300 | >300 | >300 | >300 | >300 | 79 | 22 | >300 |
| | | Regimen 2 | G4-2 | >300 | >300 | >300 | >300 | >300 | >300 | >300 | >300 | >300 | >300 | >300 | >300 | 93 | 6 | >300 |
| | | | G4-3 | >300 | >300 | >300 | >300 | >300 | >300 | >300 | >300 | >300 | >300 | >300 | >300 | >300 | 6 | >300 |
| | | | G4-4 | >300 | >300 | >300 | >300 | >300 | >300 | >300 | >300 | >300 | >300 | >300 | >300 | 44 | 4 | >300 |
| | | | G4-5 | >300 | 201 | 198 | 134 | >300 | 156 | 210 | 177 | 239 | 218 | >300 | >300 | 167 | 2 | 291 |
| | | | G4-6 | >300 | >300 | >300 | >300 | >300 | >300 | >300 | >300 | >300 | >300 | >300 | >300 | 23 | 11 | >300 |

| Vaccine | Adjuvant | Glycan trimming regimen | Animal ID | $IC_{30}$ titers (μg/ml) for each virus (clade) | | | | | | | | | | | | | | |
|---|---|---|---|---|---|---|---|---|---|---|---|---|---|---|---|---|---|---|
| | | | | TRO11 (B) | 25710 (C) | 398F1 (A) | CNE8 (AE) | X2278 (B) | BJOX-2000 (BC) | X1632 (G) | CE1176 (C) | 246F3 (AC) | CH119 (BC) | CE0217 (C) | CNE55 (AE) | BG505. T332N (A) | SF162 (B) | MLV (control) |
| BG505 E2p SApNP | AddaVax | Regimen 1 | G1-2 | >300 | >300 | >300 | >300 | >300 | >300 | >300 | >300 | >300 | >300 | >300 | >300 | 17 | 8 | >300 |
| | | | G1-6 | >300 | 272 | >300 | >300 | >300 | >300 | >300 | >300 | >300 | >300 | >300 | >300 | 116 | 2 | >300 |
| | | Regimen 2 | G2-1 | >300 | >300 | >300 | >300 | >300 | >300 | >300 | >300 | >300 | >300 | >300 | >300 | 100 | 8 | >300 |
| | | | G2-3 | >300 | 120 | 278 | 103 | >300 | 260 | >300 | >300 | >300 | >300 | >300 | >300 | 243 | 1 | >300 |
| BG505 I3-01v9 SApNP | Aluminum phosphate | Regimen 1 | G3-2 | >300 | 263 | >300 | 251 | >300 | >300 | >300 | >300 | >300 | >300 | >300 | >300 | 16 | 2 | >300 |
| | | | G3-3 | >300 | 132 | 253 | 119 | >300 | >300 | >300 | >300 | >300 | >300 | >300 | >300 | 83 | 2 | >300 |
| | | | G3-4 | >300 | 183 | 233 | 196 | >300 | >300 | >300 | >300 | >300 | >300 | >300 | >300 | 34 | 9 | >300 |
| | | Regimen 2 | G4-2 | >300 | 220 | >300 | 203 | >300 | >300 | >300 | >300 | >300 | >300 | >300 | >300 | 40 | 3 | >300 |
| | | | G4-3 | >300 | 162 | 195 | 166 | >300 | >300 | >300 | >300 | >300 | >300 | >300 | >300 | 262 | 3 | >300 |
| | | | G4-4 | >300 | 166 | 144 | 134 | >300 | 257 | >300 | >300 | >300 | >300 | >300 | >300 | 19 | 2 | >300 |
| | | | G4-5 | 148 | 86 | 85 | 58 | 192 | 67 | 90 | 76 | 103 | 94 | 158 | 182 | 71 | 1 | 125 |
| | | | G4-6 | >300 | 160 | 198 | 174 | >300 | >300 | >300 | >300 | >300 | >300 | >300 | >300 | 10 | 5 | >300 |

Week 11 rabbit IgG neutralization against HIV-1 12 virus panel, BG505.T332N, SF162 & MLV pSG3

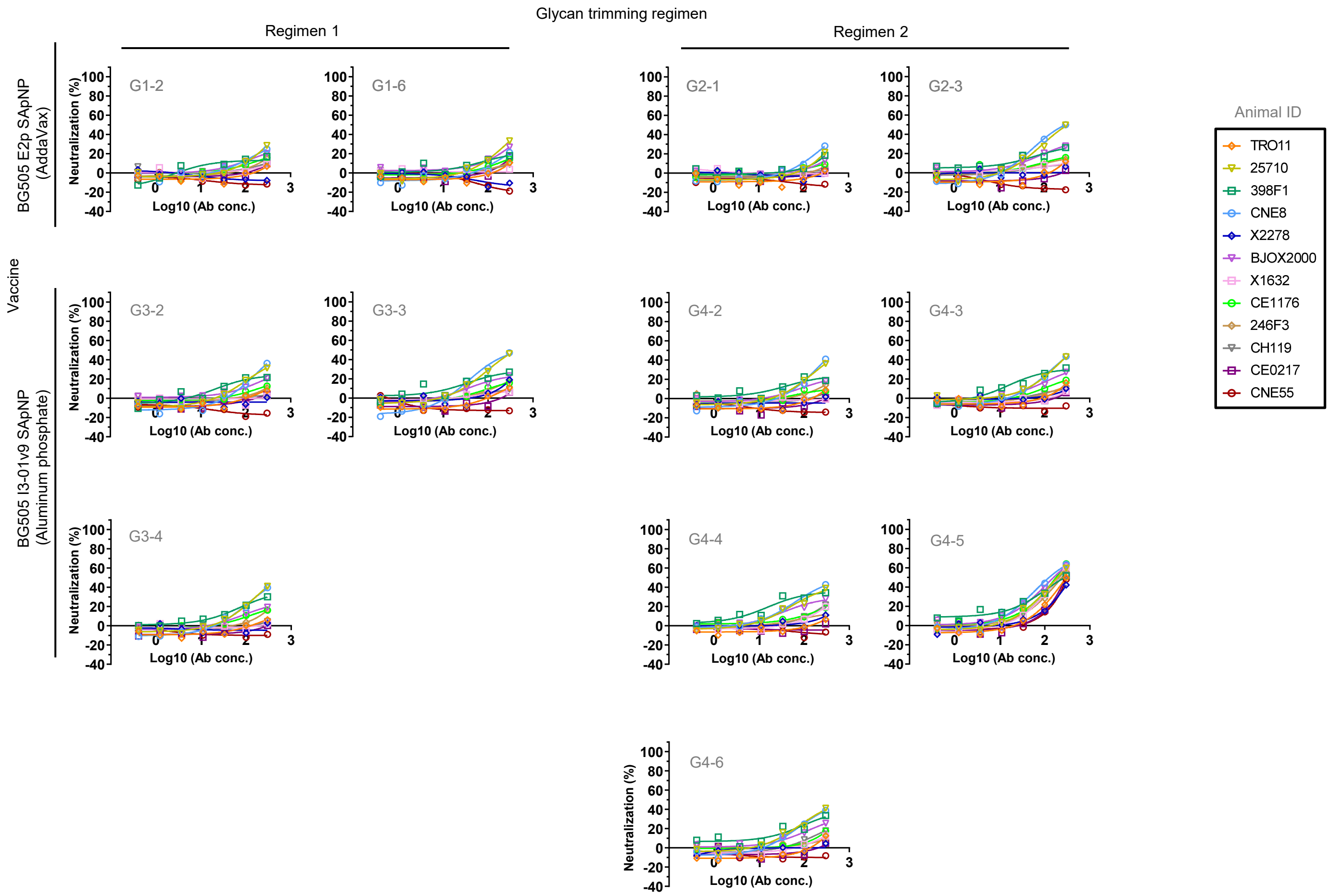

**fig. S6. Neutralization data from the evaluation of Env immunogens with trimmed glycans in mice and rabbits.** (**A**) Negative stain EM analysis to assess the structural integrity of SApNPs produced from 8 and 5 CHO-K1 cell clones that stably express E2p and I3-01v9 SApNPs, respectively. (**B**)-(**F**) Neutralization of purified IgG at week 11 from the mouse study. Groups of eight mice were immunized with BG505 Env immunogens mixed with AddaVax or aluminum phosphate adjuvants at weeks 0, 3, 6 and 9. Two glycan trimming regimens were tested: regimen 1 administers four doses of glycan-trimmed immunogens, whereas regimen 2 administers two doses of glycan-trimmed immunogens as prime and two doses of wildtype immunogens as boost. Immunization was repeated for some formulation/regimen groups to confirm the findings and are labeled in the summary. (**B**)-(**D**) Neutralization of week 11 IgG from mice immunized with (**B**) the BG505 UFO trimer and (**C**) UFO trimer-presenting SApNPs E2p and (**D**) I3-01v9 against autologous tier 2 clade A BG505.T332N, tier 1 clade B SF162, and MLV as a negative control. No IgG was available for one mouse in the UFO trimer/AP regimen 1 group and for two mice in the I3-01v9/AP regimen 1 (1st repeat) group, as the mice died during the immunization study. Of note, IgG samples from trimer/AP, E2p/AV, E2p/AP, and I3-01v9/AP in combination with regimens 1 and 2, a total of 8 groups, were further analyzed in (E) and (F). (**E**) Neutralization of week 11 IgG from select mice (≥ 30% autologous neutralization) in the abovementioned eight mouse groups against a BG505.T332N variant with the C3/V4 knockout mutation (I396R). $IC_{50}$ titers and neutralization curves are shown, along with BG505.T332N data from (B-D) for comparison. (**F**) Neutralization of week 11 IgG from two trimer groups and eight SApNP groups against a 12-virus global panel. $IC_{50}$ and $IC_{30}$ titers were calculated and are listed in two tables. For all the TZM-bl neutralization assays performed in (B)-(F), a starting IgG concentration of 300 μg/ml and a 3-times dilution series were used. Each IgG sample was run in duplicate in (B)-(D) except for some mice in the MLV and SF162 assays and all mice in the 12-virus assays in (F). Of note, mice for which IgG samples were no longer available at the time of the MLV or 12-virus assays are marked '-' in the $IC_{50}$/$IC_{30}$ titer tables. (**G**)-(**J**) Neutralization of purified IgG from the rabbit study. Four groups of rabbits (6 per group) were immunized to test the E2p/AV and I3-01v9/AP formulations for glycan trimming regimens 1 and 2, using a similar protocol to the mouse study. (**G**) Neutralization of week 0 rabbit IgG against BG505.T332N, SF162, and MLV. (**H**) Neutralization of week 11 rabbit IgG against BG505.T332N, SF162, and MLV. It should be noted that R5 IgG from the I3-01v9/AP regimen 2 group showed MLV neutralization and might contain non-specific antibodies. (**I**) Neutralization of select week 11 rabbit IgG (more than 30% autologous neutralization) against four BG505.T332N glycan hole mutant viruses. (**J**) Neutralization of select week 11 rabbit IgG (more than 30% autologous neutralization) against a 12-virus global panel. $IC_{50}$ and $IC_{30}$ titers were calculated and are listed in two tables with neutralization curves shown for each IgG sample against all 12 viruses. For all the TZM-bl neutralization assays performed in (G)-(J), a starting IgG concentration of 300 μg/ml and a 3-times dilution series were used. Each IgG sample was run in duplicate in (G-I) and in singlet in (J) due to the limited sample availability.

### fig. S7

**A** Week -2 RM serum neutralization against HIV-1 BG505.T332N. Immunization with wildtype BG505 SApNPs

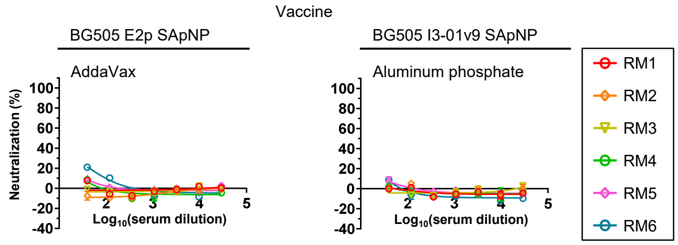


| Vaccine | Adjuvant | Week -2 $ID_{50}$ titers | | | | | |
|---|---|---|---|---|---|---|---|
| | | RM1 | RM2 | RM3 | RM4 | RM5 | RM6 |
| BG505 E2p SApNP | AddaVax | <40 | <40 | <40 | <40 | <40 | <40 |
| BG505 I3-01v9 SApNP | Aluminum phosphate | <40 | <40 | <40 | <40 | <40 | <40 |

Week -2 RM serum neutralization against HIV-1 SF162. Immunization with wildtype BG505 SApNPs

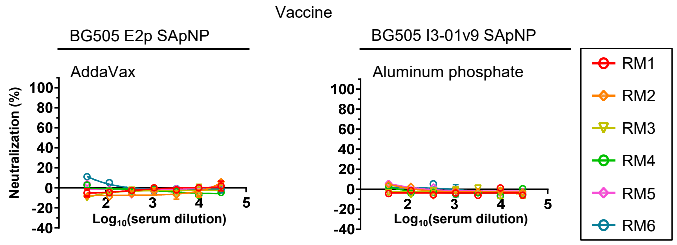


| Vaccine | Adjuvant | Week -2 $ID_{50}$ titers | | | | | |
|---|---|---|---|---|---|---|---|
| | | RM1 | RM2 | RM3 | RM4 | RM5 | RM6 |
| BG505 E2p SApNP | AddaVax | <40 | <40 | <40 | <40 | <40 | <40 |
| BG505 I3-01v9 SApNP | Aluminum phosphate | <40 | <40 | <40 | <40 | <40 | <40 |

Week -2 RM serum neutralization against MLV (control). Immunization with wildtype BG505 SApNPs

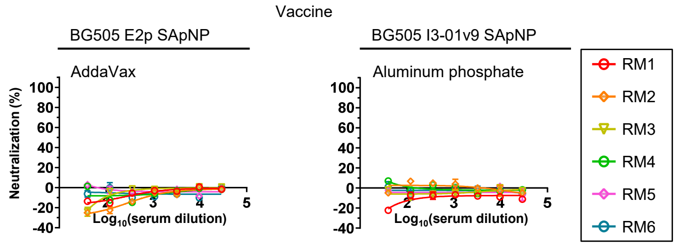


| Vaccine | Adjuvant | Week -2 $ID_{50}$ titers | | | | | |
|---|---|---|---|---|---|---|---|
| | | RM1 | RM2 | RM3 | RM4 | RM5 | RM6 |
| BG505 E2p SApNP | AddaVax | <40 | <40 | <40 | <40 | <40 | <40 |
| BG505 I3-01v9 SApNP | Aluminum phosphate | <40 | <40 | <40 | <40 | <40 | <40 |

**B** Week 28 RM serum neutralization against HIV-1 SF162. Immunization with wildtype BG505 SApNPs

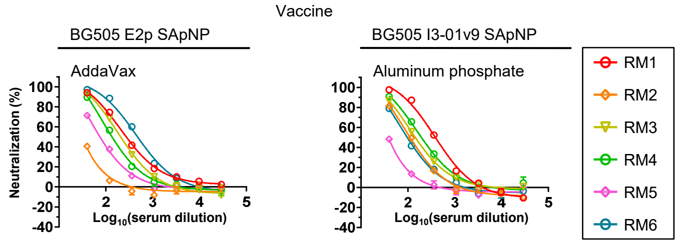


| Vaccine | Adjuvant | Week 28 $ID_{50}$ titers | | | | | |
|---|---|---|---|---|---|---|---|
| | | RM1 | RM2 | RM3 | RM4 | RM5 | RM6 |
| BG505 E2p SApNP | AddaVax | 300 | <40 | 210 | 144 | 74 | 487 |
| BG505 I3-01v9 SApNP | Aluminum phosphate | 382 | 108 | 152 | 203 | <40 | 96 |

Week 28 RM serum neutralization against MLV (control). Immunization with wildtype BG505 SApNPs

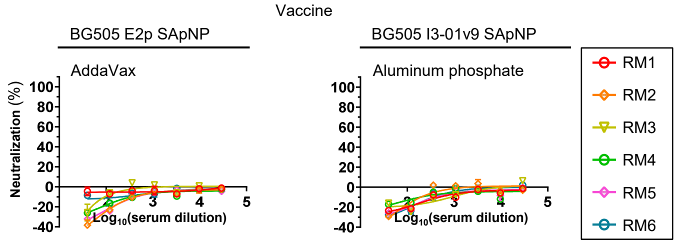


| Vaccine | Adjuvant | Week 28 $ID_{50}$ titers | | | | | |
|---|---|---|---|---|---|---|---|
| | | RM1 | RM2 | RM3 | RM4 | RM5 | RM6 |
| BG505 E2p SApNP | AddaVax | <40 | <40 | <40 | <40 | <40 | <40 |
| BG505 I3-01v9 SApNP | Aluminum phosphate | <40 | <40 | <40 | <40 | <40 | <40 |

**C** **Week 0 RM serum neutralization against HIV-1 BG505.T332N. Immunization with glycan trimmed BG505 SApNPs**

| Vaccine | Adjuvant | Glycan trimming regimen | Week 0 $ID_{50}$ titers | | | |
|---|---|---|---|---|---|---|
| | | | RM1 | RM2 | RM3 | RM4 |
| BG505 E2p SApNP | Aluminum phosphate | Regimen 1 | <40 | <40 | <40 | <40 |
| | | Regimen 2 | <40 | <40 | <40 | <40 |
| BG505 I3-01v9 SApNP | Aluminum phosphate | Regimen 1 | <40 | <40 | <40 | <40 |
| | | Regimen 2 | <40 | <40 | <40 | <40 |

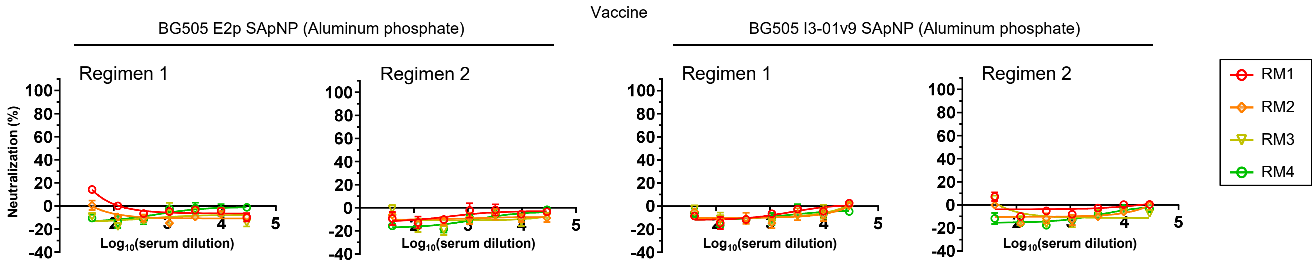


**Week 0 RM serum neutralization against HIV-1 SF162. Immunization with glycan trimmed BG505 SApNPs**

| Vaccine | Adjuvant | Glycan trimming regimen | Week 0 $ID_{50}$ titers | | | |
|---|---|---|---|---|---|---|
| | | | RM1 | RM2 | RM3 | RM4 |
| BG505 E2p SApNP | Aluminum phosphate | Regimen 1 | <40 | <40 | <40 | <40 |
| | | Regimen 2 | <40 | <40 | <40 | <40 |
| BG505 I3-01v9 SApNP | Aluminum phosphate | Regimen 1 | <40 | <40 | <40 | <40 |
| | | Regimen 2 | <40 | <40 | <40 | <40 |

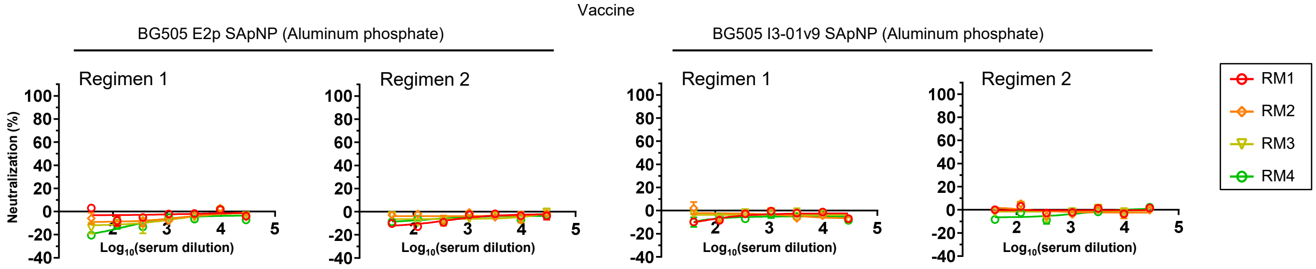


**Week 0 RM serum neutralization against MLV (control). Immunization with glycan trimmed BG505 SApNPs**

| Vaccine | Adjuvant | Glycan trimming regimen | Week 0 $ID_{50}$ titers | | | |
|---|---|---|---|---|---|---|
| | | | RM1 | RM2 | RM3 | RM4 |
| BG505 E2p SApNP | Aluminum phosphate | Regimen 1 | <40 | <40 | <40 | <40 |
| | | Regimen 2 | <40 | <40 | <40 | <40 |
| BG505 I3-01v9 SApNP | Aluminum phosphate | Regimen 1 | <40 | <40 | <40 | <40 |
| | | Regimen 2 | <40 | <40 | <40 | <40 |

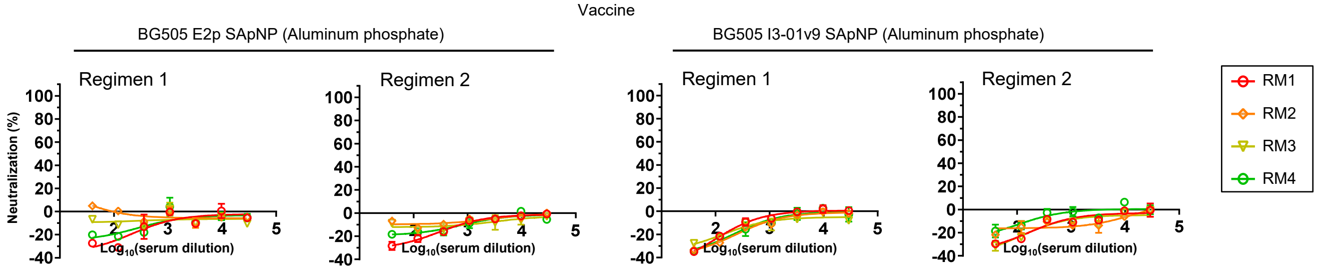

D **Week 28 RM serum neutralization against HIV-1 SF162. Immunization with glycan trimmed BG505 SApNPs**

| Vaccine | Adjuvant | Glycan trimming regimen | Week 28 $ID_{50}$ titers | | | |
|---|---|---|---|---|---|---|
| | | | RM1 | RM2 | RM3 | RM4 |
| BG505 E2p SApNP | Aluminum phosphate | Regimen 1 | <40 | <40 | <40 | 206 |
| | | Regimen 2 | 65 | 261 | 85 | 130 |
| BG505 I3-01v9 SApNP | Aluminum phosphate | Regimen 1 | 205 | <40 | 99 | 57 |
| | | Regimen 2 | 117 | 40 | 59 | <40 |

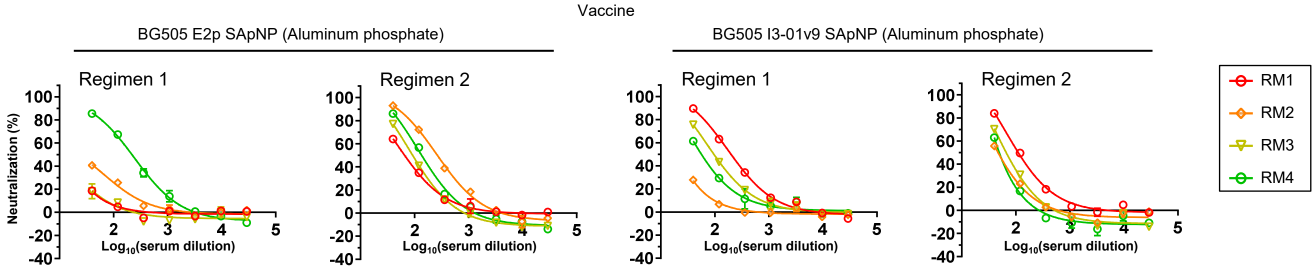


**Week 28 RM serum neutralization against MLV (control). Immunization with glycan trimmed BG505 SApNPs**

| Vaccine | Adjuvant | Glycan trimming regimen | Week 28 $ID_{50}$ titers | | | |
|---|---|---|---|---|---|---|
| | | | RM1 | RM2 | RM3 | RM4 |
| BG505 E2p SApNP | Aluminum phosphate | Regimen 1 | <40 | <40 | <40 | <40 |
| | | Regimen 2 | <40 | <40 | <40 | <40 |
| BG505 I3-01v9 SApNP | Aluminum phosphate | Regimen 1 | <40 | <40 | <40 | <40 |
| | | Regimen 2 | <40 | <40 | <40 | <40 |

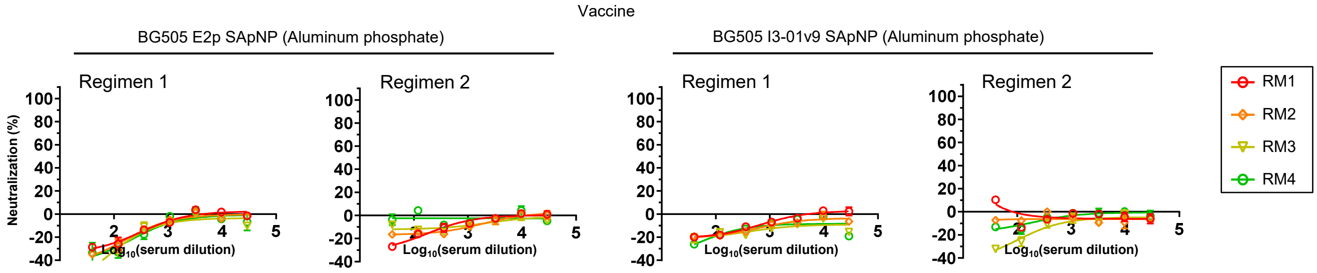


**fig. S7. Neutralization data from the evaluation of BG505 SApNP immunogens with wildtype and trimmed glycans in NHPs.** (**A**)-(**B**) Serum neutralization at weeks -2 and 28 from the immunization study of wildtype SApNPs. Two groups of six rhesus macaques were immunized with wildtype E2p and I3-01v9 SApNPs (formulated with the AV and AP adjuvants, respectively) at weeks 0, 8, and 24 with 13 blood draws. (**A**) Week -2 serum neutralization against autologous tier 2 clade A BG505.T332N, tier 1 clade B SF162, and MLV (negative control). (**B**) Week 28 serum neutralization against SF162 and MLV. (**C**)-(**D**) Serum neutralization at weeks 0 and 28 from the immunization study of glycan-trimmed SApNPs. Four groups of four rhesus macaques were used to test two SApNPs (E2p and I3-01v9) in combination two regimens. Subjects in regimen 1 (homologous) received four doses of glycan-trimmed SApNPs, whereas subjects in regimen 2 (heterologous) received two doses of glycan-trimmed immunogens as a prime and two doses of wildtype immunogens as a boost. SApNPs in both regimens were formulated with the AP adjuvant and administered at weeks 0, 4, 12, and 24 with 9 blood draws. (**C**) Week 0 serum neutralization against autologous tier 2 clade A BG505.T332N, tier 1 clade B SF162, and MLV (negative control). (**D**) Week 28 serum neutralization data against SF162 and MLV. $ID_{50}$ titers and %neutralization curves are shown in (A)-(D).

fig. S8

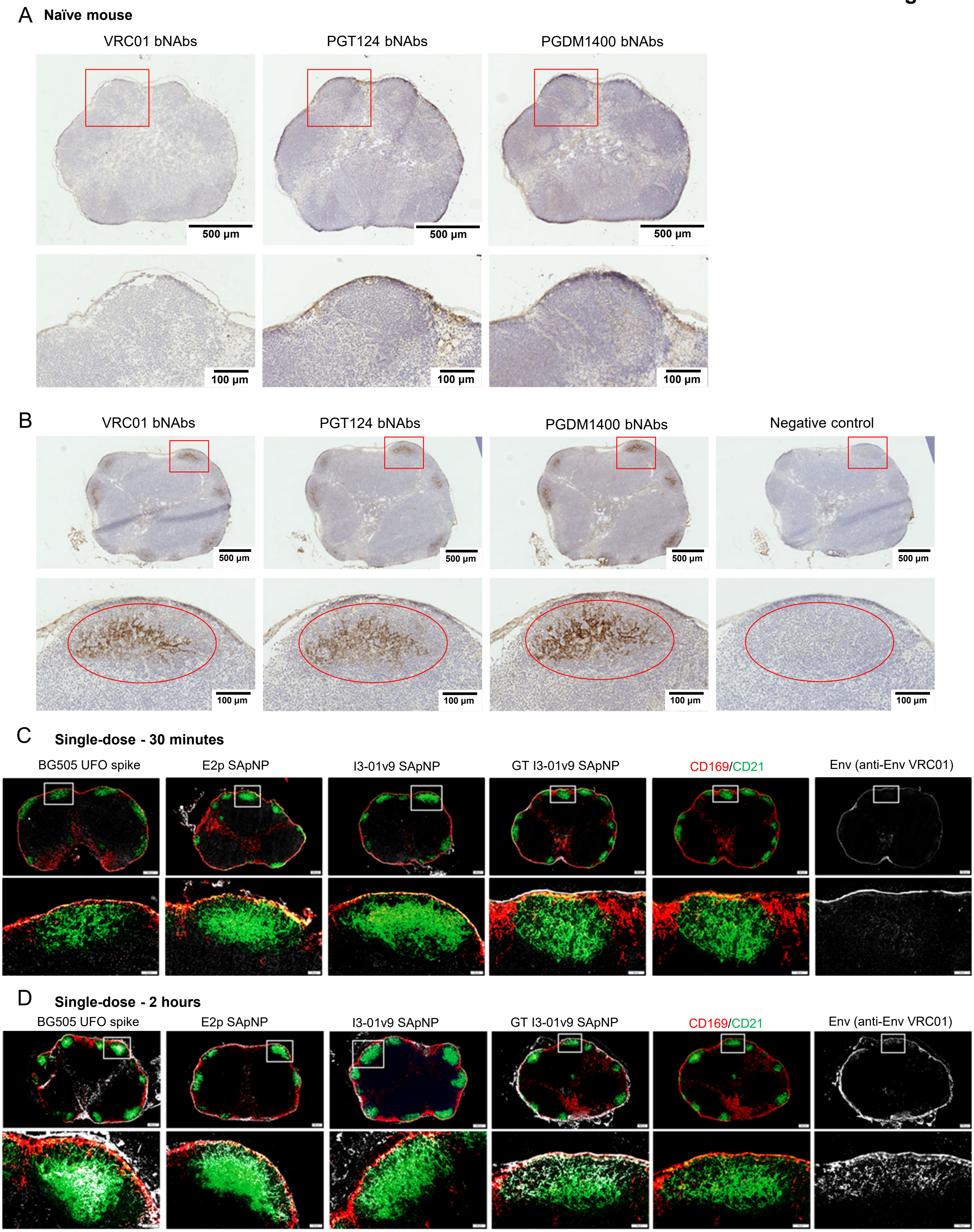
A Naïve mouse
VRC01 bNAbs
PGT124 bNAbs
PGDM1400 bNAbs
500 µm
500 µm
500 µm
100 µm
100 µm
100 µm
B
VRC01 bNAbs
PGT124 bNAbs
PGDM1400 bNAbs
Negative control
500 µm
500 µm
500 µm
500 µm
100 µm
100 µm
100 µm
100 µm
C Single-dose - 30 minutes
BG505 UFO spike
E2p SApNP
I3-01v9 SApNP
GT I3-01v9 SApNP
CD169/CD21
Env (anti-Env VRC01)
D Single-dose - 2 hours
BG505 UFO spike
E2p SApNP
I3-01v9 SApNP
GT I3-01v9 SApNP
CD169/CD21
Env (anti-Env VRC01)

fig. S8

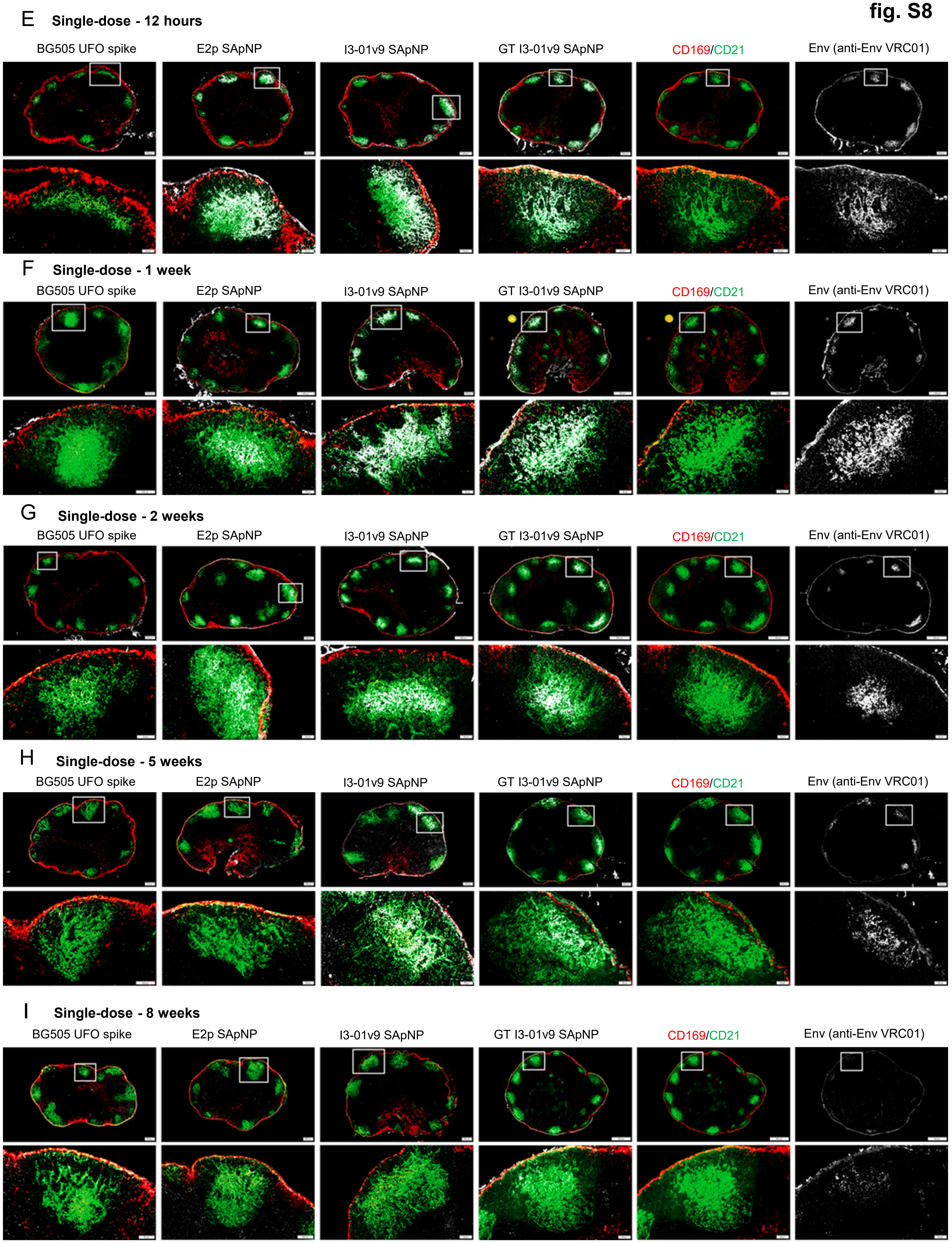
E Single-dose - 12 hours
BG505 UFO spike
E2p SApNP
I3-01v9 SApNP
GT I3-01v9 SApNP
CD169/CD21
Env (anti-Env VRC01)
F Single-dose - 1 week
BG505 UFO spike
E2p SApNP
I3-01v9 SApNP
GT I3-01v9 SApNP
CD169/CD21
Env (anti-Env VRC01)
G Single-dose - 2 weeks
BG505 UFO spike
E2p SApNP
I3-01v9 SApNP
GT I3-01v9 SApNP
CD169/CD21
Env (anti-Env VRC01)
H Single-dose - 5 weeks
BG505 UFO spike
E2p SApNP
I3-01v9 SApNP
GT I3-01v9 SApNP
CD169/CD21
Env (anti-Env VRC01)
I Single-dose - 8 weeks
BG505 UFO spike
E2p SApNP
I3-01v9 SApNP
GT I3-01v9 SApNP
CD169/CD21
Env (anti-Env VRC01)

**fig. S8. Immunohistological images of BG505 UFO trimer and SApNPs in lymph nodes.** Immunostaining images of lymph node tissues from (**A**) naïve and (**B**) BG505 UFO-presenting SApNP injected mice stained with three anti-trimer antibody VRC01, PGT124 and PGDM1400. HIV-1 vaccine constructs of BG505 UFO trimer and BG505 UFO trimer-presenting E2p, I3-01v9, and GT I3-01v9 SApNPs interaction with FDC networks in lymph node follicles (**C**) 30 minutes, (**D**) 2 hours, (**E**) 12 hours, (**F**) 1 week, (**G**) 2 weeks, (**H**) 5 weeks, and (**I**) 8 weeks after a single-dose injection (10 μg per injection, totaling 40 μg per mouse). Immunofluorescent images are pseudo color coded (CD21+, green; CD169+, red; anti-Env VRC01, White), with a complete lymph node and an enlarged image of a follicle.

**fig. S9**

A E2p SApNPs (yellow arrow) and AddaVax (green arrow)
are aligned on FDC dendrites after 2 hours of injection

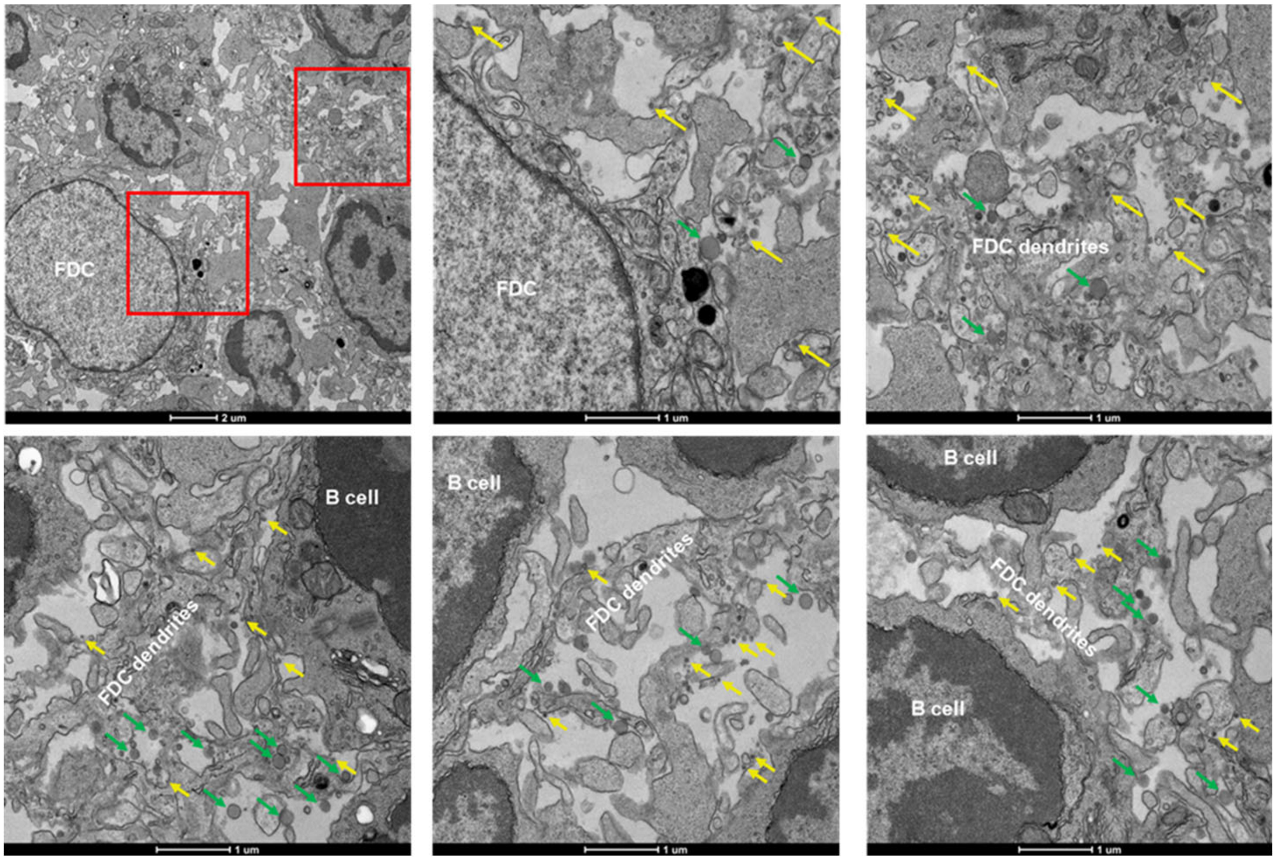


B E2p SApNPs (yellow arrow) and AddaVax (green arrow)
are aligned on FDC dendrites after 12 hours of injection

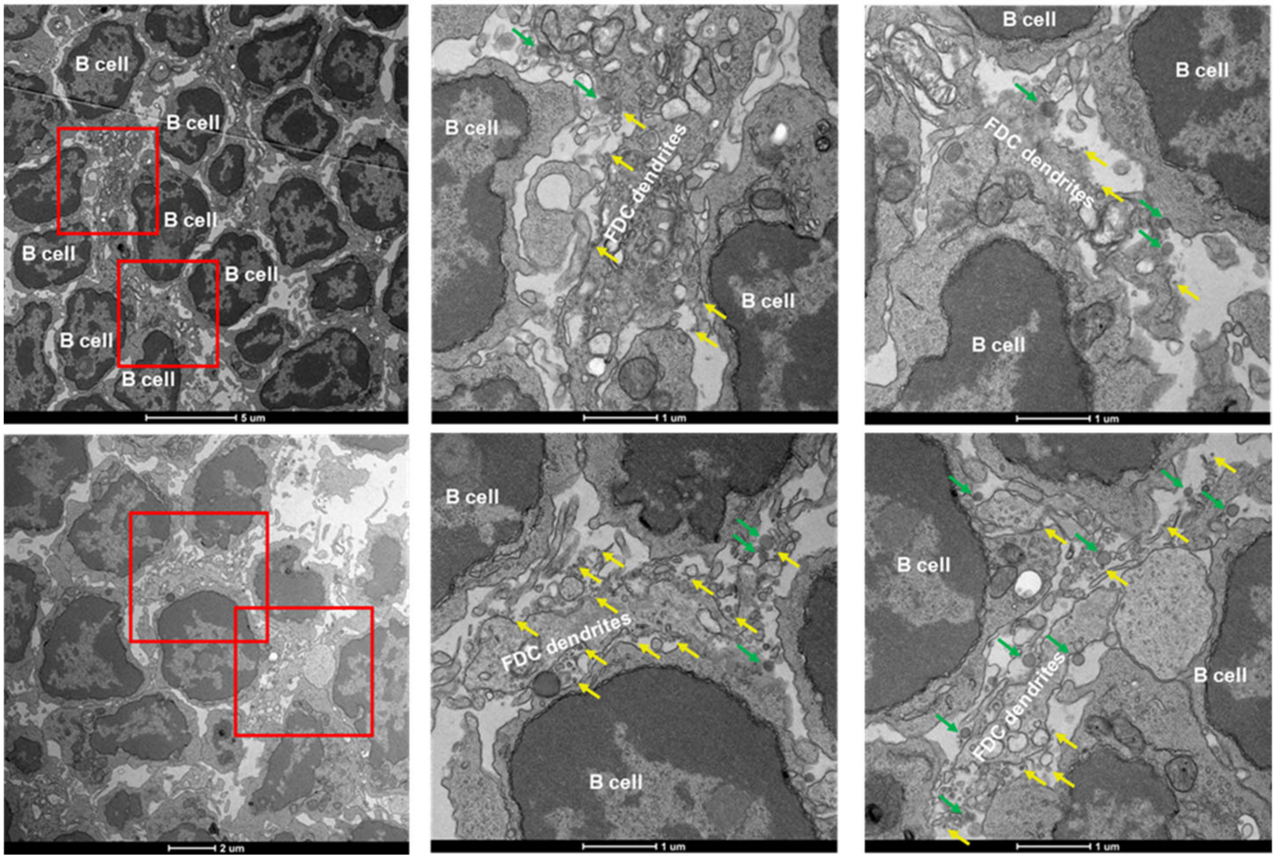

fig. S9

C E2p SApNPs (yellow arrow) are aligned on FDC dendrites or taken up by B cells after 48 hours of injection

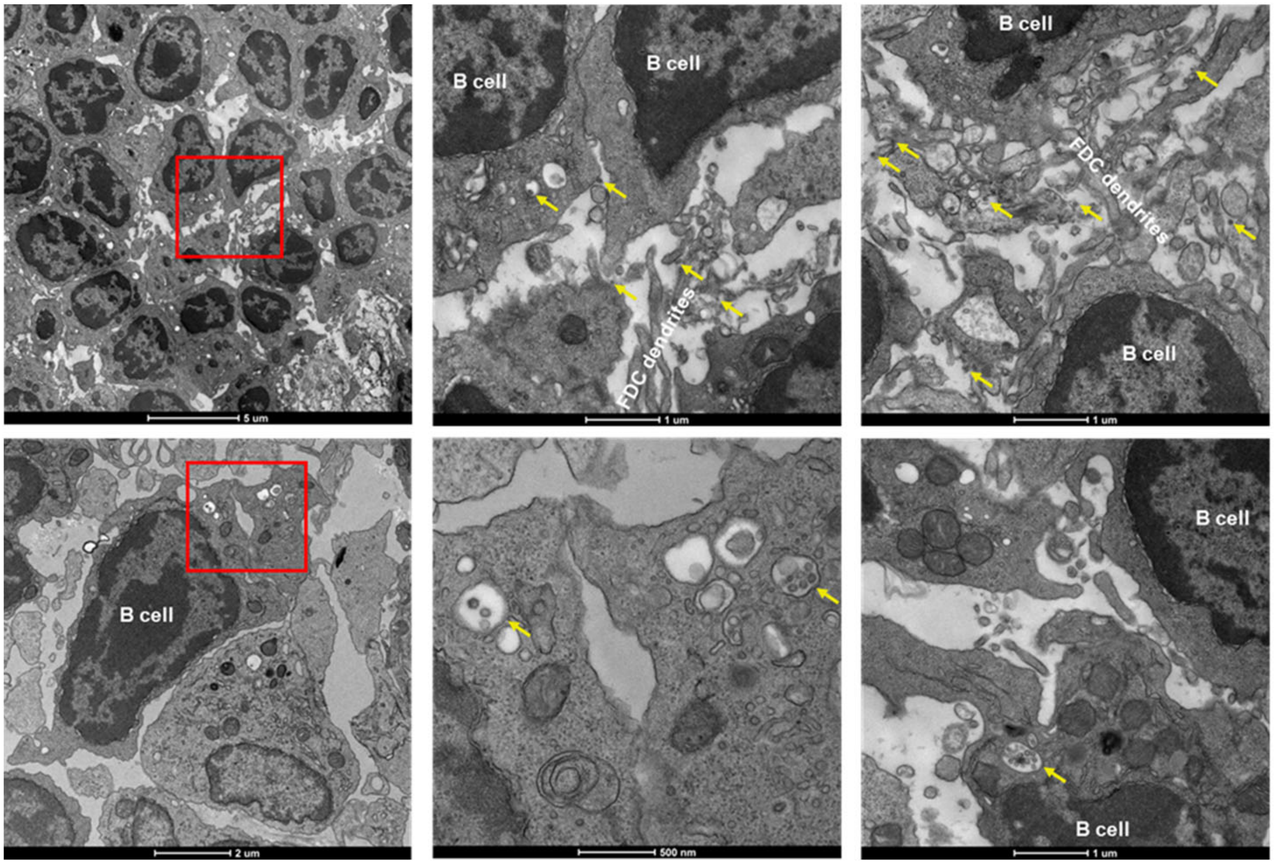


D I3-01v9 SApNPs (yellow arrow) are aligned on FDC dendrites after 2 hours of injection

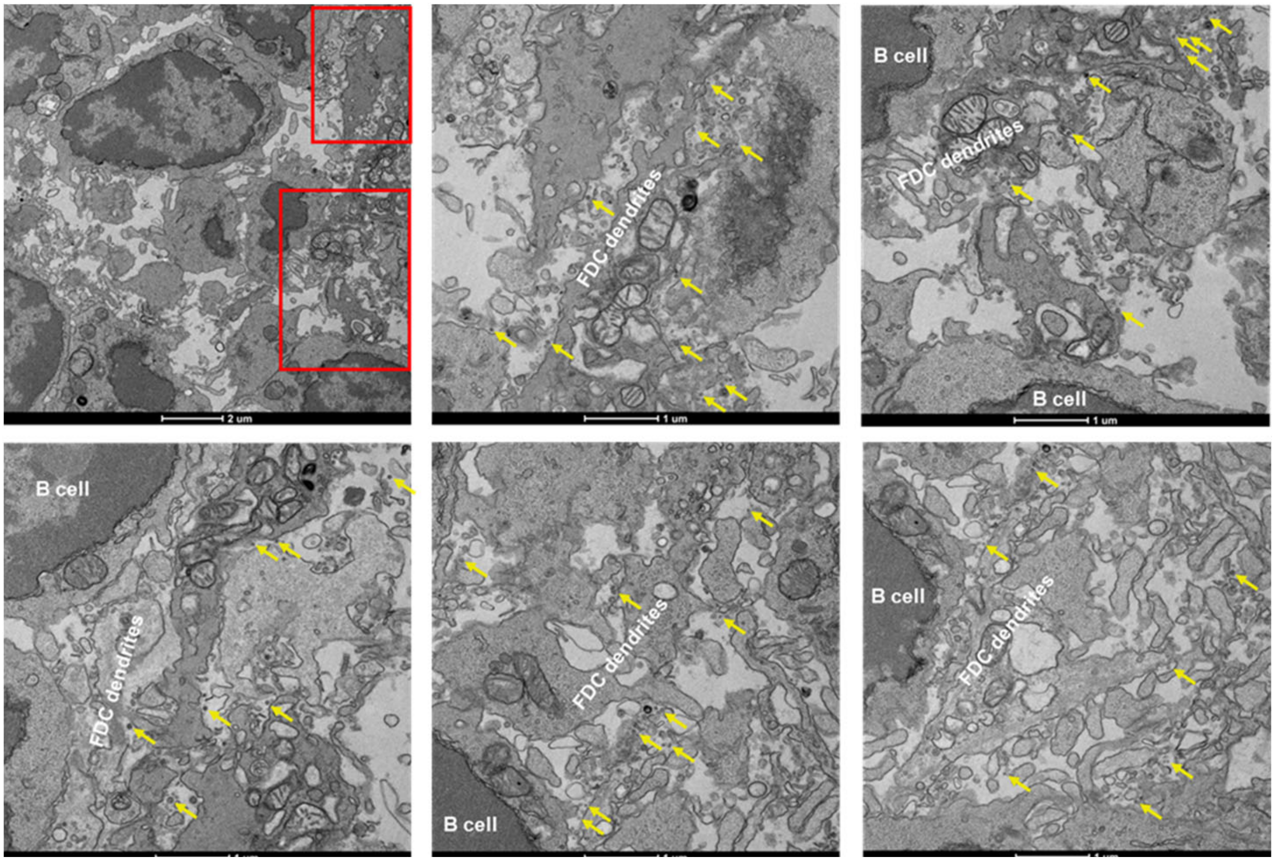

E I3-01v9 SApNPs (yellow arrow) are aligned on FDC dendrites after 12 hours of injection

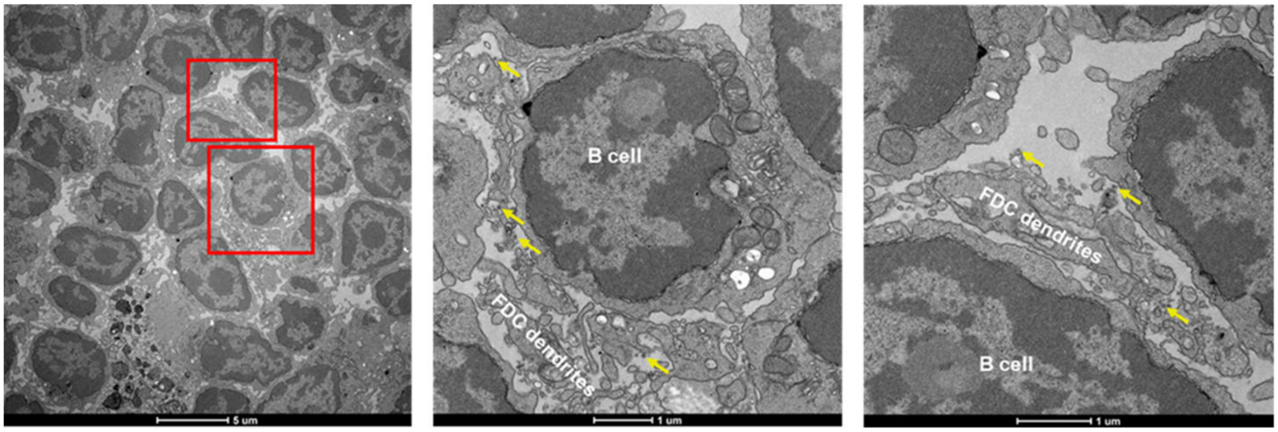


F I3-01v9 SApNPs (yellow arrow) are aligned on FDC dendrites after 48 hours of injection

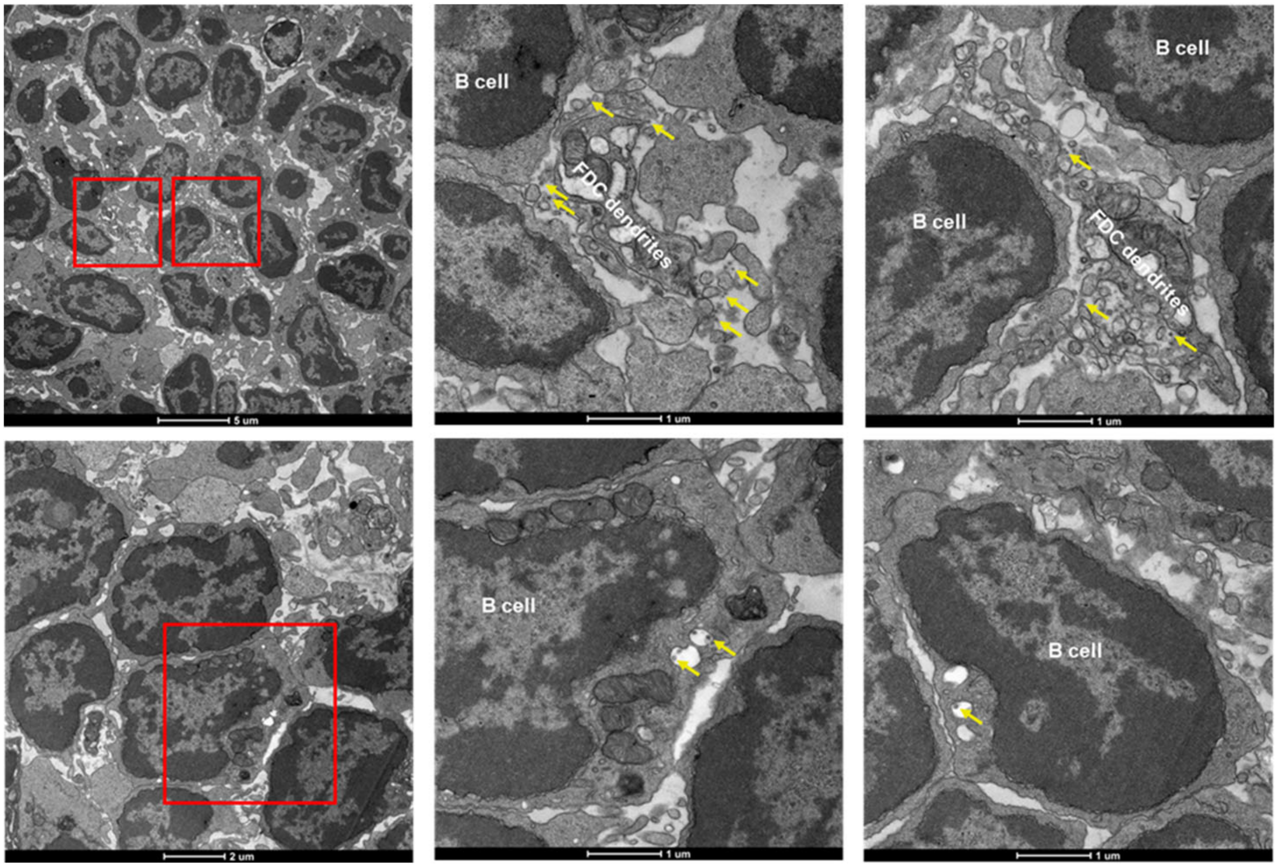

**fig. S9**

G GT I3-01v9 SApNPs (yellow arrow) are aligned on FDC dendrites after 2 hours of injection

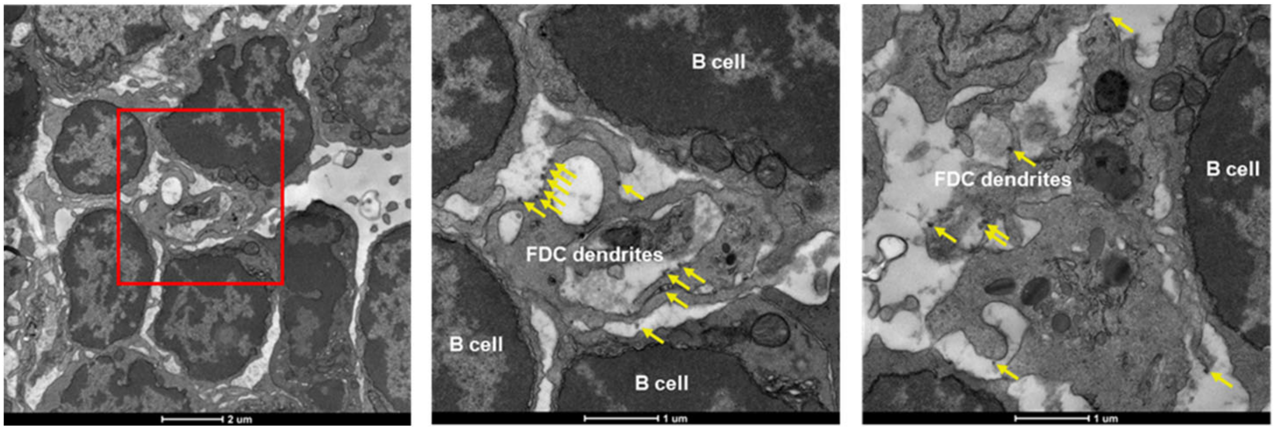


H GT I3-01v9 SApNPs (yellow arrow) are aligned on FDC dendrites after 12 hours of injection

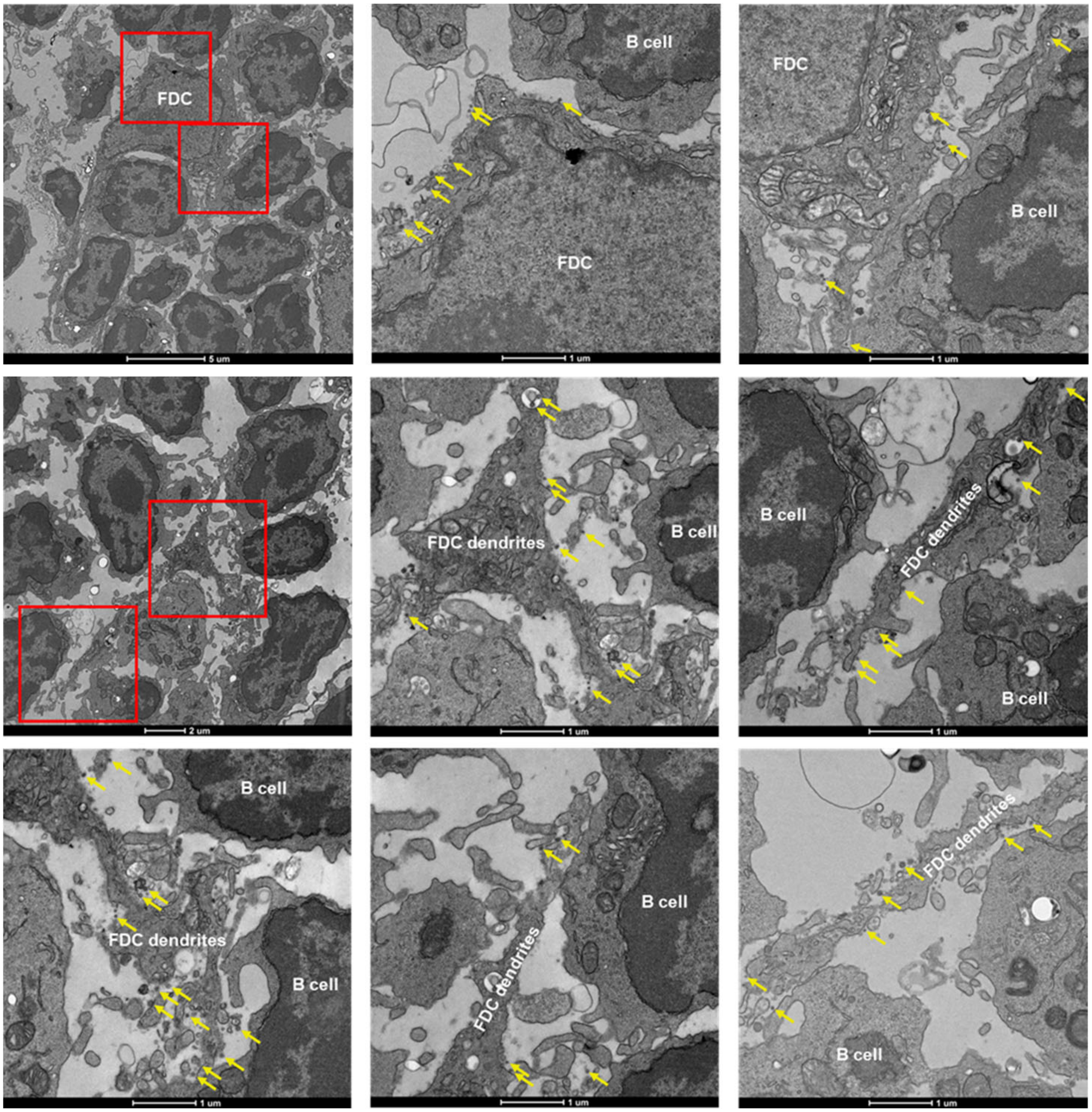

fig. S9

I GT I3-01v9 SApNPs (yellow arrow) are aligned on FDC dendrites after 48 hours of injection

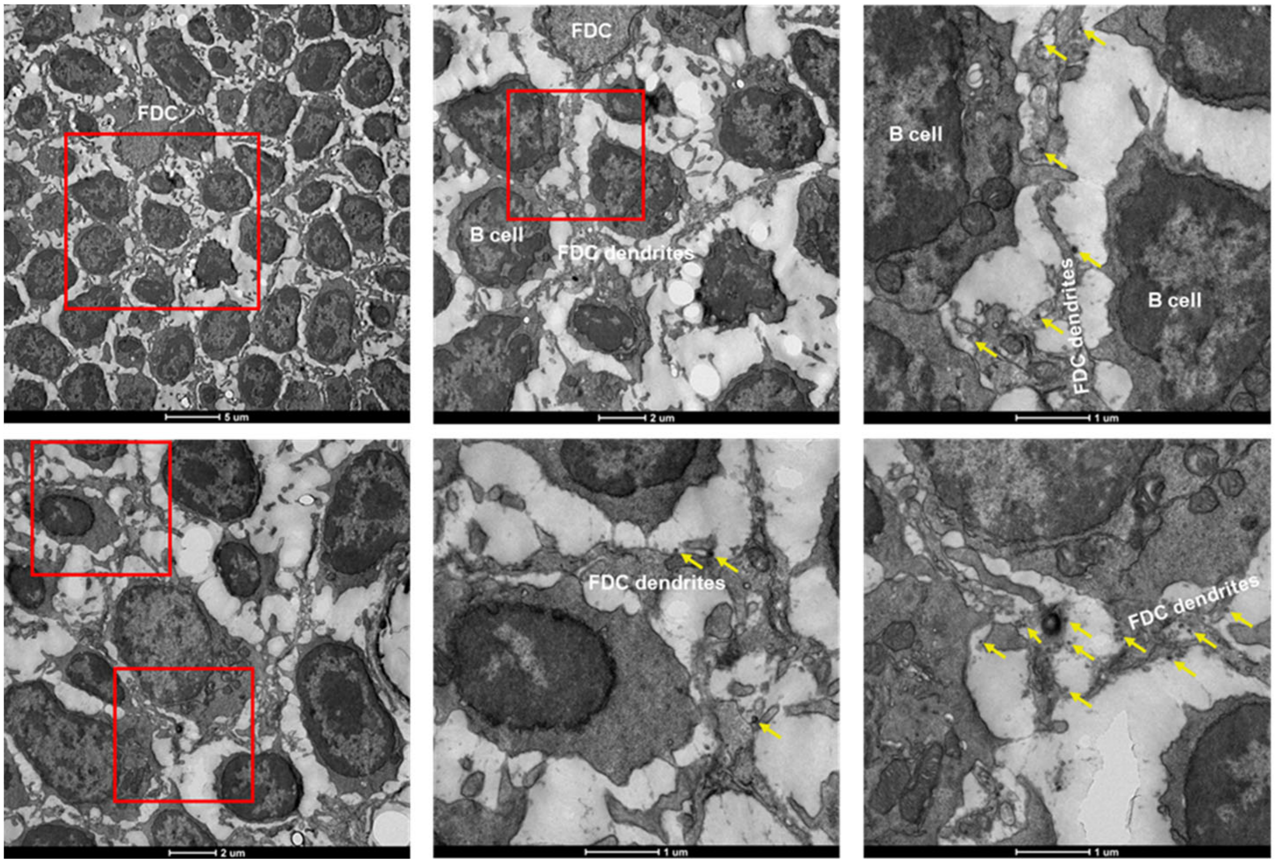


J E2p SApNPs (yellow arrow) alone are aligned on FDC dendrites after 12 hours of injection

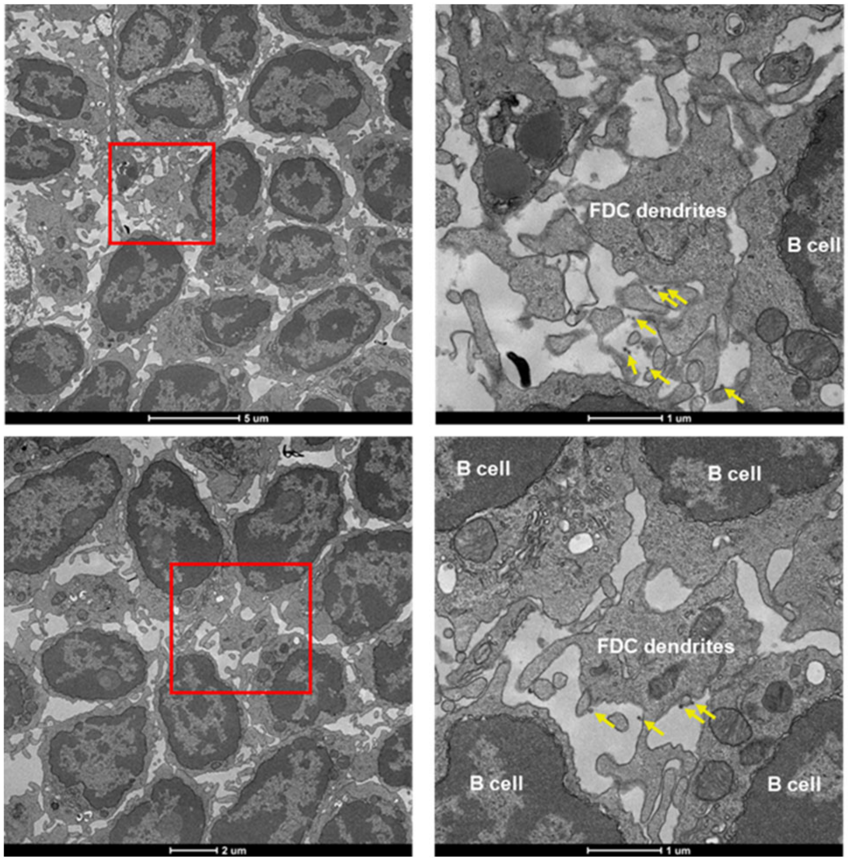

K I3-01v9 SApNPs alone (yellow arrow) are aligned on FDC dendrites after 12 hours of injection

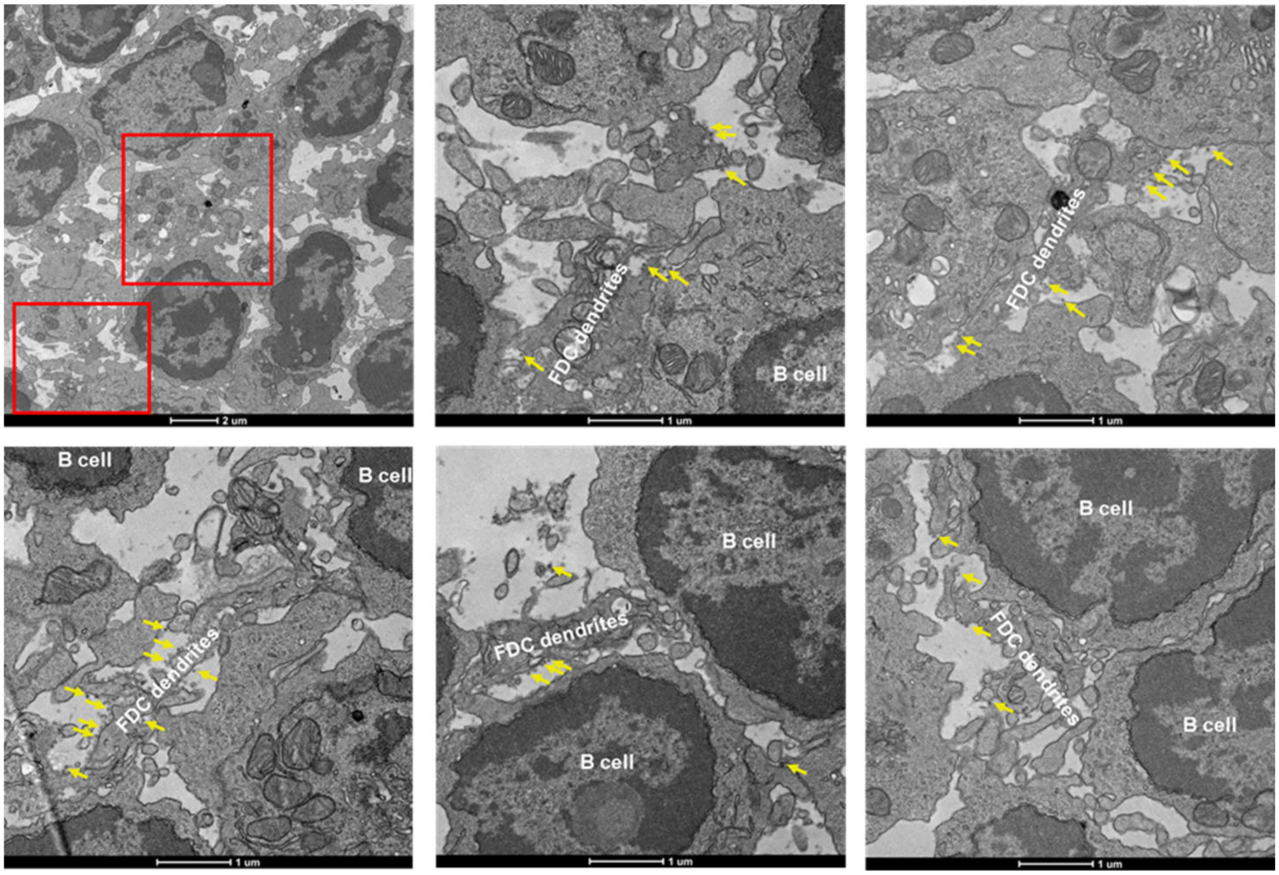


L AddaVax (green arrow) alone are aligned on FDC dendrites after 12 hours of injection

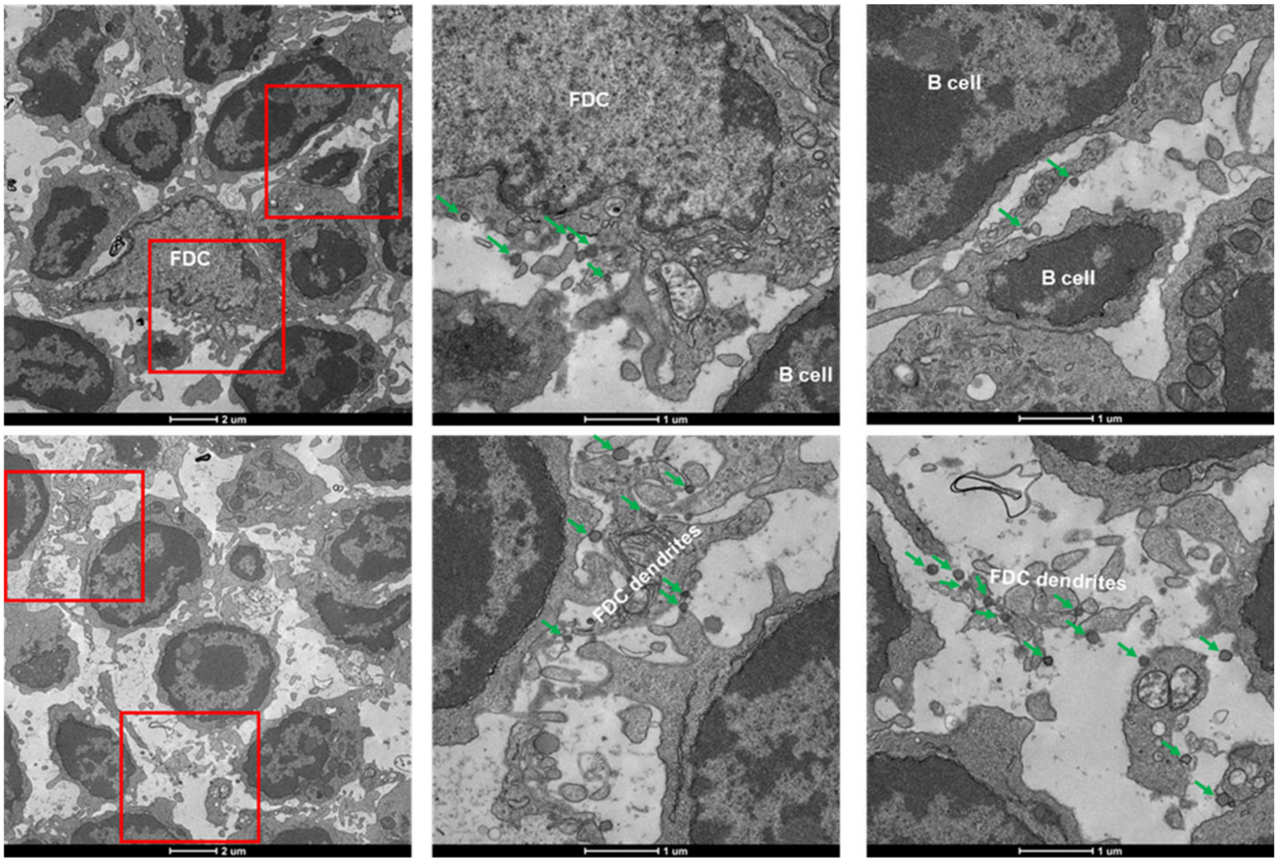

fig. S9

M FDC dendrites in Lymph nodes from naïve mouse

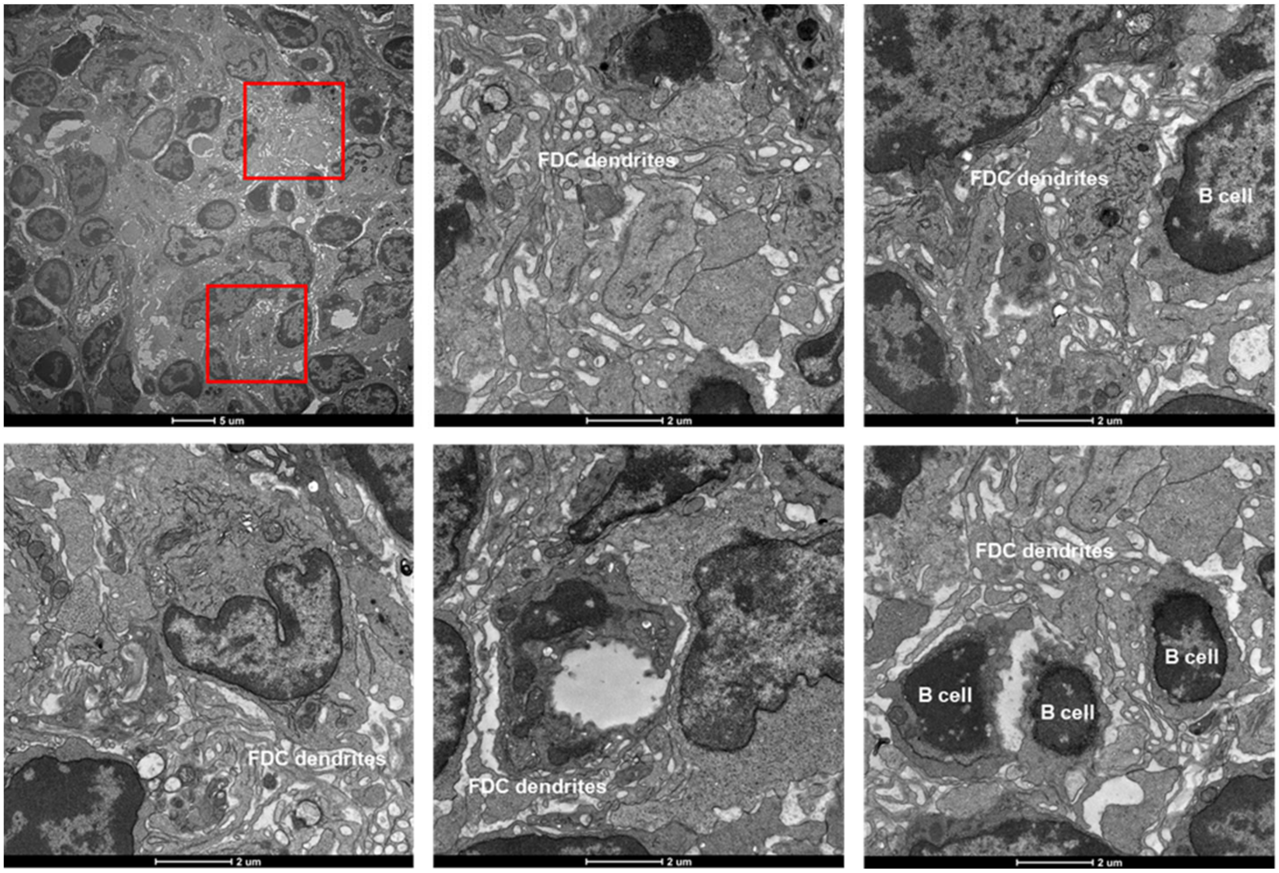


N E2p SApNPs (yellow arrow) and AddaVax (green arrow) are on the surface or inside the endolysosomes of phagocytic cells after 2 hours of injection

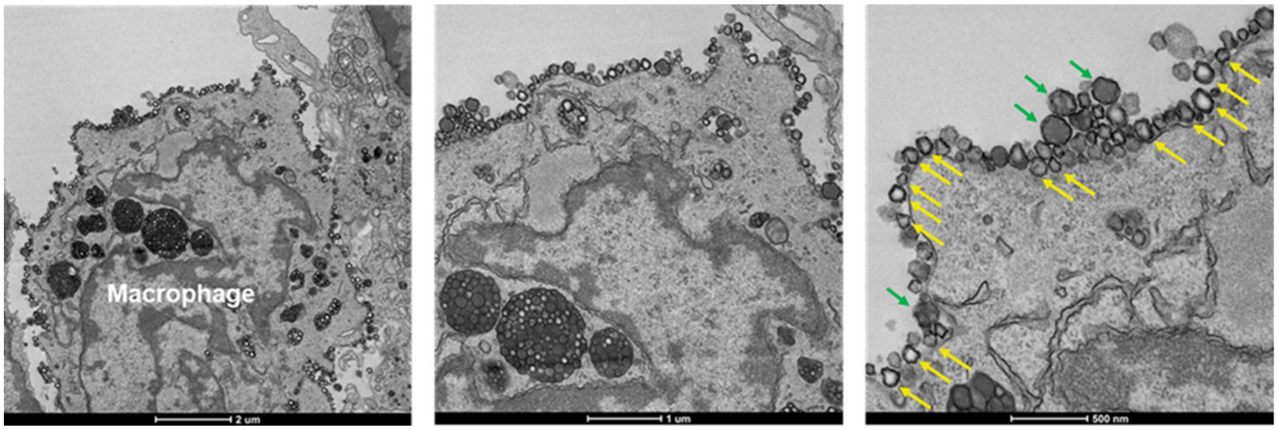

O I3-01v9 SApNPs (yellow arrow) are associated with phagocytic cells after 2 hours of injection

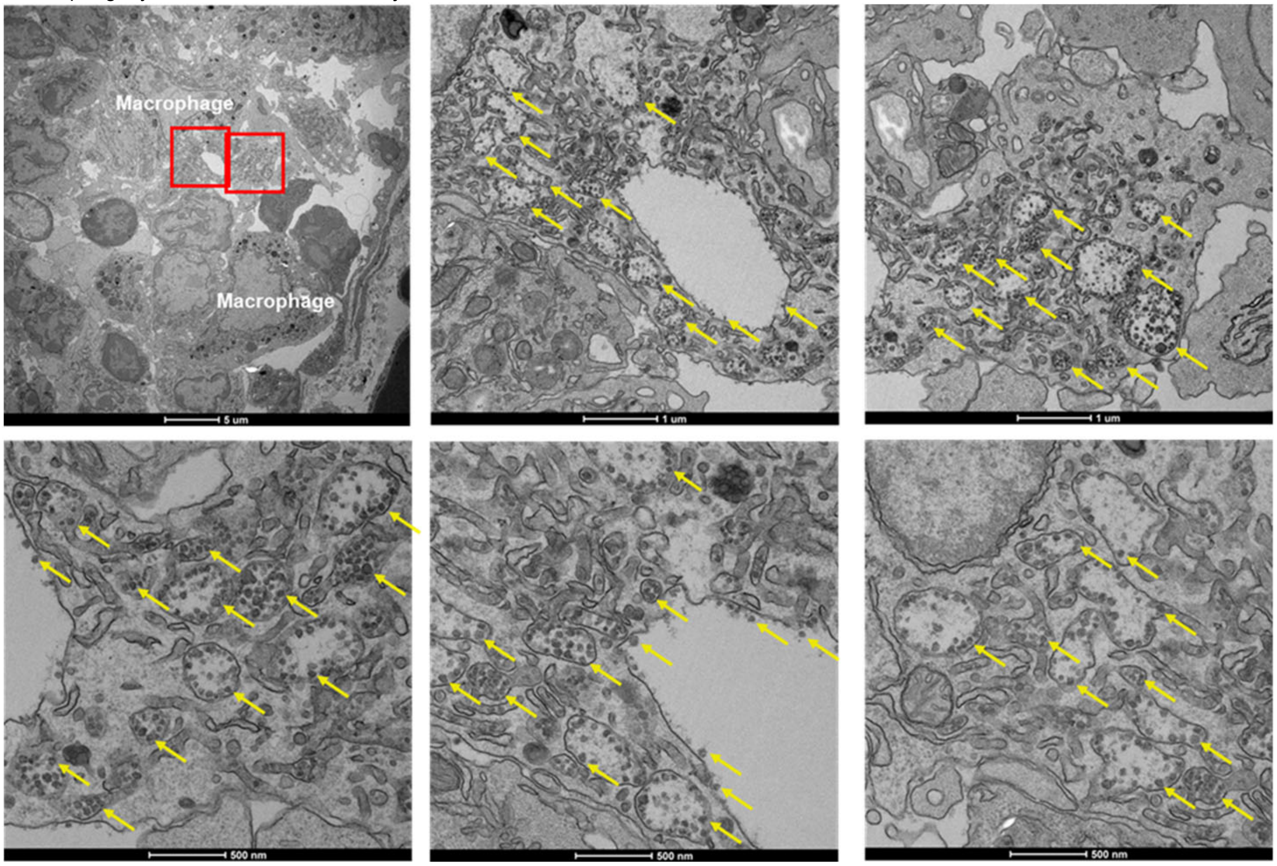


P I3-01v9 SApNPs (yellow arrow) are associated with phagocytic cells after 12 hours of injection

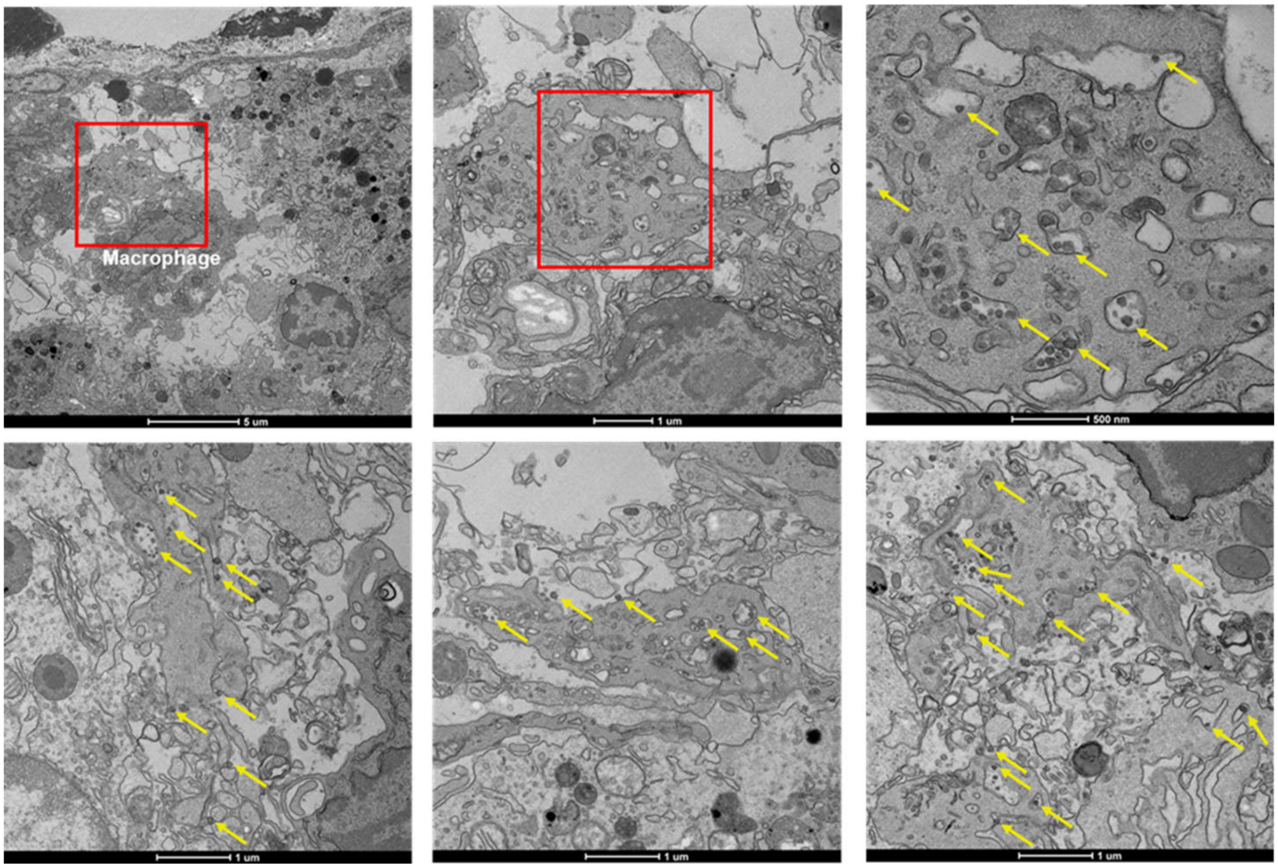

Q I3-01v9 SApNPs (yellow arrow) are associated with phagocytic cells after 48 hours of injection

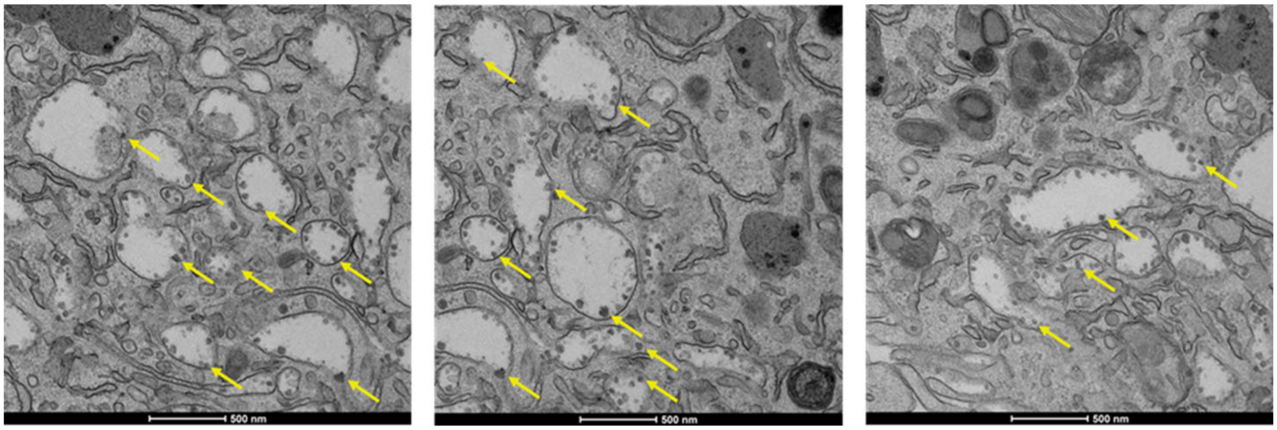


R GT I3-01v9 SApNPs (yellow arrow) are associated with phagocytic cells after 2 hours of injection

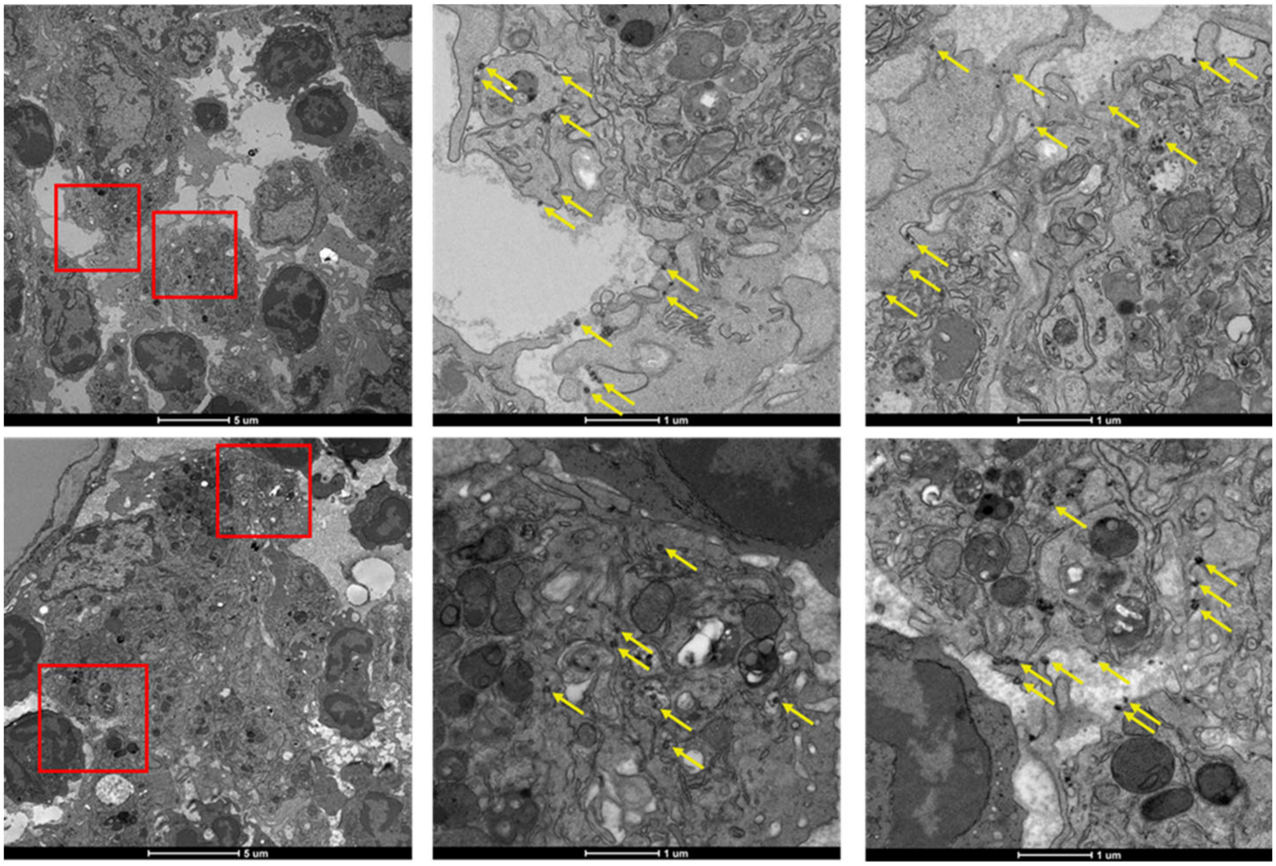

**fig. S9**

S GT I3-01v9 SApNPs (yellow arrow) and aluminum phosphate (green arrow) are associated with phagocytic cells after 12 hours of injection

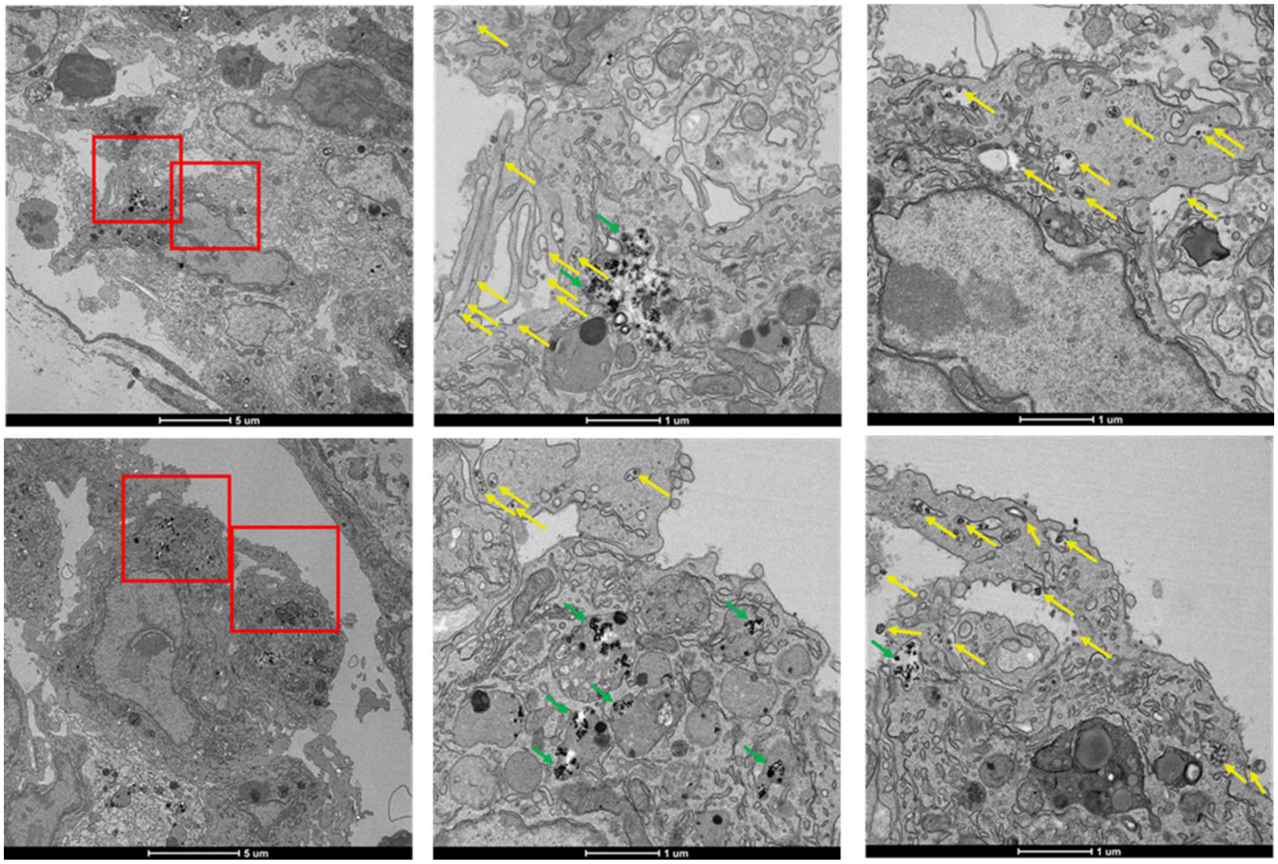


T GT I3-01v9 SApNPs (yellow arrow) are associated with phagocytic cells after 48 hours of injection

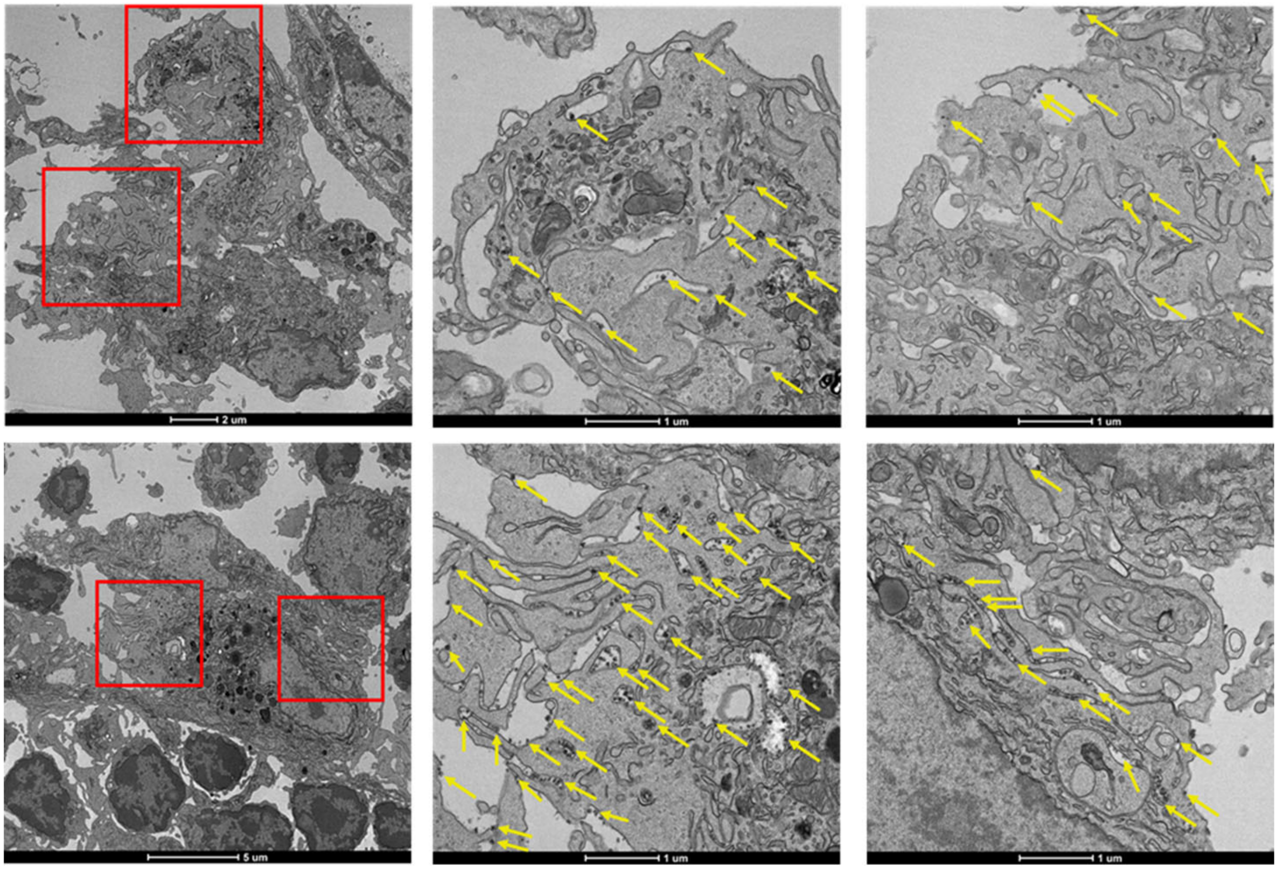

**fig. S9**

U Aluminum phosphate (green arrow) alone tends to aggregate and are observed inside the phagocytic cells, or in the ECM

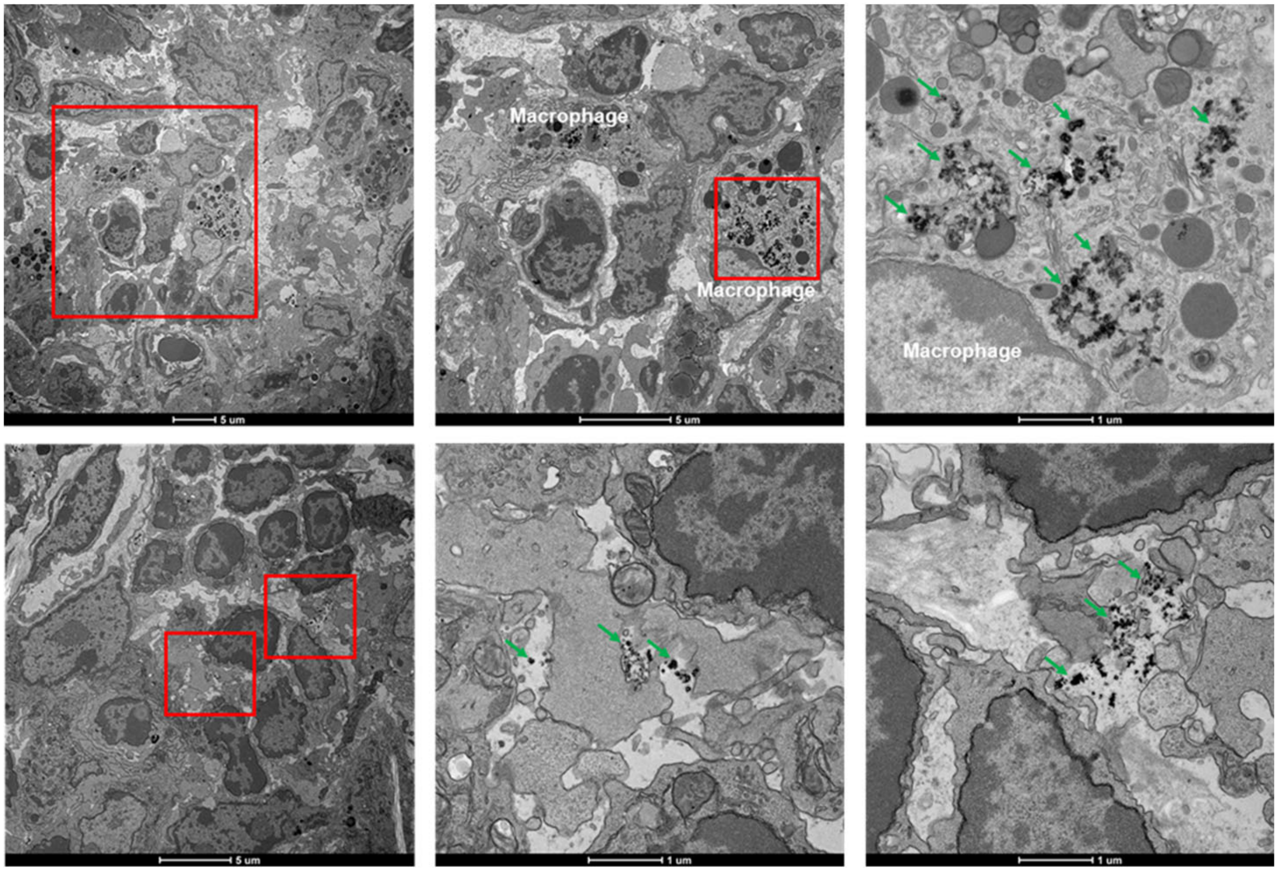


V Phagocytic cells in Lymph nodes from naïve mouse

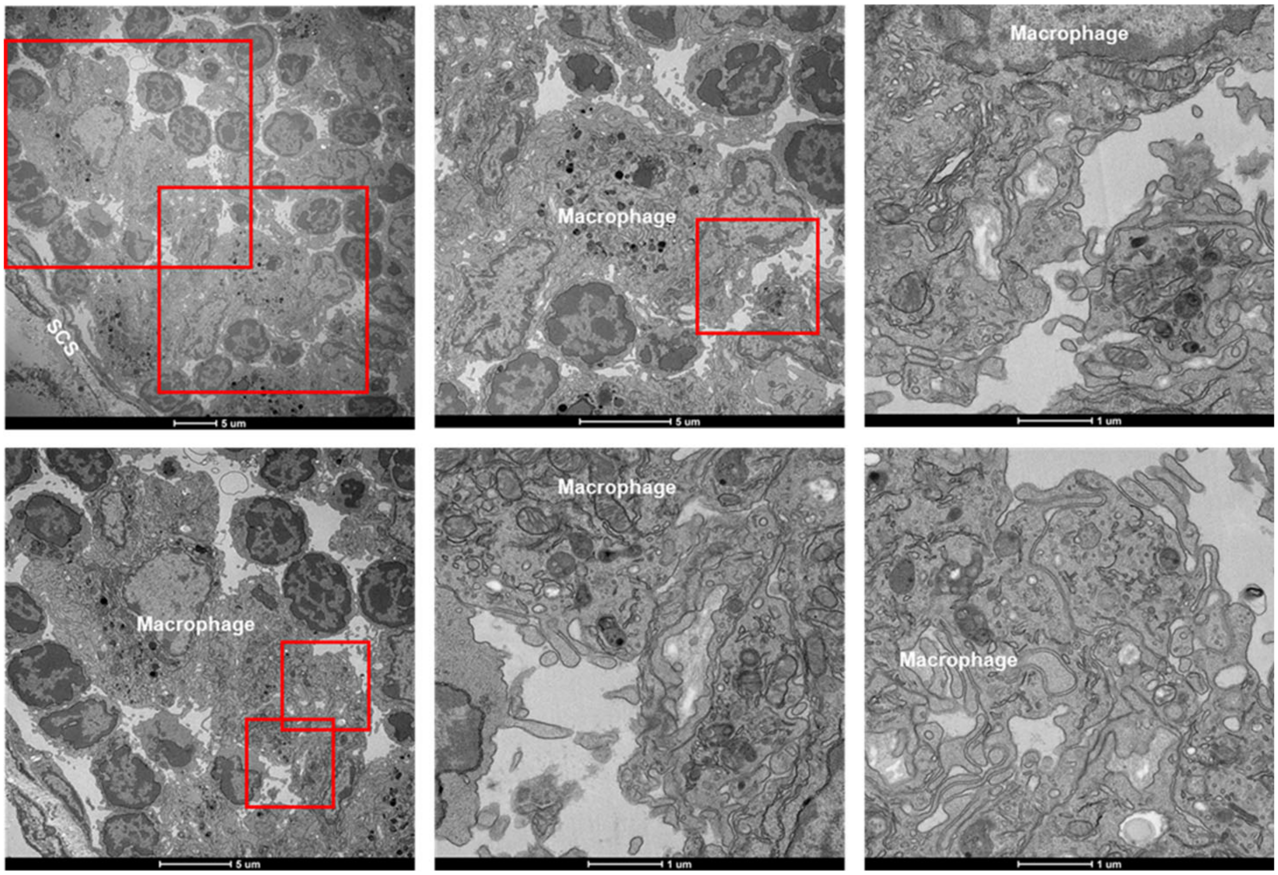

**fig. S9. TEM images of HIV-1 trimer/trimer-presenting SApNP interaction with FDCs and phagocytic cells in a lymph node.** BG505 UFO trimer-presenting E2p SApNPs formulated with AddaVax adjuvant are aligned on FDC dendrites (**A**) 2 hours, (**B**) 12 hours, and (**C**) 48 hours after a single-dose injection (50 µg). Adjuvanted BG505 UFO trimer-presenting (**D-F**) I3-01v9 and (**G-I**) GT I3-01v9 SApNPs are aligned on FDC dendrites 2, 12, and 48 hours after a single-dose injection (50 µg), whereas aluminum phosphate adjuvants were not associated with SApNPs on FDC dendrites. (**J**) BG505 UFO trimer-presenting E2p, (**K**) I3-01v9 SApNPs, or (**L**) AddaVax adjuvant alone can be presented on FDC dendrites 12 hours after injection. SApNPs are pointed by yellow arrows and adjuvants are pointed by green arrows. (**M**) FDC dendrites in Lymph nodes from naïve mouse. (**N**) BG505 UFO trimer-presenting E2p SApNPs formulated with AddaVax adjuvant are aligned on the surface or inside the endolysosomes of medullary sinus macrophages 2 hours after injection. Adjuvanted BG505 UFO trimer-presenting (**O-Q**) I3-01v9 and (**R-T**) GT I3-01v9 SApNPs are on macrophage surface or internalized by macrophages 2, 12, and 48 hours after injection. (**U**) Aluminum phosphate adjuvants are inside the phagocytic cells, or in the ECM. (**V**) Phagocytic cells in Lymph nodes from naïve mouse.

A Single-dose - 2 w

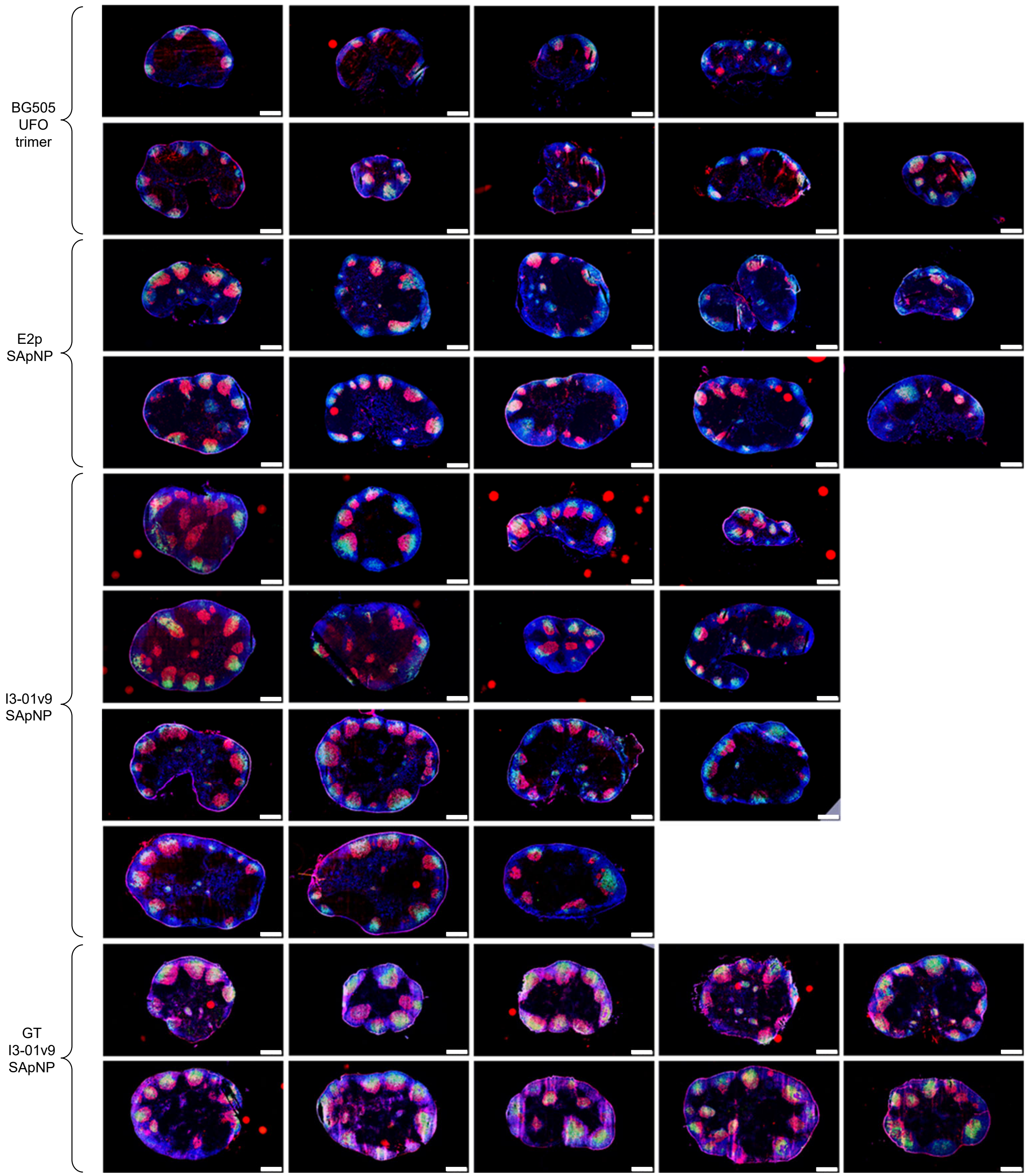

fig. S10

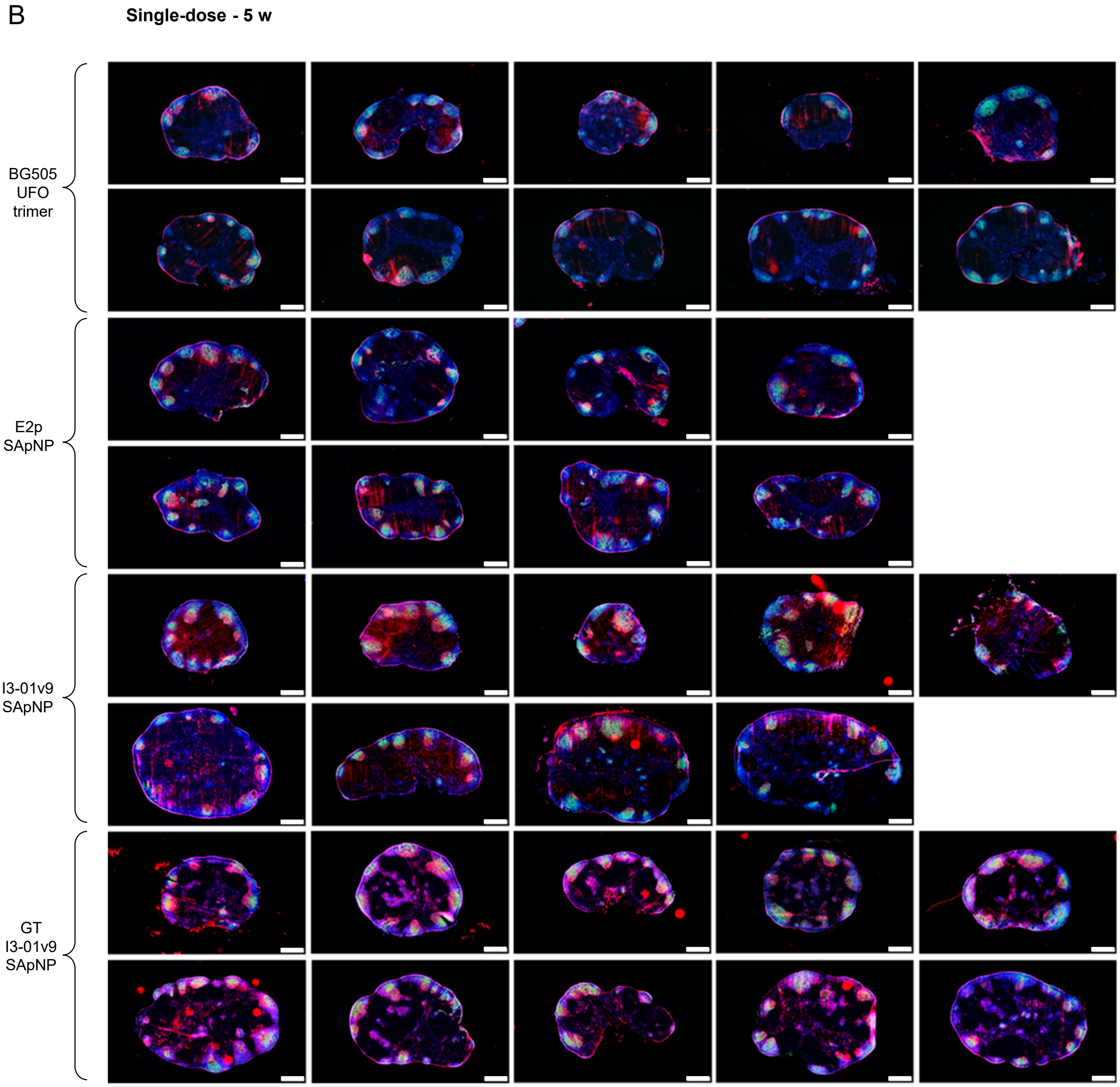
B
Single-dose - 5 w
BG505
UFO
trimer
E2p
SApNP
I3-01v9
SApNP
GT
I3-01v9
SApNP

fig. S10

C

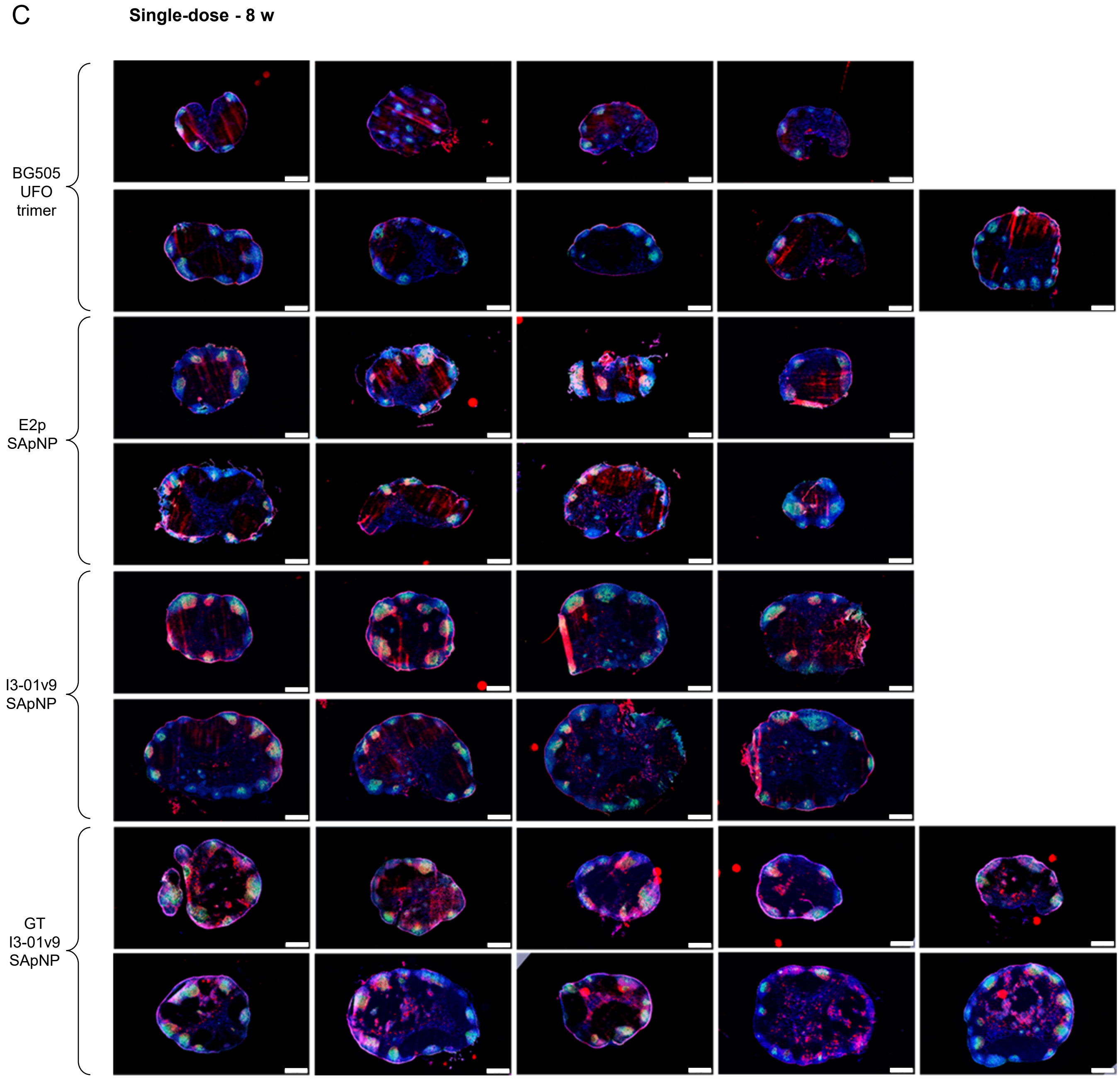

Single-dose - 8 w
BG505
UFO
trimer
E2p
SApNP
I3-01v9
SApNP
GT
I3-01v9
SApNP

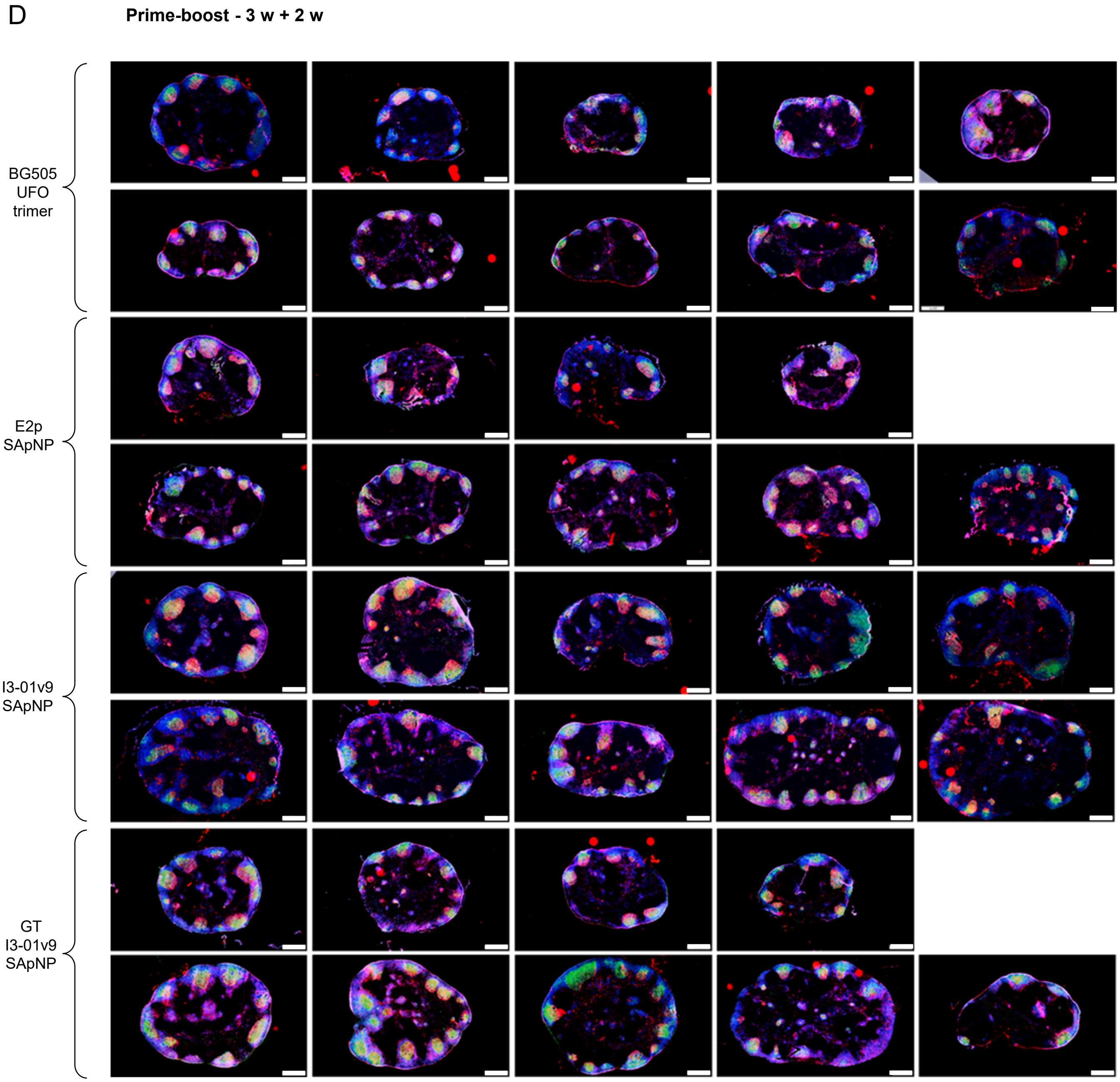
D
Prime-boost - 3 w + 2 w
BG505
UFO
trimer
E2p
SApNP
I3-01v9
SApNP
GT
I3-01v9
SApNP

fig. S10

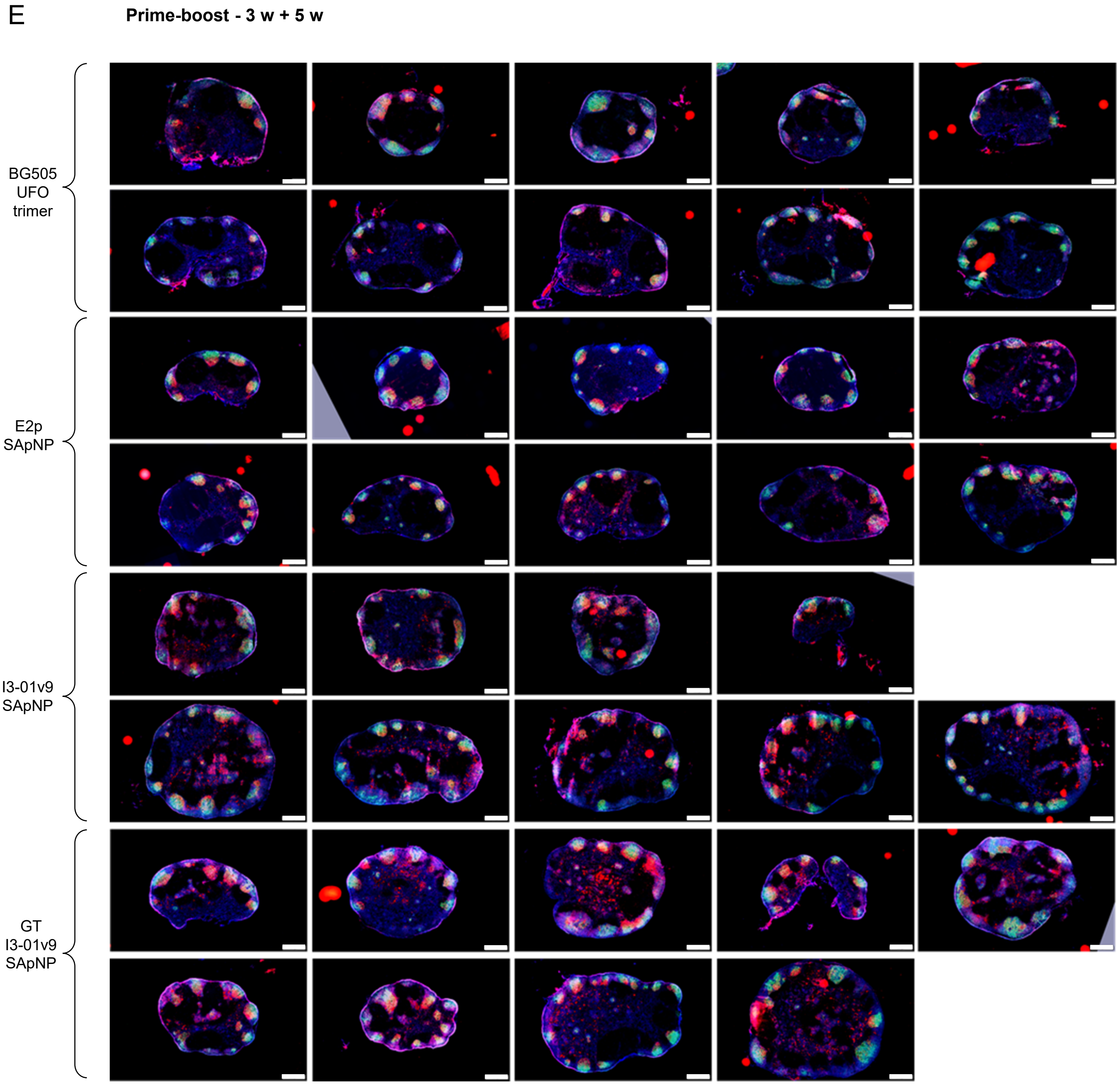


**fig. S10. Immunohistological analysis of BG505 UFO and SApNP vaccine-induced GCs.** Immunohistological images of GCs at (**A**) week 2, (**B**) week 5, and (**C**) week 8 after a single-dose injection of BG505 UFO trimer and BG505 UFO trimer-presenting E2p, I3-01v9, and GT I3-01v9 SApNP vaccines (10 μg per injection, totaling 40 μg per mouse), with a scale bar of 500 μm for each image. Images of GCs at (**D**) week 2 and (**E**) week 5 after prime-boost injections.

fig. S11

A

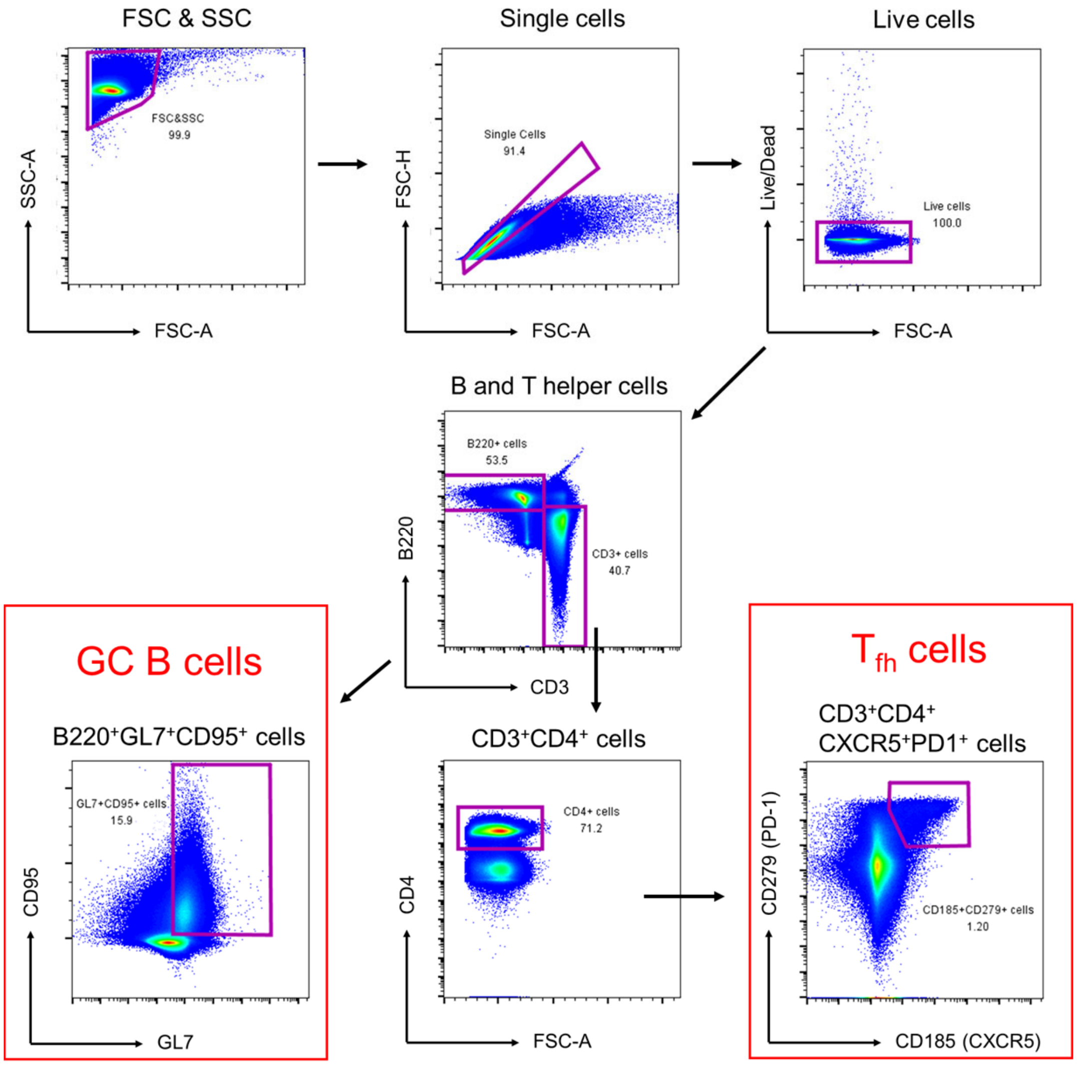

FSC & SSC
FSC&SSC
99.9
SSC-A
FSC-A
Single cells
Single Cells
91.4
FSC-H
FSC-A
Live cells
Live cells
100.0
Live/Dead
FSC-A
B and T helper cells
B220+ cells
53.5
CD3+ cells
40.7
B220
CD3
GC B cells
B220⁺GL7⁺CD95⁺ cells
GL7+CD95+ cells
15.9
CD95
GL7
CD3⁺CD4⁺ cells
CD4+ cells
71.2
CD4
FSC-A
Tfh cells
CD3⁺CD4⁺
CXCR5⁺PD1⁺ cells
CD185+CD279+ cells
1.20
CD279 (PD-1)
CD185 (CXCR5)

fig. S11

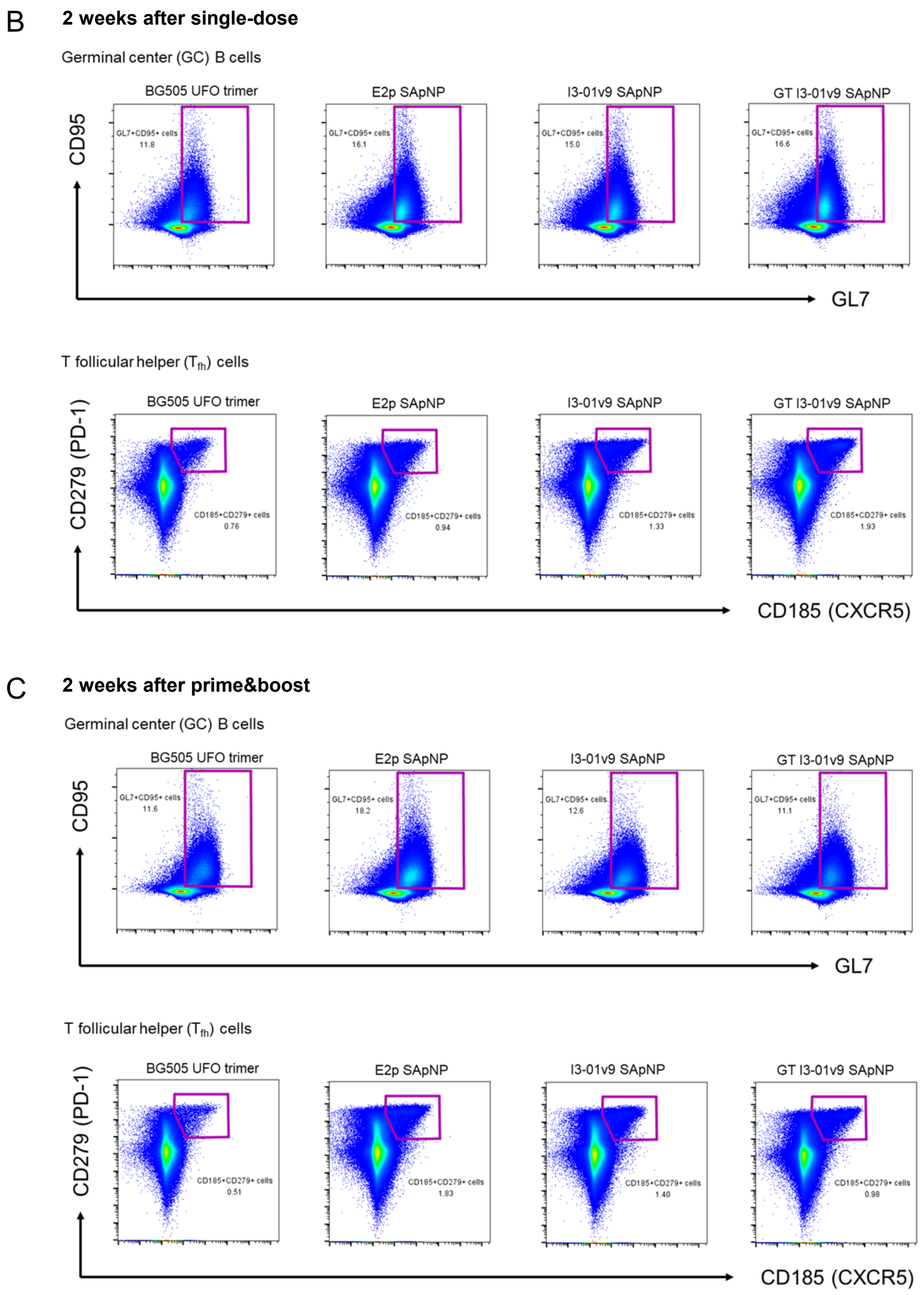


**fig. S11. Flow cytometry analysis of BG505 UFO trimer and SApNP vaccine-induced GCs.** (**A**) Gating strategy for analyzing GCs using flow cytometry. Flow plots of GC reactions (GC B cells and T follicular helper cells) at week 2 after (**B**) a single-dose (**C**) prime-boost injections of BG505 UFO trimer and BG505 UFO trimer-presenting E2p, I3-01v9, and GT I3-01v9 SApNP vaccines (10 μg per injection, totaling 40 μg per mouse).

**Table S1.** Cryo-EM data collection information.

| | BG505 UFO-10GS-I3-01v9-L7P | BG505 UFO-E2p-L4P |
|---|---|---|
| **Microscope** | Talos Arctica | Titan Krios |
| **Voltage (kV)** | 200 | 300 |
| **Detector** | Gatan K2 Summit | Gatan K2 Summit |
| **Recording mode** | Counting | Counting |
| **Magnification** | 36,000 × | 29,000 × |
| **Movie micrograph pixel size** | 1.15 | 1.03 |
| **Dose rate (e−/Å2/s)** | 4.2 | 4.9 |
| **No. of frames per movie micrograph** | 48 | 41 |
| **Frame exposure time (ms)** | 250 | 250 |
| **Movie micrograph exposure time (s)** | 12 | 10.25 |
| **Total dose (e−/Å2)** | 50.4 | 50.2 |
| **Under focus range (μm)** | 0.6 – 2.0 | 0.7 – 1.5 |
| **Number of movie micrographs** | 1200 | 2300 |

**Table S2.** Model building and refinement statistics for the BG505 UFO E2p-L4P NP core.

| | BG505 UFO-E2p-L4P core |
|---|---|
| **Map Resolution** | 3.7 |
| **EMDB ID** | - |
| **PDB** | - |
| **Residues** | 13980 |
| **Amino-acids** | 13980 |
| **Carbohydrates** | 0 |
| **RMSD Bonds** | 0.019 |
| **RMSD Angles** | 1.605 |
| **Ramachandran** | |
| **Outliers (%)** | 0.0 |
| **Allowed (%)** | 1.3 |
| **Favored (%)** | 98.7 |
| **Rotamer outliers** | 0.00 |
| **Clash score** | 0.00 |
| **Molprobity score** | 0.50 |
| **EMRinger score** | 3.51 |
